# PP2A phosphatase regulates cell-type specific cytoskeletal organization to drive dendritic diversification

**DOI:** 10.1101/2022.03.07.483285

**Authors:** Shatabdi Bhattacharjee, Erin N. Lottes, Sumit Nanda, Andre Golshir, Atit A. Patel, Giorgio A. Ascoli, Daniel N. Cox

## Abstract

Uncovering molecular mechanisms regulating dendritic diversification is essential to understanding the formation and modulation of functional neural circuitry. Transcription factors play critical roles in promoting dendritic diversity and here, we identify PP2A phosphatase function as a downstream effector of Cut-mediated transcriptional regulation of dendrite development. Mutant analyses of the PP2A catalytic subunit (*mts*) or the scaffolding subunit (*PP2A-29B)* reveal cell-type specific regulatory effects with the PP2A complex required to promote dendritic growth and branching in *Drosophila* Class IV (CIV) multidendritic (md) neurons, whereas in Class I (CI) md neurons, PP2A functions in restricting dendritic arborization. Cytoskeletal analyses reveal requirements for Mts in regulating microtubule stability/polarity and F-actin organization/dynamics. In CIV neurons, *mts* knockdown leads to reductions in dendritic localization of organelles including mitochondria and satellite Golgi outposts, while CI neurons show increased Golgi outpost trafficking along the dendritic arbor. Further, *mts* mutant neurons exhibit defects in neuronal polarity/compartmentalization. Finally, genetic interaction analyses suggest β-tubulin subunit 85D is a common PP2A target in CI and CIV neurons, while FoxO is a putative target in CI neurons.

## Introduction

Neurons must form appropriate synaptic connections to establish functional neuronal circuits which regulate cognition and behavior. These neuronal connections are reliant upon dendrites that receive, integrate, transform, and propagate synaptic and sensory inputs (Lefebvre *et al*, 2015). The importance of maintaining dendritic shape is accentuated by an array of neuropathological disorders that are associated with the aberrations in dendritic morphology as well as disruptions in the cytoskeletal architecture (Fiala *et al*, 2002; Franker & Hoogenraad, 2013; Moser, 1999). In order to ensure proper connectivity, and therefore network function, the dendritic tree must allow structural plasticity during development as well as in the mature adult neurons. Plasticity is supported by a host of intrinsic signaling molecules orchestrated by various transcription factors (TFs). These signaling cascades commonly converge on cytoskeletal components including F-actin and microtubules (MTs), that provide the underlying scaffold and fiber tracks for intracellular trafficking (Nanda *et al*, 2017; Parrish *et al*, 2007). These cytoskeletal components are organized and modulated by a wide range of proteins that regulate the assembly, disassembly, bundling, severing, stabilization, and molecular motor-based transport of vesicular cargo and organelles (Coles & Bradke, 2015; Kapitein & Hoogenraad, 2015; Nirschl *et al*, 2017).

Numerous studies in vertebrates and invertebrates have demonstrated that TFs regulate dendritogenesis (Puram & Bonni, 2013; Santiago & Bashaw, 2014; Arikkath, 2012). In *Drosophila melanogaster*, TFs act both individually and in a combinatorial fashion to give rise to subtype-specific dendritic architecture in larval multidendritic (md) sensory neurons (Grueber *et al*, 2003; Hattori *et al*, 2007; Das *et al*, 2017; Ferreira *et al*, 2014; Hattori *et al*, 2013; Iyer *et al*, 2013a, 2013b; Jan & Jan, 2010; Parrish *et al*, 2006; Sears & Broihier, 2016; Sugimura *et al*, 2004; Li *et al*, 2004; Nanda *et al*, 2017). These TFs act on diverse pathways to drive dendritic cellular diversity, many of which ultimately impact cytoskeletal architecture as terminal mediator of arbor shape (Clark *et al*, 2018; Grueber *et al*, 2003; Jinushi-Nakao *et al*, 2007; Ferreira *et al*, 2014; Iyer *et al*, 2013b; Sears & Broihier, 2016; Nanda *et al*, 2017; Das *et al*, 2021; Iyer *et al*, 2012; Nagel *et al*, 2012; Grueber *et al*, 2002; Das *et al*, 2017). However, the intermediate players in these pathways remain largely unknown as are molecular factors that drive cell-type specific dendritic diversity. Recent work has started to unravel links between combinatorial TF activity and regulation of the dendritic cytoskeleton, including the identification of novel roles for genes involved in protein homeostasis (Das *et al*, 2017; Lottes & Cox, 2020).

Protein phosphorylation is one of the most prevalent posttranslational modifications that determine the functional state of proteins (Cheng *et al*, 2011; Morrison *et al*, 2000). The phosphorylation state of a protein depends on dynamic interplay between kinases and phosphatases. While numerous kinases phosphorylate proteins at serine or threonine residues, two members of the phosphoprotein phosphatase family, Protein Phosphatase 1 (PP1) and 2A (PP2A) mainly catalyze dephosphorylation at these sites (Nasa & Kettenbach, 2018). PP2A is a highly conserved serine/threonine phosphatase required for a multitude of fundamental cellular processes (Janssens & Goris, 2001; Reynhout & Janssens, 2019). The PP2A holoenzyme is a heterotrimeric complex that consists of a catalytic (PP2AC) subunit and a scaffolding structural (PP2AA) subunit that form the core enzyme, and a variable regulatory subunit (PP2AB). In *Drosophila*, the catalytic subunit is encoded by the gene *microtubule star* (*mts*) and the scaffolding subunit by *PP2A-29B*. The regulatory subunits fall into four structurally heterogenous families denoted by B, B’, B’’ and B’’’ (Reynhout & Janssens, 2019). Compared to the 16 regulatory B subunits genes found in vertebrates, the fly genome encodes only five regulatory subunits: *twins* (B), *well-rounded* (*wrd*) (B’), *widerborst* (*wdb*) (B’), *CG4733* (B’’), and *Cka* (B’’’).

Recently published work has identified *wdb* as a downstream effector of the TF Cut (Ct) in regulating *Drosophila* md neuron dendrite development (Das *et al*, 2017). Further, two recent studies have uncovered roles of the PP2A complex in regulating dendritic pruning in CIV md neurons (Rui *et al*, 2020; Wolterhoff *et al*, 2020). Despite these advances, the mechanistic role(s) of PP2A in specifying subtype-specific dendritic architecture remains unknown. The current study is aimed at uncovering the mechanistic roles of PP2A in regulating cell-type specific dendritic arborization through modulation of the underlying cytoskeletal components. We demonstrate that mutations in PP2A subunits leads to severe reductions in dendritic complexity of CIV md neurons while increasing dendritic complexity of the simpler CI neurons. Cellularly, live imaging reveals that loss of *mts* in both CI and CIV neurons leads to MT destabilization and defects in MT polarity. In contrast to the observed effects on MTs, loss of *mts* leads to hyperstabilization of F-actin and a shift of F-actin distributions within the dendritic arbor. Loss of *mts* also results in significant reductions in the presence of organelles including mitochondria and satellite Golgi outposts along the dendritic arbor in CIV neurons. In CIV *mts* mutant conditions, Golgi outposts normally restricted to the soma and dendrites began to ectopically appear in the proximal axon. Further highlighting the difference between cell types, loss of *mts* in CI neurons results in increased Golgi outpost trafficking along the dendritic arbor. Not only do Golgi outposts appear ectopically in CIV *mts* mutants, but selective markers of dendritic and axonal compartments are also detected in the opposing compartments indicative of a defect in neuronal compartmental identity. The cell type specific defects observed in *mts* loss of function neurons may be due to differential roles of its targets. We identify regulatory interactions between PP2A and the TFs Cut and FoxO in driving CIV vs. CI cell-type specific dendritic diversity, respectively. Furthermore, phenotypic analyses reveal genetic interactions between PP2A and β-tubulin subunit 85D in both CI and CIV md neuron subtypes suggesting β-tubulin subunit 85D may represent a common target of PP2A-driven dendritic development. Collectively, these studies provide insights into the functional roles of the PP2A phosphatase in promoting dendritic diversity.

## Materials and Methods

### Drosophila husbandry and stocks

*Drosophila melanogaster* stocks were grown on standard cornmeal-molasses-agar media and maintained at 25°C. Genetic crosses were reared at 29°C. Strains used in this study are listed in Table S3.

### Generation of transgenic flies

For optimal expression, we custom synthesized a codon-bias optimized *Drosophila melanogaster β-tubulin85D* gene in which S172 and T219 were mutated to either glutamate (E) or alanine (A) (GenScript, Piscataway, NJ). Each synthesized gene was FLAG-tagged at the C-terminus and subcloned into *pUAST-attB.* Transgenic β-tubulin85D *Drosophila* strains were generated by ΦC31-mediated integration targeting 3R (*attP40)* (GenetiVision, Houston, TX). To construct *UAS-LifeAct.tdEOS* flies, we performed gene synthesis of codon-bias optimized LifeAct peptide fused to tdEOS (LifeAct.tdEOS (GeneScript)). The synthesized gene was subcloned into the *pUAST-attB* vector. Transgenic *UAS-LifeAct.tdEOS Drosophila* strains were generated by ΦC31-mediated integration targeting 3R (*attP40)* (GenetiVision).

### Immunohistochemical Analysis

Immunohistochemistry was performed as previously described (Sulkowski *et al*, 2011). Primary antibodies used were rabbit anti-PP2CB (used at 1:50 dilution) (Biorbyt); chicken anti-GFP (used at 1:1000 dilution)(Abcam); mouse anti-Cut (used at 1:100 dilution) (DSHB); mouse anti-Futsch (22C10) (used at 1:100 dilution) (Developmental Studies Hybridoma Bank); mouse anti-acetylated α-tubulin (used at 1:100) (Santa Cruz); rabbit anti-pS172 β-tubulin (used at 1:100 dilution) (ab78286, Abcam); rabbit anti-FoxO (used at 1:100 dilution) (ab195977, Abcam). Donkey anti-chicken 488 (1:1000) (Jackson Immunoresearch), donkey anti-rabbit 555 (1:200) (Life Technologies), donkey anti-mouse (1:200) (Life Technologies), and Alexa-Fluor goat anti-horseradish peroxidase (HRP) 647 (1:200) were used as secondary antibodies.

### Live Confocal Imaging, Neuronal Reconstruction, and Morphometric Analysis

Live imaging was performed as described previously (Iyer *et al*, 2013b, 2013a). Images were processed and skeletonized using ImageJ as previously described (Iyer *et al*, 2013a; Schneider *et al*, 2012). Quantitative neuromorphometric data (*e.g.* total dendritic length) were extracted and compiled using custom Python algorithms. The custom Python scripts were used to compile the output data acquired from Analyze Skeleton ImageJ plugin and the compiled output data was exported to Excel (Microsoft). For number of branches swc files were generated using NeuronStudio (Wearne *et al*, 2005) and SNT plugin on ImageJ was used. Branch density was obtained by dividing the number of branches by the total dendritic length. Dendritic field coverage was analyzed using the Internal Coverage macro for ImageJ (Sears & Broihier, 2016) (https://github.com/JamesCSears/Internal-Coverage-Macro) using a rectangular region of interest bounded by the outermost dendrite on each side, with a square size set to 20 x 20 pixel grid. Proportion covered is defined as the proportion of boxes which contain dendritic arbor against the total number of grid boxes. Sholl analysis was done using the Sholl plugin for ImageJ (Ferreira *et al*, 2014).

### Next generation multi-channel reconstructions

Multichannel cytoskeletal reconstructions and quantitative analyses were performed using a previously described method (Nanda *et al*, 2021). MT or F-actin quantity of a compartment is defined as

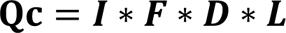

Where **Q_c_** is the cytoskeletal quantity (microtubule or F-actin quantity) of a given compartment, **I** is the relative signal intensity of the compartment, **F** is the fraction of the volume occupied by the MT or F-actin signal, **D** is the diameter of the compartment and **L** is the length of the compartment). Once quantified, the total cytoskeletal quantity of Class I and Class IV control groups (averaged across all neurons within the control groups) are normalized separately to 1 and every other mutant group within the same cell class is normalized by the same factor. Therefore, every mutant group within a neuron class is represented relative to the corresponding control group. Normalized quantity of MT or F-actin plotted against path distance from the soma are binned at 40 μm intervals.

### Denmark and Synaptotagmin analysis

Denmark signal was quantified in the same manner as MT and F-actin. The following definitions was used to measure Denmark quantity at each compartment:

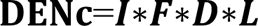

Where **DEN_c_** is the Denmark signal quantity of a given compartment, **I** is the relative signal intensity of the compartment, **F** is the fraction of the volume occupied by the Denmark signal, **D** is the diameter of the compartment and **L** is the Length of the compartment.

Synaptotagmin signal was quantified and represented by counting the number of puncta across the arbor, separately for dendrites and axons. All Synaptotagmin signals from all neurons were first measured (separately for Class IV and Class I neurons). Since the punctate expressions are not length dependent, the length of each compartment was not used (unlike the quantifications for MT, F-actin, and Denmark) to compute Synaptotagmin intensity. Instead the Synaptotagmin intensity was defined as:

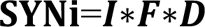

Where **SYN_I_** is the Synaptotagmin intensity of a given compartment, **I** is the relative signal intensity of the compartment, **F** is the fraction of the volume occupied by the Denmark signal and **D** is the diameter of the compartment.

Synaptotagmin intensity levels were stratified for both CI and CIV neurons, and compartments above the threshold intensity (in the top 10% of all puncta) were identified as punctate expressions of Synaptotagmin (SYN-positive). The ratio of the number of SYN-positive compartments to the total number of dendritic or axonal compartments represents the probability of finding the SYN-positive dendritic or axonal compartments, respectively.

### Tubulin and LifeAct Photoconversion Assay

To analyze MT stability/turnover, alpha tubulin 84B tagged with photoconvertible tdEOS *(UAS-alphaTUB84B.tdEOS),* was expressed under the control of *GAL^477^* in control and *mts-IR* conditions. Photoconversion experiments were conducted as previously described (Tao *et al*, 2016). Briefly, ∼30µm^2^ region of the dendrite proximal to the cell body was photoconverted from green to red by exposing to 405 nm laser for 15 seconds. The photoconverted neurons were then live imaged immediately after photoconversion (0h) and at 30 minutes intervals for up to 1 hour. Red fluorescent intensities were then measured in the photoconverted region and a neighboring non-converted region (green fluorescent) region in Zen Blue software (Zeiss). The remaining fluorescence intensities were analyzed using the following formula: (FI_converted_-FI_neuighboring_)_timecourse_/(FI_converted_-FI_neighboring_)_0h_ as previously described (Tao *et al*, 2016).

To analyze F-actin turnover, fly lines that expressed LifeAct tagged with photoconvertible tdEOS *(UAS-LifeAct.tdEOS)* were crossed to either *GAL4^ppk1.9^* or *mts-IR; GAL4^ppk1.9^* flies. Photoconversion experiments were performed as described above.

### Cell organelle imaging and analysis

Live imaging was performed on wandering third instar larvae as described previously (Iyer *et al*, 2013a, 2013b). Images were acquired on a Zeiss LSM 780 confocal microscope as z-stacks using 20x (dry) or 63x (oil immersion) objective at step size 1-1.5µm and 1024 x 1024 resolution. Maximum intensity projections of the z-stacks were then exported as .jpeg or TIFF files using the Zen Blue software. Using Adobe Photoshop, images were cropped to a fixed size (6.6 in x 4.5 in) in the same quadrant of the neurons and Golgi outposts, mitochondria, and γ-tubulin-GFP were manually counted along the dendrites in these quadrants. The cropped images were also processed to obtain the total dendritic length as described above.

### EB1 comet imaging and analysis

Time-lapse imaging of EB1::GFP comets in both control and *mts-IR* animals were acquired using 40x oil objective at a digital zoom of 2.5X. Comet movies were recorded for 100 seconds at 30 frames per second. The number of comets moving in either anterograde or retrograde directions were manually scored. The comet speeds were analyzed using Imaris software (Bitplane).

### qRT-PCR

qRT-PCR analysis was done between control and *ct-IR* expressing neurons in quadruplicates as previously described (Iyer *et al*, 2013b). Briefly, CIV neurons expressing *UAS-mCD8::GFP* were isolated using superparamagnetic beads (Dynabeads MyOne Steptavidin T1, Invitrogen) that were conjugated to biotinylated anti-mCD8a antibody (eBioscience). RNA was isolated from these CIV neurons using the miRCURY RNA Isolation Kit (Exiqon) and qRT-PCR analysis was performed using pre-validated Qiagen QuantiTect Primer Assays using *mts* (QT00502460). Expression data were normalized to *GAPDH2* (QT00922957) and reported as fold change in expression.

### Statistical analysis and data availability

Statistical analysis and data plotting were done using GraphPad Prism 8. Error bars in the study represent standard error of mean (SEM). Statistical tests performed were unpaired t-test; one-way ANOVA using Dunnett’s or Sidak’s multiple comparison test; two-way ANOVA with Dunnett’s multiple comparison test; Mann-Whitney U test; Kruskal Wallis using Dunn’s multiple. All the data were tested for normality using the Shapiro-Wilk normality test. Significant scores indicated on the graphs are (* = p ≤ 0.05, ** = p≤ 0.01, *** = p≤ 0.001). All new genotypes reported in the study are available upon request. Neuronal reconstructions have been submitted to NeuroMorpho.Org (Akram et al, 2018).

## Results

### PP2A regulates cell type specific dendritic architecture

A recent study implicated the PP2A B’ regulatory subunit *widerborst* (*wdb*) in *Drosophila* md neuron dendrite development (Das *et al*, 2017), while two other studies reveal roles of the PP2A complex in regulating dendritic pruning in CIV md sensory neurons (Rui *et al*, 2020; Wolterhoff *et al*, 2020) and an additional study identified PP2A components in regulating CIV larval md neuron dendritogenesis (Wang et al., 2020), however the putative mechanistic role(s) of PP2A in regulating subtype-specific dendritic arborization during larval development remain largely unknown. To investigate the potential functional requirements of PP2A in this process, we first conducted IHC analyses which revealed that the catalytic subunit, Mts, is expressed in md neuron subclasses (**Fig S1A,A’**). Given this expression, we next conducted *in vivo* neurogenetic analyses of loss-of-function mutants using gene specific *UAS-RNAi* (IR) lines that were expressed in CI and CIV md neurons by tissue specific GAL4 lines in order to investigate the potential role of PP2A in regulating cell type specific dendritic architecture. These RNAi knockdown analyses were further corroborated using MARCM mutant analyses. Combined, these studies revealed subtype-specific requirements of PP2A in regulating dendritic morphology.

In morphologically complex CIV neurons, cell-type specific knockdown of the catalytic (*mts*) and the scaffolding (*PP2A-29B*) subunits led to severely reduced dendritic arborization (**Fig 1 A, B, D**). Quantitative analysis revealed that compared to control, *mts* and *PP2A-29B* knockdown led to reductions in total dendritic length and total number of branches (**Fig 1 E, F**). Consistent with these phenotypic defects, we likewise observed that knockdown of the catalytic or the scaffolding subunits in CIV neurons reduced dendritic field coverage (**Fig 1 G**). Sholl analysis was done to determine the distribution of branching as a function of distance from the cell body (Sholl, 1953). Knockdown of *mts* or *PP2A-29B* showed a significant reduction in branching along the dendritic arbor (**Fig 1 H**). In addition, *mts* knockdown showed a significant reduction in the maximum number of Sholl intersections and further displayed a proximal shift in the radius corresponding to the maximum intersections **(Fig S1 B, C)**. *PP2A-29B* knockdown significantly reduced the maximum number of intersections **(Fig S1 B)** but the corresponding radius remained unaffected **(Fig S1 C)**. In controls, Strahler branch order extended from 7^th^ order, representing the primary dendritic branches closest to the soma, to 1^st^ order, representing terminal branches which account for the majority of branches in these neurons (**Fig 1 J, K**). In contrast, *mts* knockdown resulted in CIV neurons that only extend up to the 5^th^ branch order, coupled with significant reductions in number of branches in all Strahler orders from 1^st^ to 5^th^. Similarly, in *PP2A-29B-IR* neurons, branches extended up to the 6^th^ Strahler order with significant reduction in branches in the 1^st^ to 5^th^ branch orders relative to controls (**Fig 1 J, K**).

**Figure 1:**
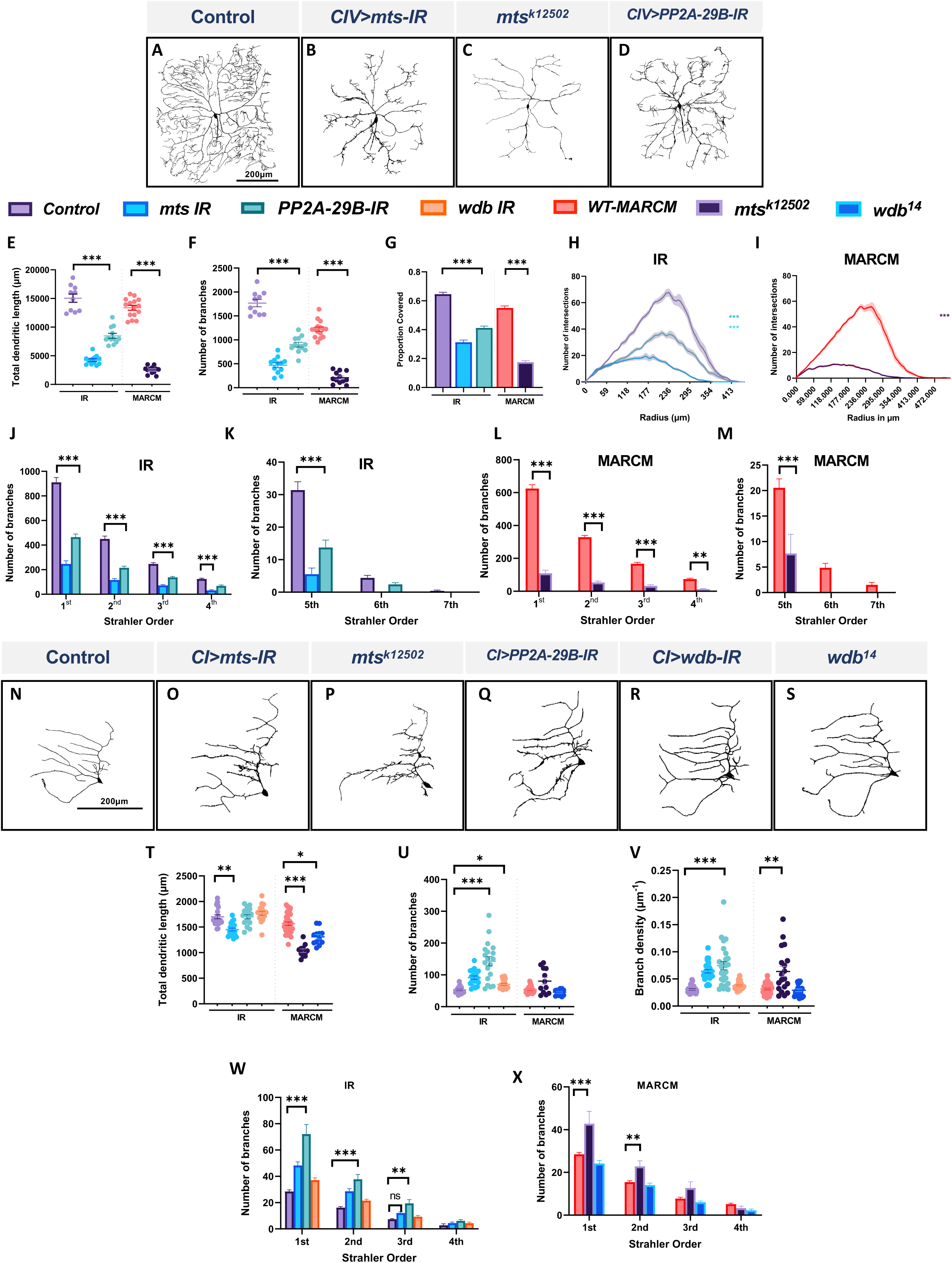
PP2A regulates cell-type specific dendritic arborization in CI and CIV md neurons. **(A-D**) Representative images of dendritic arbors of CIV (ddaC) neurons in **(A)** controls, **(B)** *mts-IR* **(C)** *mts^k12502^* MARCM clone, **(D)** *PP2A-29B-IR.* **(E-G)** Quantitative morphometric analyses. **(H, I)** Sholl profiles of control vs *mts-IR* or *PP2A-29B-IR* **(H)**, and *mts^k12502^* MARCM **(I)**. **(J-M)** Strahler analysis of control, *mts-IR, PP2A-29B-IR* **(J, K)** and *mts^k12502^* MARCM **(L, M). (N-S)** Representative images of dendritic arbors of CI (ddaE) neurons in **(N)** controls, **(O)** *mts-IR* **(P)** *mts^k12502^* MARCM, **(Q)** *PP2A-29B-IR,* **(R)** *wdb-IR,* and (**S**) *wdb^14^* MARCM. **(T-V)** Quantitative morphometric analyses. **(W, X)** Strahler analysis of control vs *mts-IR, PP2A-29B-IR,* or *wdb-IR* **(W)** and *mts^k12502^* or *wdb^14^* MARCM **(X).** Statistical tests performed: **(E-G)** One-way ANOVA with Sidak’s multiple comparisons, **(H, I)** Kruskal Wallis with Dunn’s multiple comparison, **(J-M)** Two-way ANOVA with Dunnett’s test for multiple comparison, **(T-V)** One-way ANOVA with Sidak’s multiple comparisons or Kruskal Wallis with Dunn’s multiple comparison, **(W, X)** Two-way ANOVA with Dunnett’s test for multiple comparison. ***=p≤0.001, *=p≤0.05. For **E-L**, n = 10-15 per genotype. For **S-X**, n = 12-34 per genotype. For detailed genotypes see Table S1 and for detailed statistics see Table S2. Quantitative data is reported as mean ± SEM. For Sholl analysis, values are mean ± SEM for the number of intersections as a function of radial distance from the cell body (zero). For Strahler analysis, values are mean ± SEM for the number of dendritic branches in each branch order (Strahler Order), for CIV neurons, 7^th^ is the primary branch closest to the soma and 1^st^ is the terminal branch. For CI neurons, 4^th^ is the primary branch closest to the soma and 1^st^ is the terminal branch. Scale bar = 200 µm.

We have previously shown that the PP2A regulatory subunit *wdb* is required to promote dendritic arborization in CIV md neurons (Das *et al*, 2017). Consistent with these results, *wdb* knockdown led to reductions in the total dendritic length, and number of branches in CIV neurons **(Fig S1 D, E, I, J)**. To determine if other B regulatory subunits may play in CIV larval dendrite development, we next examined potential roles of the other three regulatory subunits in CIV neuron morphology. Compared to *wdb*, knockdown of other regulatory B subunits including *twins* (*tws*), *well-rounded* (*wrd*), and *CG4733* in larval CIV neurons did not have as severe an effect on dendritic morphology. While the knockdown of *tws* and *wrd* led to a mild but significant reduction in the total dendritic length, dendritic branches remained unaffected. Knockdown of *CG4733* did not significantly affect either parameter when compared to controls **(Fig S1 D-J)**. A recently published study implicated both Wdb and Tws in regulating dendritic pruning with the effects of Tws knockdown being milder than that of Wdb (Wolterhoff *et al*, 2020). This is consistent with what we observed in larval CIV neurons, however, our results also suggest a putative role for Wrd in promoting dendritic growth.

To independently assess the roles of PP2A subunits on CIV dendrite morphogenesis, we conducted Mosaic Analysis with a Repressible Cell Marker (MARCM) studies examining cell autonomous requirements of *mts* (Lee & Luo, 1999). Consistent with the results obtained from RNAi analysis, MARCM analyses revealed that *mts* mutant neurons (*mts^k12502^*) exhibit severely impaired CIV dendritic arborization (**Fig 1 C**) with reductions in all the morphometric parameters tested (**Fig 1 E-G**).

Sholl analysis identified a significant reduction in dendritic complexity as a function of distance from the soma (**Fig 1 I**). There was a significant decrease in the maximum number of intersections combined with a shift in dendritic complexity towards the soma **(Fig S1 B, C)**. Similar to *mts-IR*, *mts^k12502^* MARCM clones showed a total loss of 6^th^ and 7^th^ order branches (**Fig 1 L, M**). These phenotypic defects are consistent with previous reports (Rui *et al*, 2020; Wolterhoff *et al*, 2020).

Given that we observed Mts expression in other md neuron subtypes, we sought to test how the PP2A complex may play a role promoting cell type specific dendritic diversity by focusing on the morphologically simpler CI md neurons. In contrast to mutant effects on CIV neurons, disruption of *mts* or *PP2A-29B* led to increased dendritic growth and branching as revealed by an increase in *de novo* short ectopic branching (**Fig 1 N, O, Q**). Quantitative analysis showed an increase in number of branches in *mts-IR* and *PP2A-29B-IR* animals compared to controls (**Fig 1 U**). Branch density, which was obtained by normalizing number of branches to the total dendritic length, also showed a significant increase in *mts-IR* and *PP2A-29B-IR* animals compared to controls (**Fig 1 V**). Strahler order analysis showed that there was significant increase in the 1^st^ and 2^nd^ order branches for both *mts-IR* and *PP2A-29B-IR* animals and 3^rd^ order branches for *PP2A-29B-IR* animals (**Fig 1 W**). *mts* knockdown also led to a reduction in total dendritic length, suggesting that the increase in branching was mostly due to short ectopic branching (**Fig 1 T**). Knockdown of *PP2A-29B* did not lead to a decrease in total dendritic length (**Fig 1 T**). Knockdown of *wdb* produced a modest increase in number of branches (**Fig 1 R, U**) while total dendritic length, and branch density remained unchanged (**Fig 1 T, V**). Strahler order analysis did not reveal any significant change in branch orders between control and *wdb-IR* (**Fig 1 W**). Consistent with the RNAi analysis, *mts^k12502^* MARCM mutant clones showed an increase in branch density, however, number of branches was not statistically different (**Fig 1 P, T-V**). Strahler analysis showed a significant increase in 1^st^ and 2^nd^ order branches, similar to that observed with *mts-IR* (**Fig 1 X**). Like *mts-IR, mts^k12502^* led to a significant decrease in total dendritic length (**Fig 1 T**). Cell autonomous analysis of *wdb^14^* mutants in CI neurons revealed a mild reduction in total dendritic length, while other neuromorphometric parameters were unchanged (**Fig 1 S, T-V, X**). Similar to knockdown in CIV, knockdown of the *tws, wrd* or *CG4733* B regulatory subunits did not dramatically impact dendritic morphology with only knockdown of *CG4733* leading to a modest reduction in the total dendritic length while the number of branches were unaffected **(Fig S1 K-P).** Collectively, these findings suggest that the PP2A holoenzyme complex exerts cell type specific regulatory effects that contribute to dendritic diversity by restricting dendritic branching in CI neurons while promoting growth and branching in CIV neurons.

To determine if homeostatic regulation of PP2A is required for cell-type specific dendritogenesis, we conducted overexpression studies for *mts*, *PP2A-29B* and *wdb* in CIV md neurons. Overexpression of *mts* or *wdb* severely disrupted dendritic morphology with reductions in total dendritic length, number of branches, and field coverage (**Fig 2 A, B, D, E-G**). Qualitatively, overexpression of *PP2A-29B* seemed to have a relatively moderate effect on dendritic morphology in comparison to that observed due to the overexpression of *mts* or *wdb* (**Fig 2 C, E-G**). *PP2A-29B-OE* led to a decrease in total dendritic length, number of branches, and field coverage compared to controls (**Fig 2 C, E-G**). Collectively, these data suggest that homeostatic regulation of PP2A component is required for normal CIV dendrite morphogenesis.

**Figure 2:**
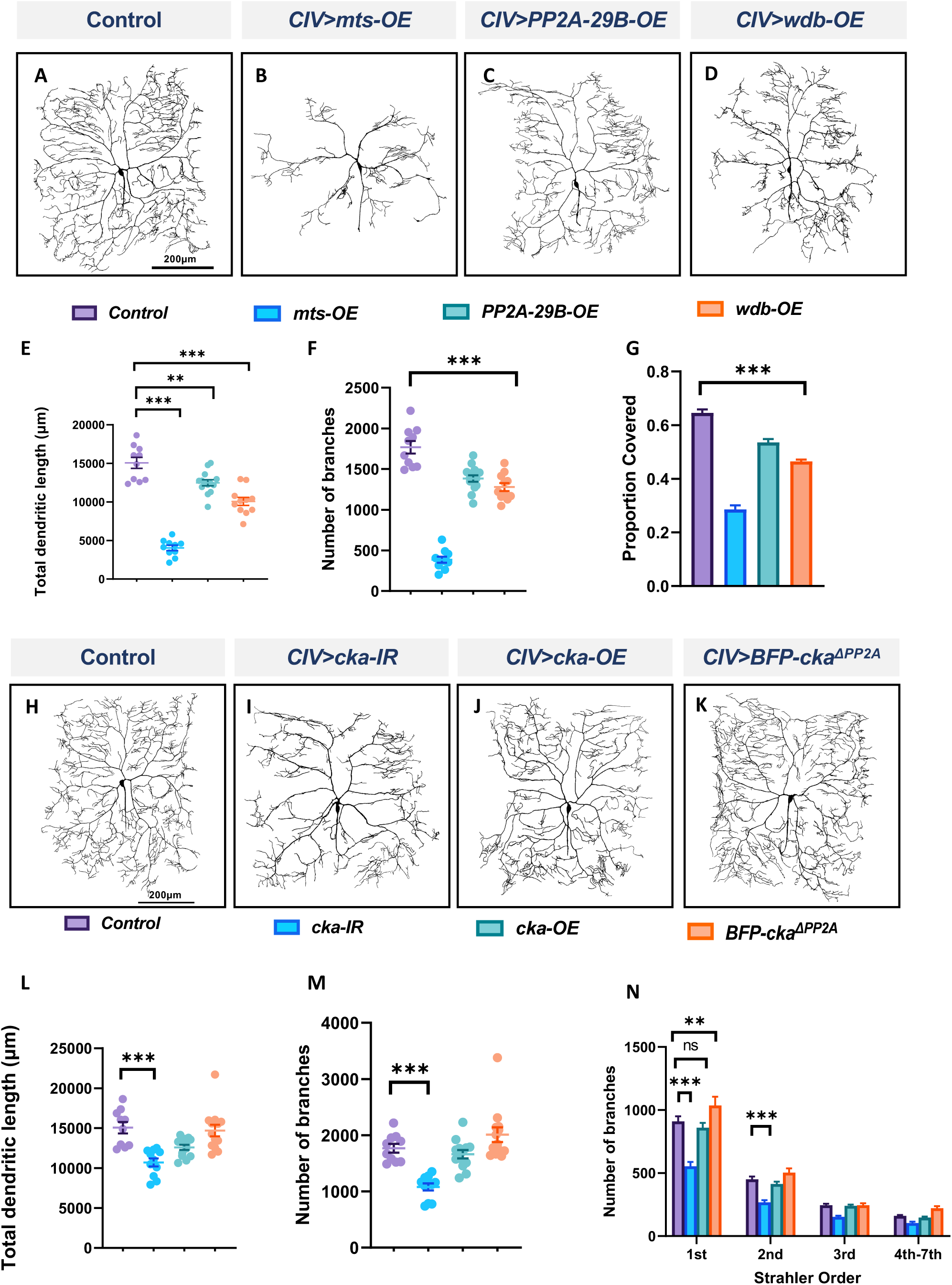
PP2A overexpression impairs dendritic morphogenesis and the PP2A and STRIPAK complexes act in parallel to regulate dendritic complexity in CIV md neurons: **(A-D)** Representative images of dendritic arbors of CIV (ddaC) neurons in **(A)** controls, **(B)** *mts-OE* **(C)** *PP2A-29B-OE* **(D)** *wdb-OE*. OE=overexpression. **(E-G)** Quantitative morphometric analyses. **(H-K)** Representative images of dendritic arbors of CIV (ddaC) neurons in **(H)** controls, **(I)** *cka-IR* **(J)** *cka-OE*, **(K)** *BFP-Cka^ΔPP2A^*, **(L,M)** Quantitative morphometric analyses. **(N)** Strahler analysis of controls, *cka-IR, cka-OE,* and *BFP-Cka^ΔPP2A^*. Statistical tests performed: **(E-G)** One-way ANOVA with Dunnett’s multiple comparison test (n=10-14 per genotype); **(L, M)** Kruskal Wallis with Dunn’s multiple comparison; **(N)** Two-way ANOVA with Dunnett’s test for multiple comparison (n=10-13 per genotype), ***=p≤0.001, **=p≤0.01, *=p≤0.05. For detailed genotypes see Table S1 and for detailed statistics see Table S2. Quantitative data is reported as mean ± SEM. For Strahler analysis, values are mean ± SEM for the number of dendritic branches in each branch order (Strahler Order) here 7^th^ is the primary branch closest to soma and 1^st^ is the terminal branch. (Scale bars = 200 µm).

### PP2A functions independently of the STRIPAK complex to regulate dendritic morphology

Striatin-interacting phosphatase and kinase (STRIPAK) is a highly conserved protein complex implicated in numerous cellular processes (Neisch *et al*, 2017; Hwang & Pallas, 2014). In *Drosophila,* the core complex consists of Mts, PP2A-29B, Connector of Kinase to AP-1 (Cka) (regulatory subunit of the complex), Mob4 (the ortholog of the mammalian striatin interactor Mob3), Ccm3, STRIP (the ortholog of mammalian STRIP1), and the kinase germinal center kinase III (GCKIII) (Kean *et al*, 2011). Due to the inclusion of PP2A components Mts and PP2A-29B in the STRIPAK complex, we sought to test whether PP2A-mediated effects on md neuron dendritic development may occur in conjunction with STRIPAK function. Disruption of *Cka* in CIV neurons led to reduction in dendritic complexity with decreases in total dendritic length, and number of branches compared to control (**Fig 2 H, I, L, M**). Strahler order analysis showed a significant reduction in 1^st^ and 2nd branches in *Cka-IR* neurons relative to control (**Fig 2 N**), however no other branch orders were affected (**Fig 2 N**). By contrast, *Cka* overexpression in CIV neurons did not affect measured dendritic parameters compared to controls (**Fig 2 J, L-N**). To determine if the *Cka-IR* phenotypic defects were due to its association with PP2A, we expressed a mutant form of Cka in CIV neurons bearing mutations in conserved residues required for Cka-PP2A binding (Neisch *et al*, 2017). Expressing the mutant *Cka^ΔPP2A^* form did not affect the dendritic morphology in CIV neurons in any of the morphometric parameters tested (**Fig 2 K-N**), These results suggest that PP2A is not operating through the STRIPAK complex to regulate CIV dendritic morphology, but rather that the STRIPAK complex may operate in parallel with PP2A to regulate dendrite development.

### PP2A is required for dendrite growth in late larval development

To assess the requirements of PP2A for dendritic arborization at different stages of larval development, we conducted developmental time course studies, analyzing control and *mts* knockdown CIV neurons at 24, 48, 72, and 96 hours after egg lay (AEL). To address the potential effects of maternal perdurance, we studied *mts-IR* disruption in both germline and CIV neurons using both *nanos-GAL4* (germline) and *ppk-GAL4* (CIV) to drive expression of *mts-IR* (Das *et al*, 2021). No phenotypic differences were observed between controls and *mts-IR* at 24h AEL (**Fig 3 A, B, C-E**). At 48h AEL, only internal field coverage was significantly reduced from controls while the other parameters remained unaffected (**Fig 3 A’, B’, C-E**). At 72h AEL, there is a significant reduction in all the parameters tested (**Fig 3 A”, B”, C-E**) which is further exacerbated at 96h AEL (**Fig 3** **A’’’, B’’’, C-E)**. Given that control CIV neuron dendrites exhibit space-filling growth properties over developmental time, while *mts-IR* CIV dendrites exhibit a progressive dendritic hypotrophy over time, these results suggest *mts* mutant neurons have a slowed growth phenotype that manifests in later stages of larval development.

**Figure 3:**
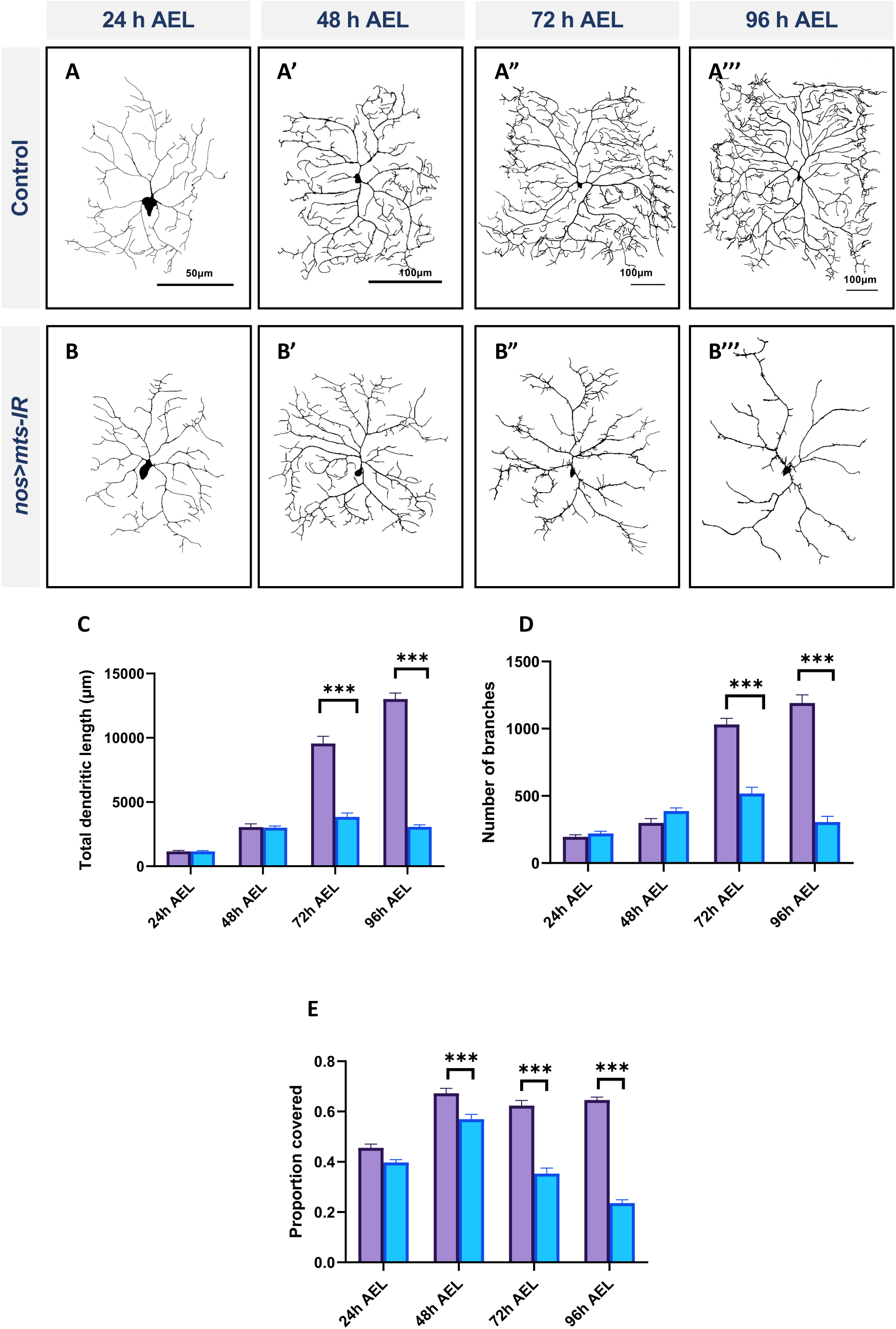
PP2A is required for dendrite growth in late larval development: Representative images of CIV md neurons of **(A-A””)** controls and **(B-B””)** *mts-IR* at 24h, 48h, 72h, and 96h after egg laying (AEL), respectively. **(C-E)** Quantitative morphometric analyses. Statistical tests performed: Two-way ANOVA with Sidak’s multiple comparisons (n = 9-20 per genotype) ***=p≤0.001. For detailed genotypes see Table S1 and for detailed statistics see Table S2. Quantitative data is reported as mean ± SEM. Scale bar = 50 µm for **A,** Scale bar = 100 µm for **A’ – A’’’.**

### PP2A acts downstream of the TF Cut to regulate dendritic morphology in CIV neurons

The transcription factor Cut (Ct) is known to regulate cell type specific dendritic morphology in md neurons in an expression level dependent manner (Grueber *et al*, 2003). We previously demonstrated that Ct positively regulates *wdb* expression and that *wdb* functions as a downstream effector of Ct-mediated dendritic development (Das *et al*, 2017), and thus we sought to determine if other components of the PP2A holoenzyme complex may also be subject to Ct regulation to promote cell type specific dendritic development. To this end, we conducted qRT-PCR analysis to measure *mts* expression levels in CIV neurons expressing *ct-IR.* Knockdown of *ct* led to a significant decrease in *mts* levels (**Fig 4** **D)** indicative of a positive regulatory relationship. To determine the potential significance of the transcriptional regulation of Mts by Ct, we knocked down *ct* in CIV neurons while simultaneously overexpressing *mts*. We hypothesized that if Ct acts through Mts to regulate dendritic morphology in these neurons, overexpression of *mts* in a *ct-IR* background may rescue some of the phenotypic defects observed due to the *ct* disruption. Knockdown of *ct* in CIV neurons leads to a dramatic decrease in dendritic complexity with decreases in total dendritic length, and number of branches (**Fig 4 A, B, E, F**). Strahler order analysis shows that in *ct-IR* neurons, dendrites extend only up to the 6^th^ order. Moreover, there was significant reduction in branches in 1^st^ to 3^rd^ orders as well as 5^th^ order in *ct-IR* compared to controls (**Fig 4 G, H**). Overexpression of *mts* in the *ct-IR* background led to a partial rescue in the phenotypic defects due to the knockdown of *ct* (**Fig 4 C**) as demonstrated by an increase in total dendritic length, and number of branches (**Fig 4 E, F**) relative to *ct-IR* alone. While there was no significant difference between the 2^nd^ to 7^th^ order branches in *ct-IR* vs. *mts-OE/ct-IR* animals, there was a significant recovery of the 1^st^ order branches in *mts-OE/cut-IR* animals compared to *ct-IR* (**Fig 4 G, H**). These data indicate that Mts acts downstream of Ct to promote proper dendritic morphology in CIV md neurons.

**Figure 4:**
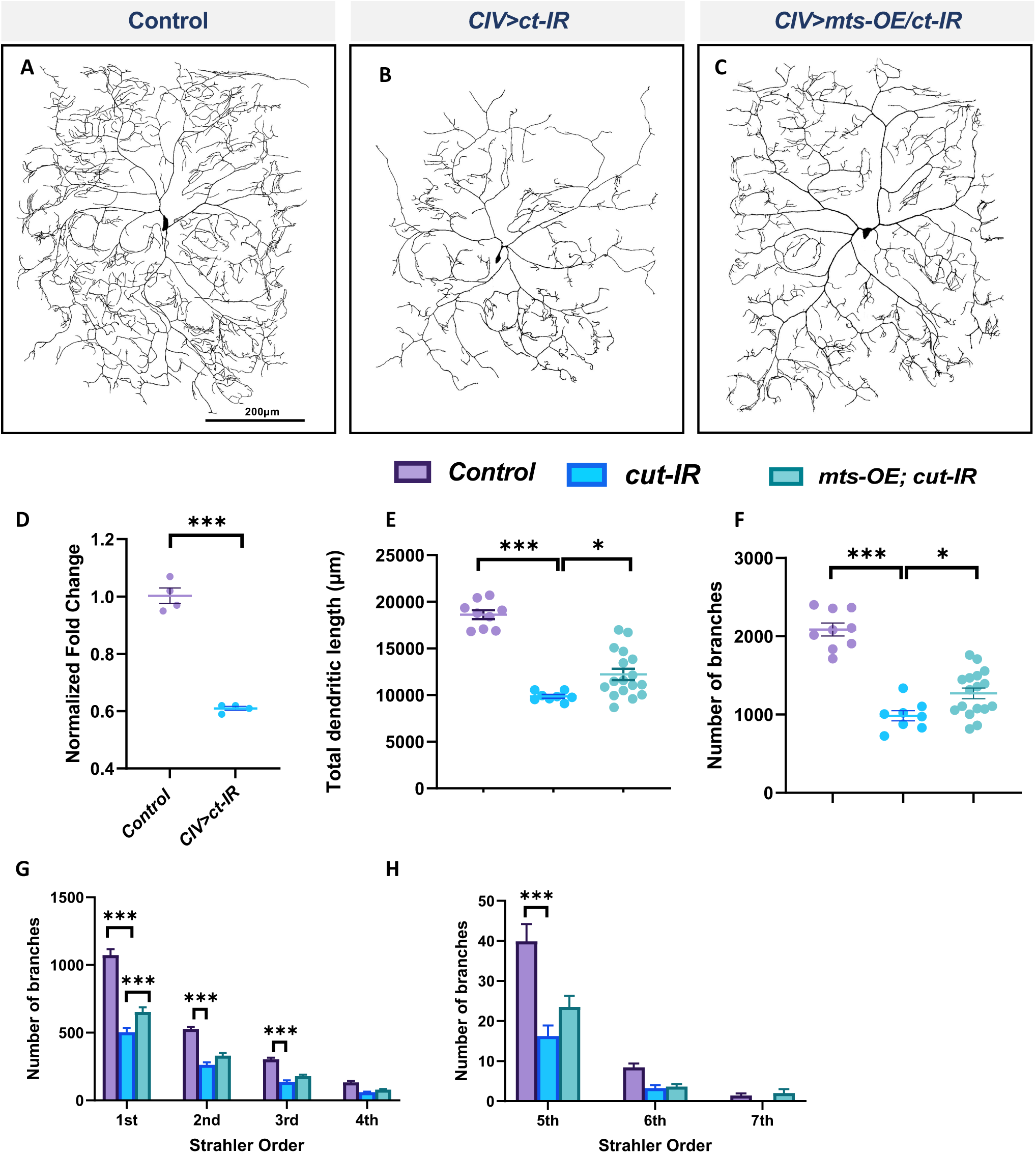
PP2A is required for Cut-mediated dendritic arborization. (**A-C**) Representative images of **(A)** control, **(B)** *ct-IR,* **(C)** *ct-IR/mts-OE* of CIV ddaC md neurons. **(D)** qPCR analyses of *mts* expression levels in control and *ct-IR* CIV neurons. **(E, F)** Quantitative morphometric analyses. **(G, H)** Strahler analysis of controls, *ct-IR,* and *ct-IR; mts-OE*. Statistical tests performed: **(D)** Unpaired *t-test* (n = 4); **(E,F)** One-way ANOVA with Sidak’s multiple comparison); **(G, H)** Two-way ANOVA with Tukey’s test for multiple comparison (n = 9-17 per genotype), ***=p≤0.001, *=p≤0.05. For detailed genotypes see Table S1 and for detailed statistics see Table S2. Quantitative data is reported as mean ± SEM. For Reverse Strahler analysis, values are mean ± SEM for the number of dendritic branches in each branch order (Strahler Order) here 7^th^ is the primary branch closest to soma and 1^st^ is the terminal branch. Scale bar = 200 µm.

### PP2A regulates microtubule and F-actin based cytoskeletal dendritic architecture

Cut is known to affect dendritic morphology by regulating the cytoskeleton (Jinushi-Nakao *et al*, 2007; Das *et al*, 2017). Since PP2A subunits including Mts (**Fig 4**) and Wdb (Das *et al*, 2017) function as downstream effectors of Cut, and our previous studies implicated Wdb in regulating dendritic cytoskeletal architecture (Das *et al*, 2017), we hypothesized that disrupting the catalytic function of the PP2A holoenzyme would impact the dendritic cytoskeleton. To study cytoskeletal dendritic architecture, we used multi-fluorescent *Drosophila* transgenic lines to simultaneously visualize the F-actin (*UAS-GMA*) and MT (*UAS-mCherry::Jupiter*) cytoskeletons in a subtype-specific manner for both control neurons and those with gene-specific mutations (Das *et al*, 2017). We also built next generation neuroanatomical reconstruction tools for quantitative descriptions of the dendritic effects of gene-specific disruptions on cytoskeletal architecture (Das *et al*, 2017; Nanda *et al*, 2018, 2020).

In CIV neurons, *mts* knockdown led to severe reductions in dendritic stable MTs together with a proximal shift in peak F-actin signal (**Fig 5 A-D’, E-J**). With respect to MTs, *mts-IR* neurons exhibit reductions in MT quantity across most of the dendritic arbor (**Fig. 5 E**). Normalized to total dendritic length, we observed significant reductions in total MT quantity in the 1^st^-to-5^th^ Strahler orders (**Fig. 5 F, G**). With respect to F-actin, the proximal shift in distribution was accompanied by a significant reduction in the total F-actin quantity (**Fig 5 H, J**) indicative of an overall reduction in F-actin levels. However, when F-actin quantity for each *mts* mutant Strahler order was normalized to the corresponding dendritic length for that branch order, we observed significant increases in F-actin in 1^st^ to 4^th^ orders and a notable increase in local F-actin quantity in the 6^th^ order branches near the soma (**Fig 5 K**). This finding is further supported by the peak level of F-actin which occurs more proximal to the soma in *mts-IR* neurons relative to controls (**Fig. 5 I**). The overall reduction in F-actin levels could thus be attributed to the reduction in branching.

**Figure 5:**
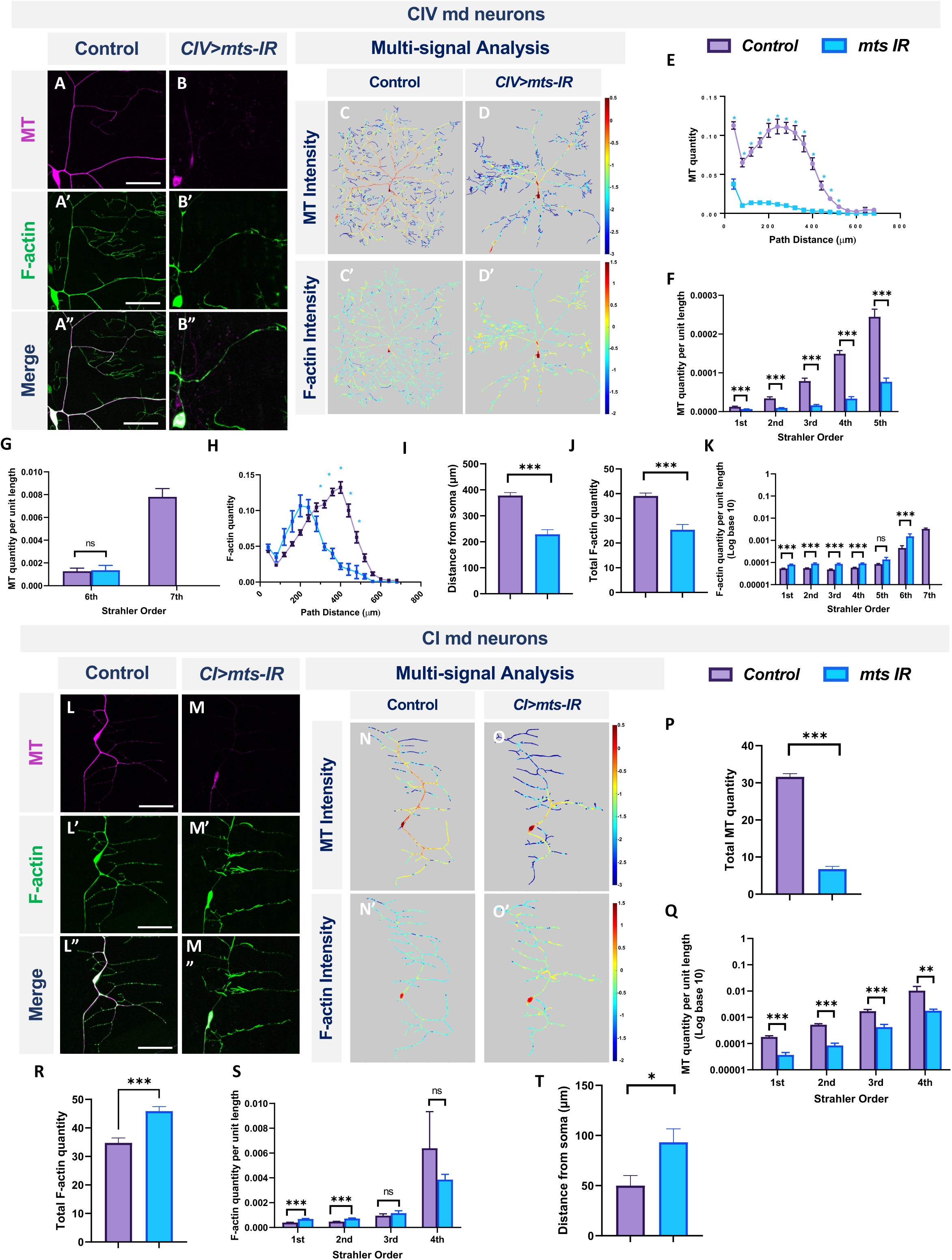
PP2A regulates MT stability and F-actin organization in md neurons: Representative images of CIV md neurons in **(A-A”)** control and **(B-B”)** *mts-IR* expressing *UAS-GMA* (labeling F-actin) and *UAS-mCherry::JUPITER* (labeling MTs). **(C-D’, M-N’)** Intensity heatmaps of MT and F-actin distributions in the dendritic arbor generated by a combination of Vaa3D multi-signal plugin, Neutube, and TREES Toolbox (MATLAB). **(E, H)** Relative subcellular distribution of MTs and F-actin along the dendritic arbor as a function of path distance from the soma. **(F, G, K, Q, S)** MT and F-actin quantities by Strahler order distribution normalized to dendritic length. **(J, P, R)** Total MT **(P)** and F-actin **(J, R)** quantities normalized to 1 for either Class I or Class IV control neurons. **(I, T)** Peak F-actin intensity as function of distance from soma. Representative images of CI md neurons in **(L-L”)** control and **(M-M”)** *mts-IR* expressing *UAS-GMA* and *UAS-mCherry::JUPITER*. Statistical tests performed: **(E, F, G, H, K, Q, S)** Multiple t-test or Mann-Whitney test with False Discovery Rate correction; **(I, J, P, R, T)** Unpaired *t-test* or Mann-Whitney U test. ***=p≤0.001, *=p≤0.05 (For **E** and **G**, *=p≤0.001). For detailed genotypes see Table S1 and for detailed statistics see Table S2. Scale bar = 20 µm for **(A-A”),** and scale bar = 50 µm for **(L-L”).** For **(E-K)** n = 11, and for **(P-T)** n = 8-9 per genotype.

Similar to findings in CIV neurons, *mts* knockdown in CI neurons led to reductions in stable MT quantities (**Fig 5 L-M, N, O, P, Q**). However, the F-actin quantities increased for 1^st^ and 2^nd^ branch orders while remaining unaffected for 3^rd^ and 4^th^ order branches (**Fig 5S**). In addition, the maximal F-actin intensities shifted distally towards the dendritic terminals with *mts* knockdown (**Fig 5 L’-M”, N’-O’, S, T**). In controls, the peak F-actin intensities correspond to ∼40μm from the soma, however, in *mts-IR* neurons, this peak intensity shifts distally to ∼100μm from the soma (**Fig 5T**). Total F-actin quantity, obtained from area under the curve analysis, showed that there was an overall increase in the F-actin quantity in *mts-IR* CI neurons (**Fig 5R**). Thus, in contrast to a recently published study that did not reveal differences in F-actin levels due to the knockdown of *mts* (Rui *et al*, 2020), our data indicates otherwise. In both CI and CIV neurons, *mts-IR* led to a reorganization of the F-actin cytoskeleton. Taken together, these results suggest that subtype-specific dendritic aberrations due to the disruption of the PP2A complex may be caused, at least in part, by regulatory effects on cytoskeletal architecture.

To independently validate that the observed defects on stable dendritic MTs were not due to non-specific effects of *mts* knockdown on expression of the mCherry-tagged microtubule associated protein (MAP) Jupiter, we conducted immunohistochemical (IHC) analyses using antibodies against the *Drosophila* MAP1B protein Futsch. Previous studies have shown that Futsch marks the population of stable microtubules (Cabernard & Doe, 2009; Weiner *et al*, 2016). IHC analyses demonstrate that *mts* knockdown in md neurons results in reductions in Futsch levels **(Fig S2 A, A’, B)**. MT stabilization is essential for dendritic development as MT destabilization leads to severe dendritic atrophy (Das *et al*, 2021). Acetylation of α-tubulin at lysine 40 residue has been associated with microtubule stabilization (Eshun-Wilson *et al*, 2019; Perdiz *et al*, 2011). Acetylation of α-tubulin shields MTs from mechanical aging and prevents breakage (Janke & Magiera, 2020). To determine if *mts* knockdown affects MT stabilization through changes of acetylated microtubule levels, we conducted immunohistochemistry by staining for acetylated α-tubulin in controls and *mts-IR* animals **(Fig S2 C-D’)**. Compared to controls, *mts* knockdown led to a decrease in acetylated tubulin levels in CIV neurons **(Fig S2 E)**. Collectively, these data support a functional role for PP2A in regulating the population of stable dendritic MTs.

Multi-channel imaging and IHC analyses in *mts* mutant neurons revealed significant reduction in the overall population of stable MTs along the dendrite **(Figs 5, S2)**. To investigate potential roles of PP2A in regulating MT dynamics, we examined the effects of *mts* knockdown on MT turnover. We performed time-lapse image of CIV neurons expressing an *α-tub-tdEos* reporter in control and *mts-IR* knockdown conditions. When small segments of the *α-tub-tdEos* expressing CIV dendrites are exposed to UV laser (405 nm), the dendritic segments undergo photoconversion from green to red. The duration of the photoconverted signal over time can be used to track MT stability and turnover. In controls, at 30 minutes after UV exposure, the photoconverted signal (**Fig 6 A”**), remains relatively unchanged from the initial photoconverted signal at time point 0 (**Fig 6 A’, A’’**), however the photoconverted signal drops by 26% at 60 minutes post-photoconversion (**Fig 6** **A’’’)**. In contrast, *mts-IR* dendrites showed a higher MT turnover rate when compared between various time points (**Fig 6** **B-B’’’, C)**. At 30 minutes after UV exposure, the photoconverted signal in neurons expressing *mts-IR* reduced by 33%, while at 60 minutes after photoconversion, the photoconverted signal reduced by 55% when compared to the initial photoconverted signal at time point 0. These findings suggest that MTs in *mts* mutant dendrites are more labile with higher turnover rates further supporting a role for Mts in MT stability.

**Figure 6:**
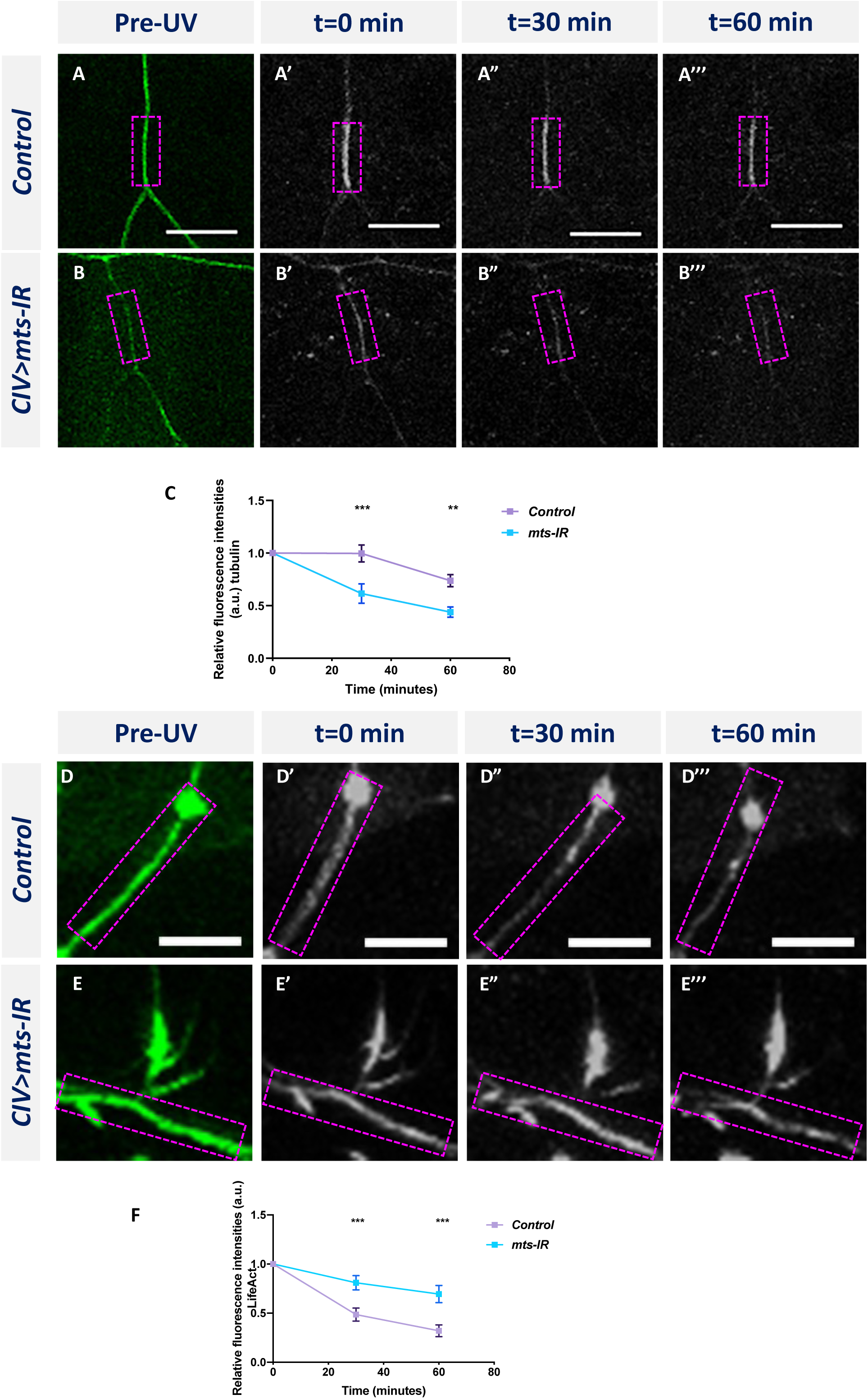
PP2A is required to regulate cytoskeletal dynamics: Time-lapse imaging of MT turnover in **(A-A’’’)** controls and **(B-B’’’)** *mts-IR*. **(C)** Quantitative analysis showing the turnover rate of MTs over a period of 60 minutes. Time-lapse imaging of F-actin turnover in **(D-D’’’)** controls and **(E-E’’’)** *mts-IR*. **(F)** Quantitative analysis showing the turnover rate of F-actin over a period of 60 minutes. Statistical tests performed: Two-way ANOVA with Sidak’s multiple comparisons (n = 8-13 per genotype). ***=p≤0.001, **=p≤0.01. For detailed genotypes see Table S1 and for detailed statistics see Table S2. Scale bars = 10 µm.

In contrast to its effect on MTs, *mts* knockdown led to a local reorganization in the F-actin signal of CIV neurons manifesting as a proximal shift in peak intensity towards the soma (**Fig 5** **G**). Moreover, in CI neurons, we observed an increase in F-actin levels at higher branch orders of *mts-IR* neurons relative to controls (**Fig 5** **Q**). To determine the effect of Mts disruption on F-actin turnover, we generated transgenic fly lines that expressed the F-actin binding peptide LifeAct (Riedl *et al*, 2008) fused to the photoconvertible tdEOS fluorescent protein. We then expressed *UAS-LifeAct:tdEos* in CIV neurons and studied the effect of *mts* knockdown on F-actin turnover. As with *α-tub-tdEOS*, we photoconverted a small segment of the dendrite and imaged the photoconverted signal at 0, 30, and 60 minutes. In controls, the photoconverted F-actin signal is very dynamic (**Fig 6** **D-D’’’, F)**. In controls, at 30 minutes after photoconversion, there is a 51% reduction in the photoconverted red signal which is further reduced to 68% of baseline (t=0 min) levels at 60 minutes after photoconversion. In contrast, the F-actin signal appears to be relatively stable in *mts-IR* with very little loss in the photoconverted signal even at 60 minutes post-photoconversion. Only 19% of the photoconverted red signal is lost in *mts-IR* animals at 30 minutes after photoconversion and the reduction in photoconverted signal is only 31% of baseline levels at 60 minutes after photoconversion (**Fig 6** **E-E’’’, F)**. A between-animals comparison shows a significant difference in the photoconverted F-actin signal between control and *mts-IR* at both 30 minutes as well as 60 minutes after photoconversion (**Fig 6** **F)**. These data suggest that disruption of Mts leads to increased stabilization of F-actin thereby implicating PP2A in F-actin turnover dynamics.

### Mts is required for MT polarity

Dendritic development is dependent upon MT motor based trafficking regulated by MT polarity (Lefebvre *et al*, 2015; Nanda *et al*, 2017). In *Drosophila* md neuron dendrites, MTs are arranged with their minus ends distal and plus ends proximal relative to the cell body (Rolls *et al*, 2007; Stone *et al*, 2008). In light of the regulatory role of PP2A on MT stability, we sought to determine how PP2A may impact MT polarity. To this end, we examined dynamic localization of the plus-end MT marker EB1 in controls and *mts* mutants. In control CIV primary dendrites, 92% of the EB1::GFP comets move in a retrograde direction towards the cell body (**Fig 7** **A, C)**; however, in CIV *mts* knockdown conditions, EB1 comets show a reversal in polarity with 91% of the comets moving in an anterograde direction away from the cell body (**Fig 7** **B, C)**. In control CI primary dendrites, 90% of the comets move towards the cell body while in CI *mts-IR* primary dendrites, only 52% of the comets move towards the cell body **(Fig S3 A, B,** **Fig 7** **K)**. Thus, *mts-IR* disrupts MT polarity in both CI and CIV neurons. In both CI and CIV neurons, *mts* knockdown also led to an increase in the speed of the EB1::GFP comets (**Fig 7** **H, L)**, however, the comet track length in both CI and CIV *mts-IR* neurons remained unaffected relative to controls (**Fig 7** **G, N)**. In contrast to primary dendritic branches, md neuron higher order branches exhibit mixed polarity. In both CI and CIV neurons, controls as well as *mts-IR* showed no significant difference in MT polarity **(Fig S3 C, D)**, indicative of a role for Mts in regulating MT polarity of primary dendritic branches.

**Figure 7:**
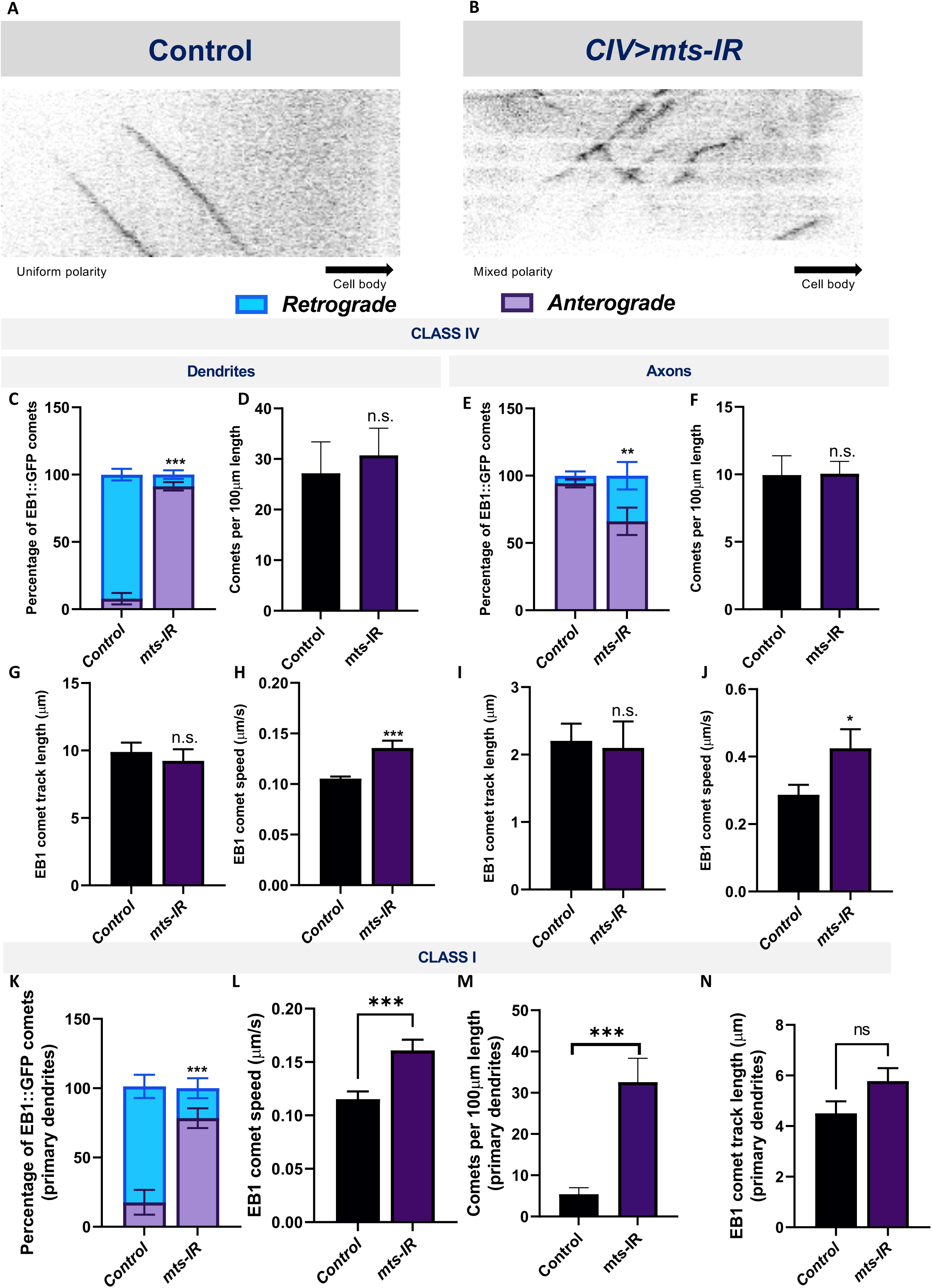
PP2A disruption leads to reversal of MT polarity: Kymographs of EB1::GFP comet trajectories in CIV neurons in **(A)** control and **(B)** *mts-IR*. Relative to controls which show minus end out microtubule polarity in proximal dendrites and plus end out polarity in axons, knockdown of *mts* leads to reversal of microtubule polarity in md neuron dendrites **(C, K)** and axons **(E).** Further, *mts* knockdown leads to an increase in EB1 comet velocities compared to controls in both CI and CIV md neurons in both dendrites and axons **(H, J, L).** Comet track length and number of comets are unaffected due to knockdown of *mts* in CIV dendrites **(D, F, G, I)**, while in CI, there is significant increase in number of comets while track length is unchanged **(M, N)**. Identical settings for laser intensity and other image capture parameters were applied for comparisons of control vs. experimental samples. Statistical tests performed: **(C, E, K)** two-way ANOVA with Sidak’s or Dunnett’s multiple comparison test (n = 42-72 comets), **(D, F, G-J, L-N)** Student’s t-test or Mann-Whitney U test (n = 9-67 per genotype). ***=p≤0.001, **=p≤0.01, *=p≤0.05. For detailed genotypes see Table S1 and for detailed statistics see Table S2.

In contrast to dendrites, axons contain microtubules with primarily plus-end-out orientation (Stone *et al*, 2008). In control CI and CIV axons, EB1 comets primarily move away from the cell body (**Fig 7** **E, Fig S3 E, G, I)**. However, knockdown of *mts* in either CI or CIV neurons led to a mixed polarity with a significant number of EB1 comets moving towards the cell body (**Fig 7** **E, Fig S3 F, H, I)**.

Similar to what was observed in dendrites, *mts* knockdown led to an increase in comet speed but no change in track length or number of comets (**Fig 7** **F, I, J)**. In CI axons, *mts* knockdown led to an increase in the number of comets compared to controls **(Fig S3 J**), similar to what was observed in CI primary dendrites (**Fig 7** **M)**. These results suggest PP2A also regulates axonal MT polarity.

γ-tubulin forms a part of the centrosome which is required for microtubule assembly (Raynaud-Messina *et al*, 2001). The *Drosophila* genome encodes two γ-tubulins: γ-tubulin 23C and γ-tubulin 37C (Raynaud-Messina *et al*, 2001). Although both are expressed at the centrosome, γ-tubulin 37C expression is primarily restricted to the ovaries, while γ-tubulin 23C has a more global expression pattern and is found in most adult fly cells (Wiese, 2008). Previous studies have shown that loss, as well as overexpression, of γ-tubulin 23C reverses polarity of EB1 comets in CI neurons (Nguyen *et al*, 2014). To determine if the disruption in MT polarity in *mts* knockdown could be due to its effects on γ-tubulin localization, we expressed γ-tubulin 23C-GFP in CIV neurons in control and *mts-IR* conditions. As previously reported, in controls, γ-tubulin 23C-GFP appeared as puncta that are primarily localized at dendritic branch points (Nguyen *et al*, 2014) (**Fig 8** **A-A”)**. Knockdown of *mts* severely disrupted γ-tubulin 23C-GFP localization (**Fig 8** **B-B”)**. Quantitative analysis showed a significant reduction in γ-tubulin 23C-GFP localization along the dendritic arbor (**Fig 8** **C)**.

**Figure 8:**
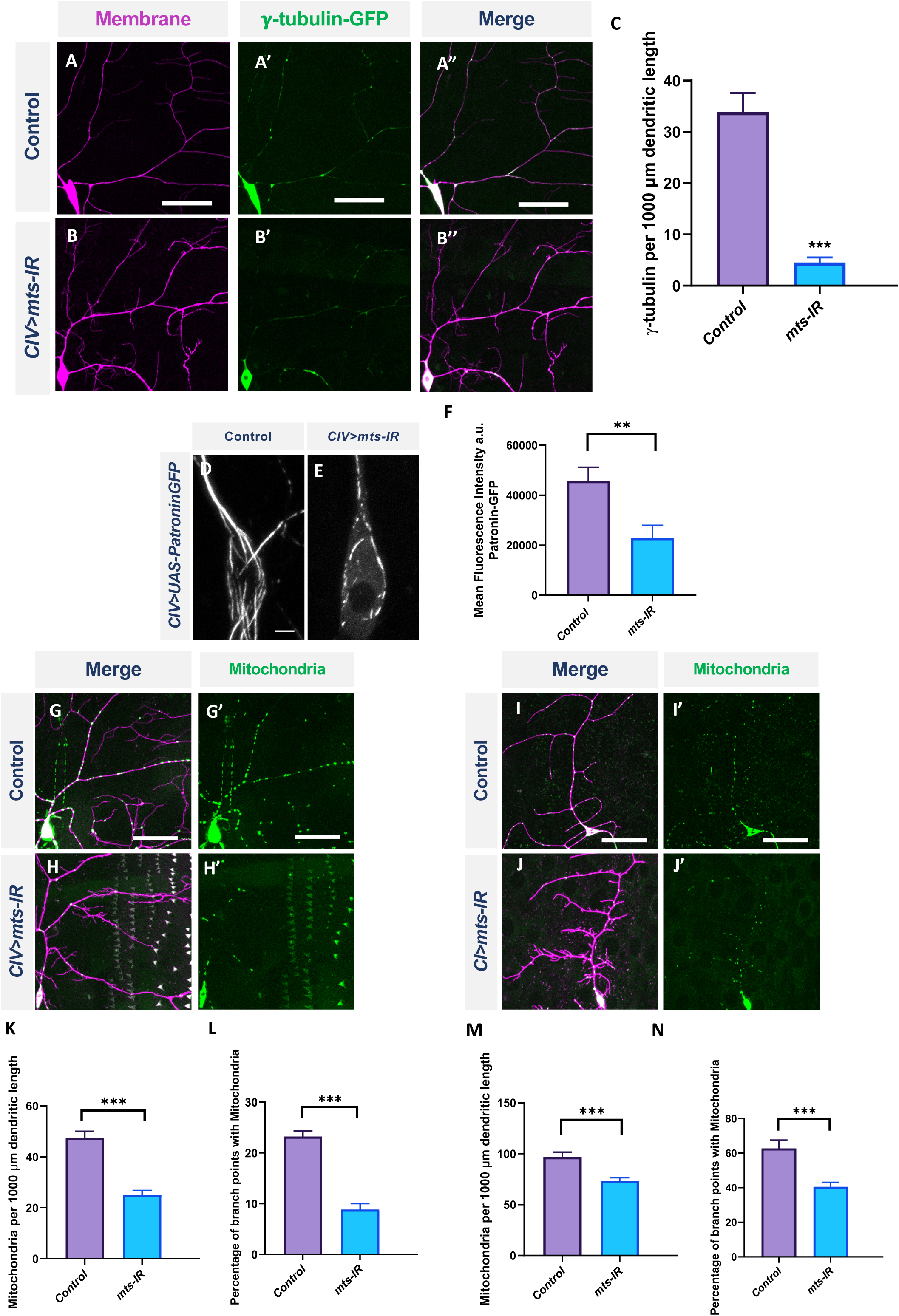
PP2A is required for organelle trafficking in md neurons: Representative images of CIV neurons in **(A-A’’)** control and **(B-B’’)** *mts-IR* expressing γ-tubulin-GFP under the control of a CIV-GAL4 driver. **(C)** Quantitative analysis showing γ-tubulin localization along the dendrite normalized to dendrite length. Representative images of CIV neurons in **(D)** control and **(E)** *mts-IR* expressing Patronin-GFP under the control of CIV-GAL4 driver. **(F)** Quantification of mean fluorescence intensities of Patronin-GFP normalized to area. Representative images of CIV and CI neurons in **(G, G’) (I, I’)** control and **(H, H’) (J, J’)** *mts-IR* animals, respectively, expressing *UAS-mito-GFP* under the control of CIV or CI GAL4 driver. **(K, M)** Quantitative analysis showing mitochondria localization along the dendrite normalized to dendrite length and **(L, N)** percentage of branch points with mitochondria. Statistical tests performed: **(C, F, K-N)** Student’s t-test or Mann-Whitney U test (n = 7-14 per genotype). ***=p≤0.001, **=p≤0.01. For detailed genotypes see Table S1 and for detailed statistics see Table S2. Scale bars: **(A-A”, G, G’, I, I’)** = 50μm, **(D) =** 5μm.

Patronin is a microtubule minus end binding protein that belongs to the CAMSAP/Patronin/Nezha family of proteins (Akhmanova & Hoogenraad, 2015). The *Drosophila* genome encodes a single Patronin gene which has been shown to associate with the minus end of MTs and prevent MT disassembly in *Drosophila* S2 cells *in vitro* (Goodwin & Vale, 2010; Akhmanova & Hoogenraad, 2015). A recently published study showed that expression of YFP-tagged Patronin in md neurons marked the whole microtubule matrix and that the loss of Patronin led to reversal of MT polarity in md neurons (Feng *et al*, 2019). To determine if *mts* knockdown impacts Patronin expression and localization in md neurons, we co-expressed Patronin-YFP in combination with *mts-IR*. As previously reported (Feng *et al*, 2019), Patronin-YFP appeared as long lattices along the dendrite and throughout the cell body in control neurons (**Fig 8** **D)**. However, in *mts-IR* conditions, the Patronin signal was reduced and the lattice was disrupted (**Fig 8** **E, F)**. These data indicate that Patronin is also affected by loss of Mts, and reveal another potential mechanism that may underlie changes in MT polarity in *mts* mutant neurons.

### PP2A is required for cytoskeletal-based organelle trafficking

Mitochondrial dysfunction leads to severe dendritic atrophy in CIV md neurons while CI neurons are largely resilient to mitochondrial fragmentation (Tsubouchi *et al*, 2009; Tsuyama *et al*, 2017). In addition, disruption in mitochondrial trafficking along the dendrites is associated with reduced dendritic complexity (López-Doménech *et al*, 2016; Das *et al*, 2021). To study the effects of PP2A disruption in mitochondrial localization, we expressed *UAS-mito-GFP* in CIV and CI neurons with the simultaneous knockdown of *mts*. Knockdown of *mts* led to a significant reduction in mitochondrial localization along the dendritic arbor in both CI and CIV neurons (**Fig 8** **G-J’)**, both at branch points and along the arbor (**Fig 8** **K-N)**. Taken together, these data indicate that Mts is required for proper mitochondrial localization in md neurons.

Previous studies in *Drosophila* md neurons have demonstrated that Golgi outposts are trafficked along the dendrites and are localized preferentially at branch points where they act as sites of microtubule nucleation and also regulate terminal dendrite dynamics (Nanda *et al*, 2017; Iyer *et al*, 2013b; Stone *et al*, 2008; Ori-McKenney *et al*, 2012). Golgi trafficking along the dendrite is mediated by Dynein and dependent on the autoinhibition of axon-targeted Kinesin-1, and is thus dependent on MT polarity and cellular compartmentalization (Arthur *et al*, 2015; Jaarsma & Hoogenraad, 2015; Kelliher *et al*, 2018). To determine the consequences of disrupted MT architecture resulting from defects in PP2A, we examined the distribution of satellite Golgi outposts in both CI and CIV md neurons. We expressed the medial Golgi marker, ManII, in CI or CIV with simultaneous knockdown of *mts*. In CIV neurons, *mts* knockdown inhibited Golgi outpost trafficking onto dendrites and led to a reduction in Golgi outposts at branch points and along the dendrites (**Fig 9** **A-B’, E, F)**. In contrast, in CI neurons, disruption in *mts* led to an increased Golgi trafficking along the dendrites compared to controls (**Fig 9** **C-D’, G)**. Golgi localization at branch points remained unaffected in *mts-IR* animals compared to controls (**Fig 9** **H)**. Collectively, these data show that PP2A regulates Golgi trafficking in a subtype-specific manner: in the simpler CI neurons, PP2A restricts Golgi outpost trafficking while in the more complex CIV neurons, PP2A is required to promote Golgi trafficking.

**Figure 9:**
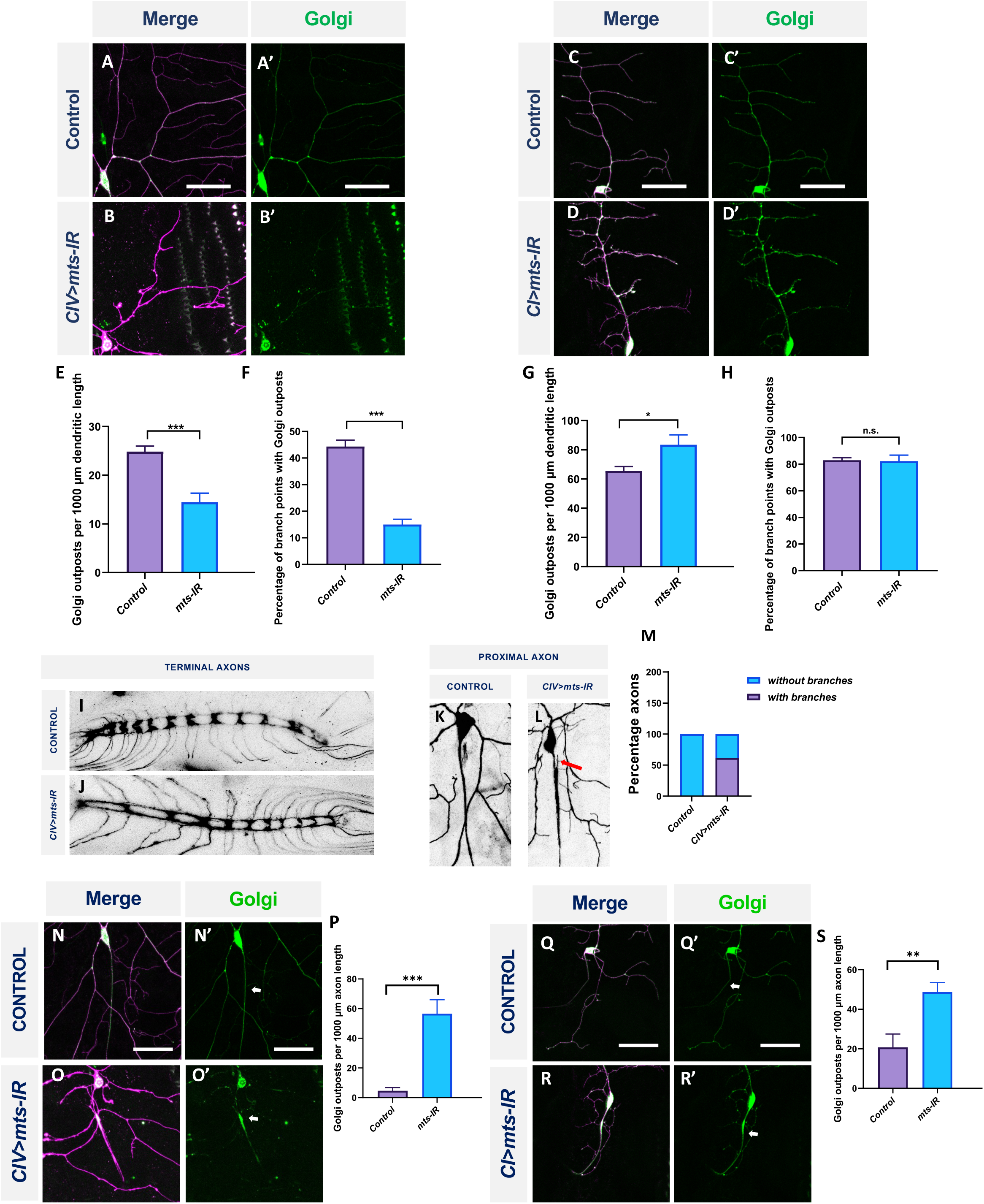
PP2A has cell-type specific effects of Golgi outpost localization in md neurons: Representative images of CIV and CI neurons in **(A, A’) (C, C’)** control and **(B, B’) (D, D’)** *mts-IR* animals, respectively, expressing *UAS-MANII-eGFP* under the control of CIV or CI GAL4 driver. **(E, G)** Quantitative analysis showing Golgi outposts localization along the dendrite normalized to dendrite length and **(F, H)** percentage of branch points with Golgi outposts. Representative images of the CIV axon terminals in the ventral nerve cord (VNC) in **(I)** control and **(J)** *mts-IR* animals. Representative images of the proximal axons of CIV neurons in **(K)** control and **(L)** *mts-IR* animals. Arrow indicates ectopic axon branching. **(M)** Quantitative analysis showing the percentage of proximal axons with branching in control and *mts-IR* neurons. Representative images of CIV and CI axons in **(N, N’) (Q, Q’)** control and **(O, O’) (R, R’)** *mts-IR* animals, respectively, expressing *UAS-MANII-eGFP* under the control of CIV or CI GAL4 driver. Arrows indicate axons. **(P, S)** Quantitative analysis showing Golgi outposts localization along the axon normalized to length. Statistical tests performed: **(E-H, M, P, S)** Student’s t-test or Mann-Whitney U test (n = 10-19 per genotype). ***=p≤0.001, *=p≤0.05. For detailed genotypes see Table S1 and for detailed statistics see Table S2. Scale bar = 50μm.

### PP2A is required to maintain neuronal polarity

Golgi outpost localization is highly polarized and restricted to the dendrites (Horton *et al*, 2005; Ye *et al*, 2007). However, we discovered that expression of *mts-IR* led to aberrant localization of Golgi outposts in axons. In both CI and CIV neurons, *mts* knockdown led to Golgi mislocalization in the proximal axons emerging from the cell body (**Fig 9** **N-O’, Q-R’)**. Quantitative analysis showed that compared to controls, *mts* knockdown led to increased Golgi localization in the axons of CI and CIV neurons, respectively (**Fig 9** **P, S)**. Gene mutations that lead to mislocalization of Golgi in the axons are also associated with ectopic branching in the axons (Kelliher *et al*, 2018; Ye *et al*, 2007; Arthur *et al*, 2015; Zheng *et al*, 2008). Therefore, we examined the proximal and distal axonal segments of CIV neurons in both controls and *mts-IR* animals. While a substantial number of CIV neurons in *mts* knockdown conditions showed ectopic branching in the proximal axon (**Fig 9** **K-M)**, we did not observe any gross anatomical defects in the distal axon terminals in the ventral nerve cord (**Fig 9** **I, J).**

Golgi mislocalization in axons led us to hypothesize that Mts may be required to maintain neuronal compartmentalization in md neurons. To test this hypothesis, we expressed DenMark and Synaptotagmin under the control of a CI *GAL4* driver in control and *mts-IR* conditions. DenMark is the ICAM5 molecule fused to mCherry which traffics to dendrites, whereas Synaptotagmin is a presynaptic marker fused to GFP (Nicolai *et al*, 2010). Consistent with previous reports (Nicolai *et al*, 2010), in control neurons, DenMark signal is predominantly localized at the cell body and dendrites **(Fig S4 A)**. However, in *mts-IR* conditions, DenMark signal extends into the axons **(Fig S4 B)**. In control CI md neurons, there is no discernible Synaptotagmin signal in the distal dendrites **(Fig S4 C)**; however, quantitative analysis showed a significant increase in Synaptotagmin signal in distal dendrites due to the knockdown of *mts* **(Fig S4 D, E)**. In addition, loss of *mts* also led to an increase in the DenMark signal in the axons **(Fig S4 F)**. These results are consistent with what was previously reported for CIV neurons due to the disruption of Mts (Rui *et al*, 2020). Taken together, these data support a role for PP2A in maintaining neuronal compartmentalization in md neurons.

### PP2A regulates dendritic morphology in CI neurons through genetic interaction with FoxO

The Forkhead box TF, FoxO, has been previously implicated in md neuron dendritic development (Sears & Broihier, 2016). FoxO is involved in regulating a number of cellular processes and its functional activity is largely regulated by posttranslational modifications including phosphorylation (Zhang *et al*, 2011). Dephosphorylation of FoxO by PP2A is associated with its translocation into the nucleus and transcriptional control of numerous genes (Tzivion *et al*, 2011). To determine how PP2A may mechanistically regulate subtype-specific dendritic architecture, we investigated FoxO as a putative target of PP2A in CI md neurons. Quantitative analysis showed that *foxo* overexpression led to increases in branches, and branch density but reduction in the total dendritic length when compared to controls (**Fig 10** **A, B, F-H)**. This was phenotypically similar to what we observed for *mts* knockdown in CI neurons (**Fig 10** **J**). In addition, IHC analyses showed that *mts* knockdown in CI md neurons led to an increase in FoxO expression compared to controls (**Fig 10** **C-D’, E)**. Furthermore, co-overexpression of *foxo* and *mts* together in CI neurons led to a significant reduction in number of branches, and branch density (**Fig 10** **K, M. N)**. Quantitative analyses showed that co-expression of *mts* and *foxo* returned all the dendritic parameters to the levels comparable to controls (**Fig 10** **M, N)**.

**Figure 10:**
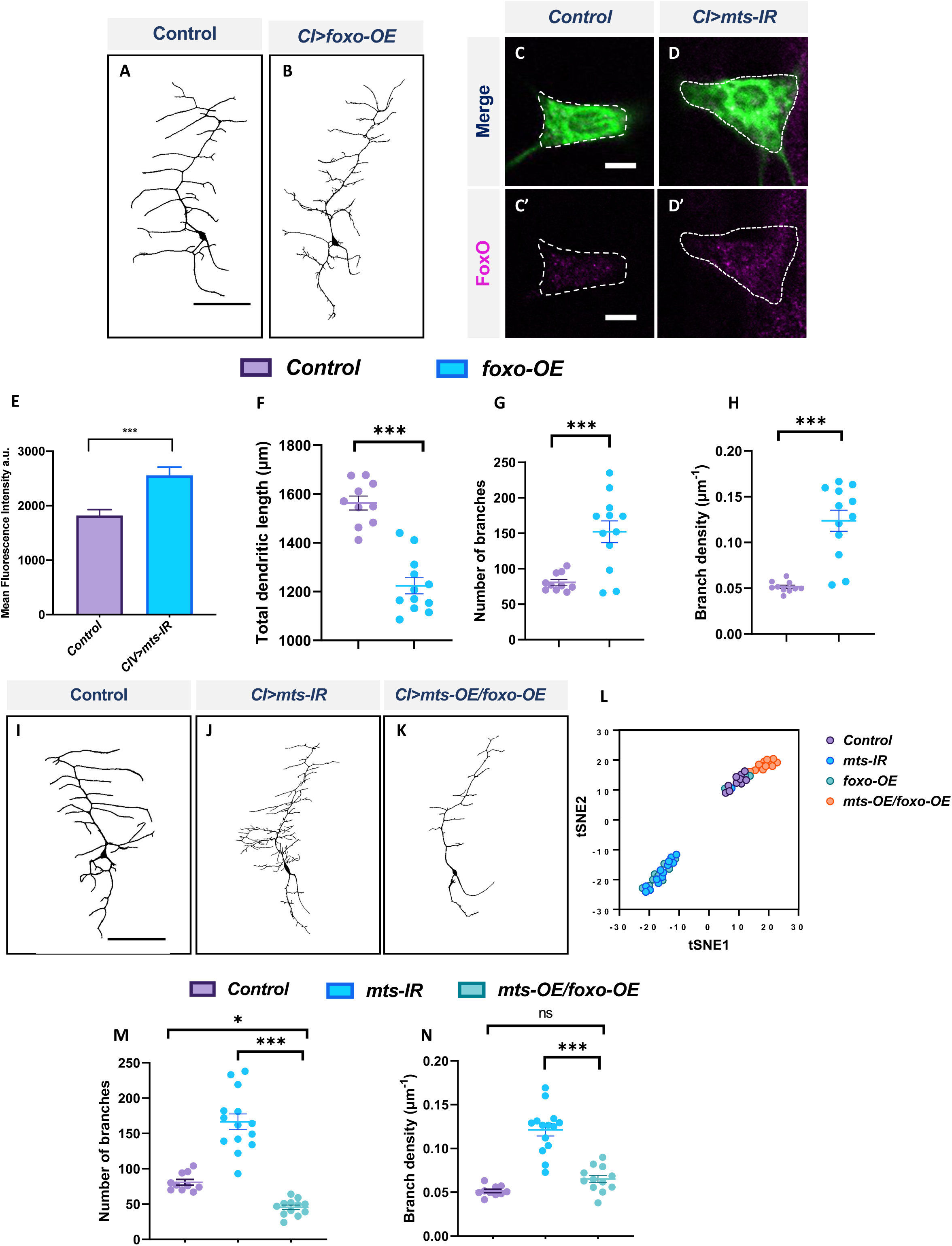
PP2A and FoxO genetically interact in CI md neurons: Representative images of CI neurons in **(A)** control and **(B)** *foxo-OE*. Representative images showing immunohistochemical analysis of FoxO expression in **(C, C’)** control, and **(D, D’)** *mts-IR* expressing CI neurons. **(E)** Quantitative analysis of mean fluorescence intensities of FoxO normalized to area in CI neurons. **(F-H)** Quantitative morphometric analyses. Representative images of CI neurons in **(I)** control, **(J)** *mts-IR* and **(K)** *mts-OE/foxo-OE* conditions. **(M, N)** Quantitative morphometric analyses. **(L)** tSNE plot showing the clustering of control, *mts-IR, foxo-OE, and mts-OE/foxo-OE* neurons. Statistical tests performed: **(F-H)** Student’s t-test (n = 10-29 per genotype); **(M, N)** One-way ANOVA with Sidak’s multiple comparison (n = 10-14 per genotype). ***=p≤0.001, **=p≤0.01, *=p≤0.05. For detailed genotypes see Table S1 and for detailed statistics see Table S2. Scale bars = 100 µm for **(A, I)**, and scale bar = 5 µm for **(C-C’)**.

To better visualize the effects of co-overexpression of *mts* and *foxo* in rescuing the morphological defects caused by the overexpression of *foxo* alone, we conducted tSNE analysis. T-distributed stochastic neighbor embedding, or tSNE, is a machine learning algorithm that allows us to visualize high dimensional data in two-dimensional space ( Maaten & Hinton, 2008; Maaten, 2014). tSNE analysis reveals that *mts-IR* and *foxo-OE* phenotypes largely co-cluster together indicating that both these perturbations exert similar effects on dendritic morphology in CI md neurons. Neurons co-overexpressing *mts* and *foxo* cluster separately from *mts-IR* or *foxo-OE*. Instead these neurons phenotypically cluster with control CI neurons, showing that simultaneous overexpression of *mts* and *foxo* can suppress dendritic defects caused by *foxo-OE* alone (**Fig 10** **L)**. Collectively, these data suggest that FoxO and PP2A genetically interact and that FoxO may be a regulatory target of PP2A phosphatase activity in CI md neurons.

### PP2A genetically interacts with β-tubulin 85D in md neurons

Tubulin phosphorylation occurs on both the α- and β-subunits at serine, threonine and tyrosine residues (Wloga *et al*, 2017; Yu *et al*, 2015). Dephosphorylation of β-tubulin is required for tubulin polymerization and mutations in S172 of β-tubulin affects microtubule dynamics (Wloga *et al*, 2017; Caudron *et al*, 2010). Further, studies have shown that dephosphorylation of βIII tubulin in bovine brain requires PP2A (Khan & Ludueña, 1996).

To determine the putative role of β-tubulin in regulating dendritic architecture, we investigated the impact of *β-tubulin* knockdown on CIV dendritic morphology. The *Drosophila* genome encodes five *β-tubulin* genes (Fyrberg & Goldstein, 1990; Findeisen *et al*, 2014). Knockdown of *β-tubulin85D* and *β-tubulin56D* severely disrupted dendritic morphology (**Fig 11** **B, D, E, Fig S5 B, F, G)**, whereas knockdown of the *β-tubulin60D* and CG32396 led to a modest increase in total dendritic length without affecting number of branches. Knockdown of *β-tubulin97EF* did not affect dendritic morphology **(Fig S5 A, C-G)**.

**Figure 11:**
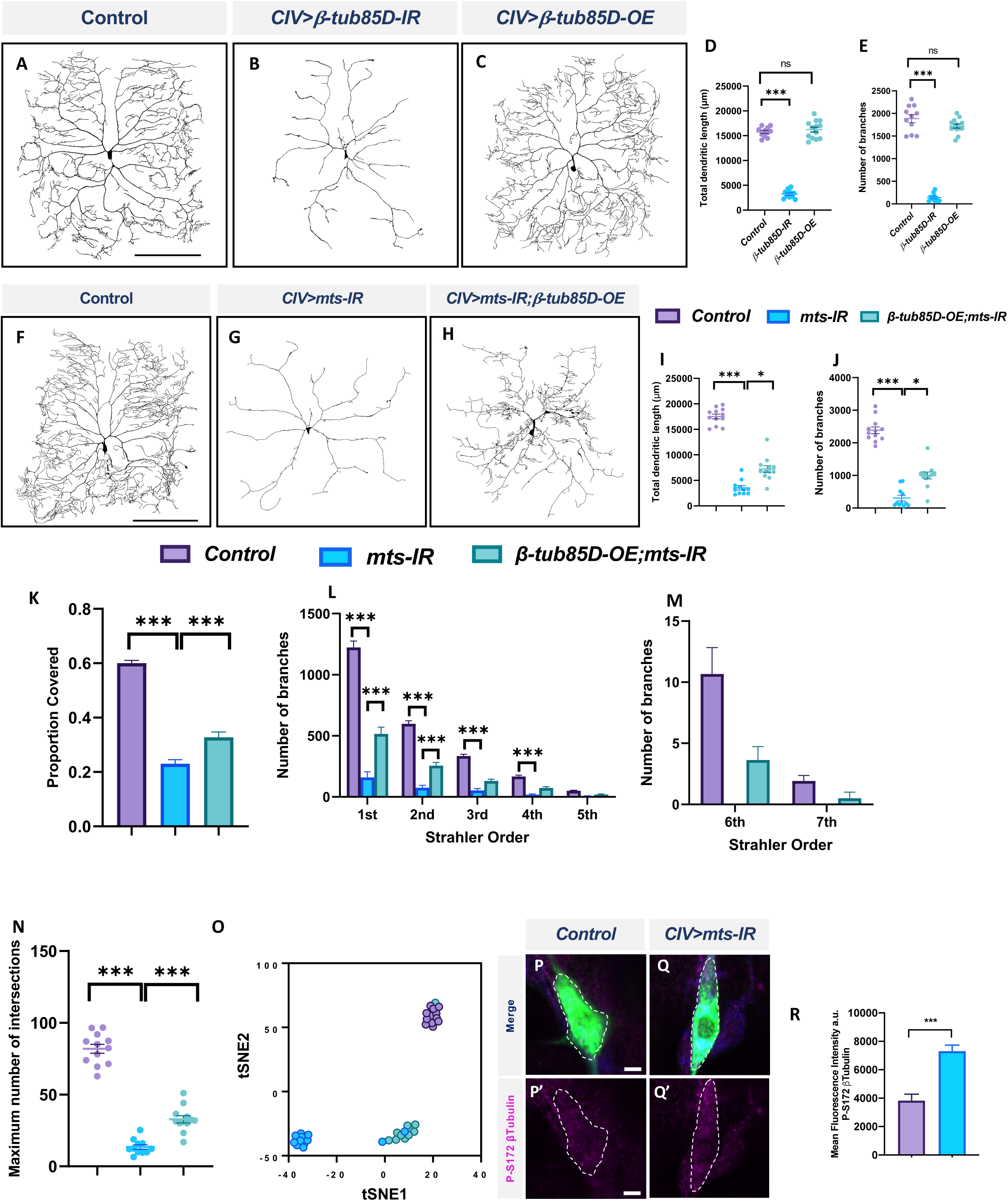
PP2A and β-tubulin 85D genetically interact in CIV md neurons: Representative images of CIV neurons in **(A)** control, **(B)** *β-tubulin85D-IR*, and **(C)** *β-tubulin85D-OE*. **(D, E)** Quantified morphometric analyses. Representative images of CIV neurons in **(F)** control, **(G)** *mts-IR*, and **(H)** *β-tubulin85D-OE/mts-IR* animals. **(I-K, N)** Quantitative morphometric analyses. **(O)** tSNE plot showing clustering of control, *mts-IR*, and *β-tubulin85D-OE/mts-IR* expressing neurons. Representative images showing immunohistochemical analysis of p-S172 β-tubulin expression in CIV neurons in **(P, P’)** control and **(Q, Q’)** *mts-IR* conditions. **(R)** Quantitative analysis of mean fluorescence intensities of p-S172 β-tubulin normalized to area in CIV neurons. Statistical tests performed: **(D, E)** One-way ANOVA with Dunnett’s multiple comparison (n = 13 per genotype), **(I-K, N)** One-way ANOVA with Sidak’s multiple comparison and Kruskal-Wallis with Dunn’s multiple comparison (n = 10-12 per genotype), **(L, M)** two-way ANOVA with Tukey’s multiple comparison (n = 10-12 per genotype), **(R)** Student’s t-test (n= 24-32 per genotype). ***=p≤0.001, *=p≤0.05. For detailed genotypes see Table S1 and for detailed statistics see Table S2. Scale bar = 200 µm for **(A, F);** scale bar = 5 µm for **(P-P’’)**.

The effects on dendritic morphology due to the knockdown of both *β-tubulin56D* and *β-tubulin85D* were phenotypically similar to those observed with *mts* knockdown in CIV md neurons. However, *β-tubulin56D* encodes the *Drosophila* βI tubulin which is maternally expressed (Rudolf *et al*, 2012). We therefore decided to pursue *β-tubulin85D* as a putative target for PP2A in md neurons.

To determine if *β-tubulin85D* and *mts* genetically interact with one another, we conducted rescue experiments. Overexpression of *β-tubulin85D* with the simultaneous knockdown of *mts* partially rescued the phenotypic defects caused by the knockdown of *mts* alone (**Fig 11** **F-H)**. Overexpression of *β-tubulin85D* by itself did not have an effect on CIV dendritic morphology (**Fig 11** **C-E)**. *mts-IR;β-tubulin85D-OE* neurons showed an increase in total dendritic length, number of branches, and proportion of area covered when compared to *mts-IR* alone (**Fig 11** **I-K)**. Knockdown of *mts* led to a significant reduction in 1^st^ to 4^th^ order branches and complete loss of 6^th^ and 7^th^ order branches. Overexpression of *β-tubulin85D* in this sensitized background led to a partial rescue of the 1^st^ and 2^nd^ order branches and recovery of 6^th^ and 7^th^ order branches (**Fig 11** **L, M)**. Sholl analysis demonstrated that compared to *mts-IR,* there was an increase in the maximum number of intersections in *mts-IR;β-tubulin85D* animals suggesting a partial rescue in the dendritic complexity that was lost with the knockdown of *mts* (**Fig 11** **N)**. tSNE analysis revealed *mts-IR* neurons phenotypically cluster separately and distantly from control neurons due to the severe disruption in dendritic morphology. In contrast, neurons expressing both *UAS-β-tubulin85D* and *UAS-mts-IR* clustered separate from *mts-IR* neurons and closer to the control neurons with one neuron clustering with controls, suggesting that overexpression of *β-tubulin85D* in CIV neurons can partially rescue the phenotypic defects caused by the knockdown of *mts-IR* alone (**Fig 11** **O)**. Further, IHC analyses revealed an increase in the levels of phosphorylated β-tubulin (phospho-Ser172) in CIV *mts-IR* md neurons compared to control suggesting that *β-tubulin85D* is a target of PP2A phosphatase activity in these neurons (**Fig 11** **R-R)**.

We next sought to determine whether *β-tubulin85D* may be a common target of PP2A regulation in md neurons given that both CIV and CI neurons knocked down for *mts* exhibit defects in MTs. Knockdown of *β-tubulin85D* in CI md neurons led to an increase in ectopic dendritic branching. (**Fig 12** **A, B)**. Neurons expressing *UAS-β-tubulin85-IR* showed an increase in number of branches and branch density compared to control, but did not affect total dendritic length (**Fig 12** **C, D, E)**. Strahler order analysis revealed a significant increase in 1^st^, 2^nd^ and 4^th^ order branches similar to that observed in *mts-IR* (**Fig 12 F**). Co-expression of *UAS-β-tubulin85D* and *UAS-mts-IR* rescued the phenotypic defects of *mts-IR* (**Fig 12 G-I**). Neuromorphometric analyses revealed that *mts-IR; β-tubulin85D-OE* neurons showed a reduction in number of branches, and branch density when compared to *mts-IR* (**Fig 12 K, L**). Strahler analysis revealed a significant reduction in the 1^st^, 2^nd^ and 3^rd^ order dendrites in neurons co-expressing *UAS-β-tubulin85D* and *mts-IR* compared to *mts-IR* alone (**Fig 12 M**). tSNE analysis revealed that control neurons and *mts-IR;β-tubulin85D-OE* neurons phenotypically clustered more closely together than *mts-IR* neurons, further supporting the hypothesis that *β-tubulin85D* overexpression in this sensitized background can rescue phenotypic defects of *mts-IR* alone (**Fig 12 J**). Collectively, these data suggest that β-tubulin85D is a target of PP2A in CI md neurons.

**Figure 12:**
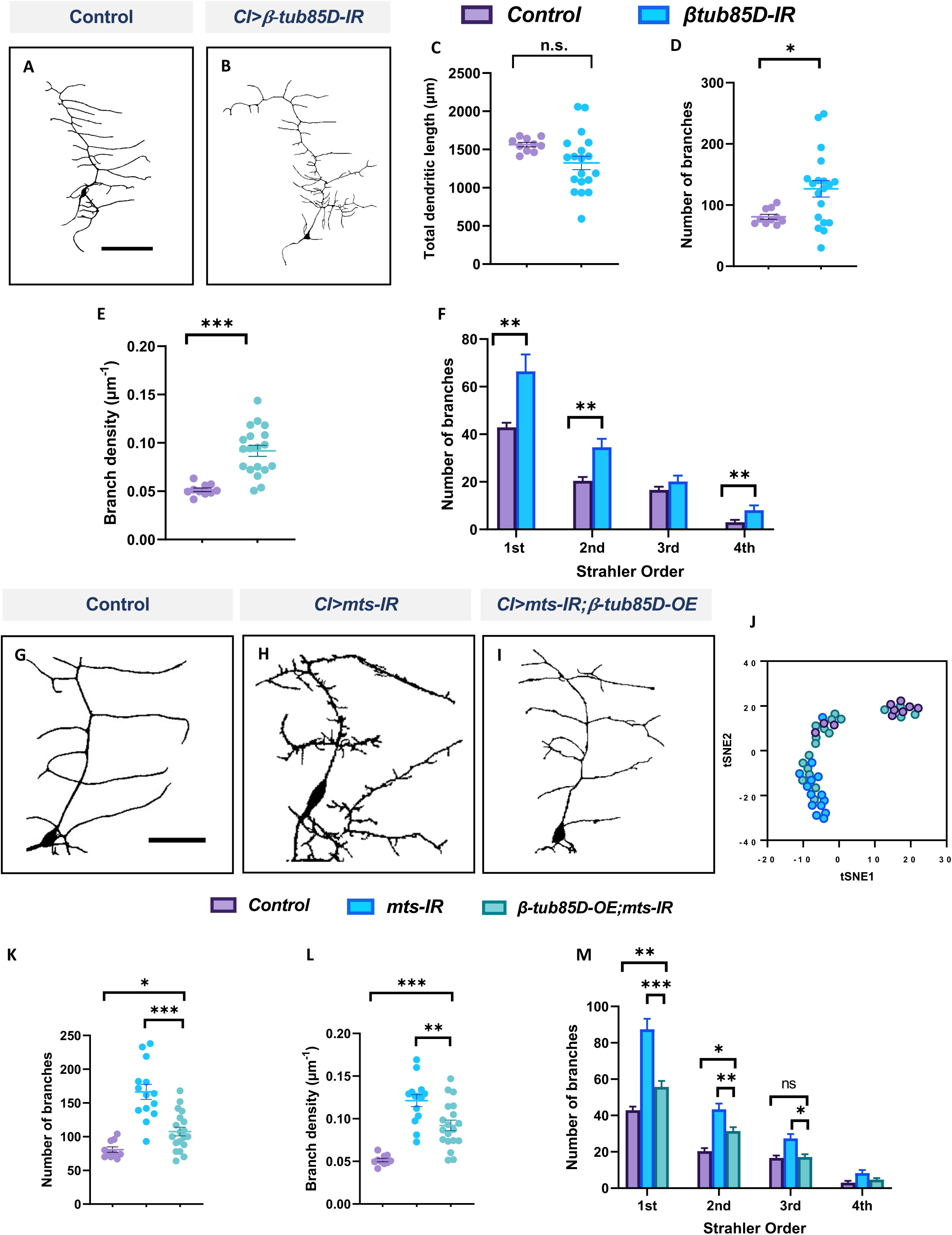
PP2A genetically interacts with β-tubulin85D in CI md neurons: Representative images of CI neurons in **(A)** control and **(B)** *β-tubulin85D-IR*. **(C-F)** Quantitative morphometric analyses. Representative images of CI neurons in **(G)** control, **(H)** *mts-IR* and **(I)** *β-tubulin85D-OE/mts-IR* animals. **(J)** tSNE plot showing clusters of control, *mts-IR*, and *β-tubulin85D-OE/mts-IR* neurons. (**K-M**) Quantitative morphometric analyses. Statistical tests performed: **(C-E)** Student’s t-test (n = 10-19 per genotype), **(F)** Unpaired t-test with False Discovery rate correction (n = 10-19 per genotype), **(K, L)** One-way ANOVA with Sidak’s multiple comparison or Kruskal-Wallis with Dunn’s multiple comparison (n = 10-19 per genotype), **(M)** Two-way ANOVA with Tukey’s multiple comparison test (n = 10-19 per genotype). ***=p≤0.001, **=p≤0.01, *=p≤0.05. For detailed genotypes see Table S1 and for detailed statistics see Table S2. Scale bar = 100 µm for **(A)**, and scale bar = 50 µm for **(G)**.

In order to further examine the putative functional significance of the phosphorylation state of β-tubulin85D, we generated novel transgenic *Drosophila* lines in which the S172 and/or T219 residues of β-tubulin85D were mutated to either glutamic acid (E) or alanine (A). Mutation of the S172 and T219 residues to glutamic acid generates a mutant form of the protein that mimics its phosphorylated state (phosphomimetic), and mutation of the S172 and T219 residues to alanine generates a mutant form that mimics its dephosphorylated state (phospho-resistant) **(Fig S6 A)**. If phosphorylation of β-tubulin85D impedes MT polymerization or dynamics, we hypothesized that expression of the phosphomimetic form of the protein would lead to dendritic hypotrophy. Expression of the phosphomimetic *β-tubulin85D-S172E_T219E* in CIV md neurons led to phenotypic disruptions with reductions in total dendritic length, and number of branches **(Fig S6 B, D-F)**. In contrast, expression of the phosphoresistant *β-tubulin85D-S172A_T219A* in CIV md neurons did not affect dendritic morphology **(Fig S6 B, C, E-F)**. These data suggest that dephosphorylation of β-tubulin85D at S172 and/or T219 is required for dendritogenesis in CIV md neurons.

## Discussion

### PP2A is required for cell-type specific dendritic architecture

Maintaining proper dendritic morphology is essential for neurons to form a functioning nervous system. The broad significance of dendritic form and regulation of cytoskeletal architecture in neural function is underscored by the wide spectrum of neurological and neurocognitive disorders that have been linked to disruptions in these processes (Franker & Hoogenraad, 2013). The present study reveals that the PP2A phosphatase has a differential regulatory effect on dendritic morphology in different classes of *Drosophila* md neurons. Consistent with recent studies (Rui *et al*, 2020; Wolterhoff *et al*, 2020, Wang et al., 2020), we discovered that loss of catalytic subunit *mts* or scaffolding subunit *PP2A-29B* in CIV md neurons severely disrupt dendritic morphology. However, disruption of only one regulatory subunit, *wdb,* had a notable effect on larval dendritic morphology, suggesting Wdb functions as a key regulatory subunit in these neurons during larval dendritic development. Recent studies identified the role of *tws* in regulating dendritic pruning in CIV neurons (Rui *et al*, 2020; Wolterhoff *et al*, 2020) and overexpression of Tws was also found to disrupt CIV dendrite morphogenesis (Wang et al., 2020). However, our analyses showed that knockdown of *tws* had a relatively mild effect on dendritic morphology in wandering third instar larvae. In addition, knockdown of *wrd* also affected dendritic morphology but the phenotypic effects of *tws* and *wrd* knockdown were subtle compared to the effects of *wdb* knockdown. This suggests that regulatory subunits may play differential/combinatorial roles in dendritic arborization vs. pruning depending on stage of development.

We found that there was subtype-specific reliance on PP2A for the formation of dendritic arbors. Disruption of *mts* and *PP2A-29B* led to an increase in dendritic complexity in CI md neurons, but a decrease in dendritic complexity in CIV md neurons. In CI md neurons, knockdown of *wdb* led to a mild reduction in dendritic length and an increase in branches. Knockdown of *wrd* and *tws* did not affect dendritic morphology while the loss of *CG4733* led to a mild reduction in total dendritic length. This suggests that Wdb may work in combination with CG4733 in CI md neurons as a part of the PP2A complex to regulate dendritic arborization. Due to the phenotypic similarities between *mts* and *wdb* knockdown on md neurons, our data suggest that cell-type specific differences in CIV and CI neurons to PP2A loss of function may be due to varying target proteins in these neuronal subtypes. Previous comparative neurogenomic studies have highlighted differential mRNA content between CIV and CI md neurons, (Iyer *et al*, 2013a) and there is ample evidence for subtype-specific TF-mediated regulation of gene expression (Nanda *et al*, 2017). Our results reveal that PP2A contributes to cell-type specific dendritic diversity by promoting growth and branching in CIV neurons, while restricting dendritic branching in CI neurons.

Developmental time course analysis in CIV neurons revealed that PP2A is required for dendritic elaboration in late larval periods rather than establishment of dendritic arbor patterns. In CIV md neurons, we found no discernible phenotypic differences between controls and *mts-IR* neurons at either 24h or 48h AEL. However, at 72h AEL, a significant difference in total dendritic length appears between control and *mts-IR* neurons. The gap between control and *mts-IR* widens at 96h AEL, as *mts-IR* neurons actually show a significant reduction in field coverage between the two time points. These data suggest that PP2A is critical both for elaboration of dendritic arbors and sustainment of dendritic complexity.

In addition to the canonical regulatory subunits of PP2A, Mts and PP2A-29B also associate with Cka, the regulatory subunit of the STRIPAK complex. Cka is the *Drosophila* ortholog of the human Striatin protein (Hwang & Pallas, 2014). In *Drosophila,* the STRIPAK complex has been shown to positively regulate the Ras-Raf-Erk signaling pathway and negatively regulates the Hippo pathway (Hwang & Pallas, 2014; Sakuma & Chihara, 2017; Sakuma *et al*, 2016). Our study shows that loss of *Cka* severely disrupts dendritic morphology indicating that the STRIPAK complex regulates dendritic arborization in CIV md neurons. However, this regulation is independent of its interaction with the PP2A complex as the expression of *Cka* mutants unable to interact with the PP2A complex did not affect dendritic morphology. Collectively, our analyses show that while the PP2A and STRIPAK complexes are required for regulating dendritic morphology, they function independent of each other in these neurons.

### Regulatory interactions between PP2A and the TFs Cut and FoxO contribute to cell-type specific dendritic diversity

The TF *cut* was previously shown to be expressed in CIV neurons, but not CI neurons (Grueber *et al*, 2003). Additionally, Cut has previously been found to regulate the cytoskeleton in CIV neurons (Jinushi-Nakao *et al*, 2007; Das *et al*, 2017). Our data shows that *mts* overexpression in a *ct-IR* knockdown background partially rescued the dendritic defects caused by *ct* knockdown. This suggests that PP2A acts downstream of Cut in regulating dendritic morphology in CIV md neurons.

In CI neurons that do not express detectable levels of Cut, we examined the relationship between PP2A and FoxO. FoxO has been implicated in regulating morphology in md neurons (Sears & Broihier, 2016), and its phosphorylation state has been found to impact its subcellular localization as well as activity (Hu *et al*, 2019). Phosphorylated FoxO is localized in the cytoplasm which prevents its transcriptional activity thus promoting cellular growth (Zhang *et al*, 2011; Hu *et al*, 2019). IHC analysis shows that *mts* knockdown increases global FoxO levels in CI md neurons. In addition, the phenotypic defects due to *foxo* overexpression are similar to those observed for *mts* knockdown. Coupled together, these results suggest that PP2A and FoxO genetically interact with each other. This is further supported by our rescue analysis where the aberrant ectopic branching observed in *foxo-OE* is rescued by simultaneous overexpression of *mts.* Moreover, *foxo* overexpression is known to reverse microtubule polarity and also reduce Futsch levels in md neurons (Sears & Broihier, 2016), consistent with what we observed for *mts* knockdown. Taken together, this suggests that FoxO may be a regulatory target of PP2A in CI neurons.

### PP2A is required for cytoskeletal organization

Neurons require a properly formed cytoskeleton in order to achieve correct dendritic architecture. Through live imaging and IHC analysis we show that loss of *mts* leads to significant decrease of dendritic stable MTs in both CI and CIV neurons. Furthermore, time-lapse imaging of EB1 comets in CIV and CI neurons showed that *mts* knockdown reverses microtubule polarity in both dendrites and axons. While the reversal in MT polarity due to the knockdown of *mts* was previously reported (Rui *et al*, 2020), we observed a more robust effect of the loss of *mts* on EB1 polarity. This is possibly due to differences in the specific transgenic strain used for *mts* knockdown or to developmental stage, as our experiments were carried out at approximately 120h AEL, while a previous report (Rui *et al*, 2020) focused on studies at 96 h AEL. The reversal in EB1 comet polarity, however, was restricted to primary dendrites. Further, CIV neurons expressing *UAS-mts-IR* also showed a significant reduction in γ-tubulin along the dendritic arbor together with disruption of the microtubule lattice formed by the minus end protein Patronin. In CIV neurons, PP2A has been shown to regulate MT orientation by suppressing Klp10 protein (Rui *et al*, 2020), which has also been implicated in regulating MT polarity through its interaction with Patronin (Wang *et al*, 2019). Loss of γ-tubulin and Patronin, have both previously been shown to alter MT polarity in CIV neurons (Nguyen *et al*, 2014; Feng *et al*, 2019; Wang *et al*, 2019). Previous work suggested that Patronin and PP2A regulated MT polarity independently through their interactions with Klp10 (Rui *et al*, 2020; Wang *et al*, 2019); however, our finding that loss of *mts* resulted in reductions of *UAS-Patronin-GFP* signal indicates that PP2A may also work via Patronin to regulate MT polarity. Both hyperstabilization of MTs as well as overly dynamic MTs can lead to neuronal cell death (Feinstein & Wilson, 2005). Tubulin tdEOS imaging shows that *mts* knockdown leads to an increase in MT turnover rate, suggesting that PP2A is required to maintain the balance between dynamic and stabilized MT.

Like MT levels, F-actin levels in CIV neurons are also decreased due to *mts* knockdown, though neurons additionally display a shift in peak F-actin quantity. However, there are subtype-specific effects: F-actin peak intensities in CIV neurons shift proximally towards the soma when compared with controls, whereas F-actin peak intensities in CI neurons shift distally from the soma relative to controls. Further, there is an increase in F-actin levels in CI neurons compared to controls. This suggests that *mts* knockdown leads to a reorganization of the F-actin cytoskeleton. In CIV md neurons, F-actin is highly dynamic and forms actin ‘blobs’ that move along the dendrites and act as sites of new branch formation (Nithianandam & Chien, 2018). Hyperstabilization of actin leads to reduction in actin dynamics, including the abolishment of actin blobs, while the F-actin severing protein Twinstar (Tsr)/Cofilin regulates actin blob dynamics and dendritic branching (Nithianandam & Chien, 2018). Moreover, increased F-actin stabilization also leads to loss of neuronal growth which has been attributed to increased levels of phosphorylated cofilin (Cook *et al*, 2014). Further, studies in dendritic pruning have shown that PP2A disruption in CIV md neurons can decrease the levels of Tsr/Cofilin which could contribute to changes in actin dynamics by impairing F-actin disassembly (Wolterhoff *et al*, 2020). Photoconvertible LifeAct-tdEOS imaging revealed that *mts* knockdown leads to an increase in F-actin stability when compared to controls suggesting that PP2A is required to maintain actin dynamics. Combined, our data suggest that PP2A has distinct effects in CI and CIV neurons on cytoskeletal components which modulate subtype-specific dendritic architecture. While PP2A knockdown severely disrupts MTs in both CI and CIV neurons, loss of Mts has differential effects on F-actin in these neurons.

### PP2A is required for organelle trafficking and maintaining neuronal compartmentalization

Disruption of cytoskeletal organization leads to disruption of cytoskeletal based organelle trafficking (Das *et al*, 2021). Neurons have high demands of protein synthesis and respond rapidly to local cues through localization of organelles along the dendritic arbor, primarily the Golgi apparatus, endoplasmic reticulum, and mitochondria. The transport of these organelles is mediated by MTs and disruption of cytoskeletal components has been found to severely impact organelle trafficking (Das *et al*, 2021). Loss of *mts* significantly decreases the localization of mitochondria along the dendritic arbor in both CI and CIV md neurons. However, disruption of PP2A has a context-dependent effect on Golgi outpost localization in md neurons. In CIV neurons, the loss of *mts* severely disrupts Golgi outpost localization along the dendritic arbor and at branch points, corresponding to the severe disruption of gross dendritic morphology. However, in CI neurons *mts* disruption leads to increased Golgi outpost trafficking throughout the arbor, corresponding to an increase in dendritic complexity. Golgi outpost localization at branch points act as sites of microtubule nucleation (Ori-McKenney *et al*, 2012; Yalgin *et al*, 2015). The decrease in Golgi outpost trafficking along the CIV dendritic arbor may occur as a result of the severe reduction in the population of stable MTs due to *mts* knockdown; however, a decrease in branching may also result from the loss of Golgi outposts which serve as nodes for new branch extension. By contrast, in CI md neurons which show increased ectopic branching with *mts* knockdown, Golgi outpost trafficking is upregulated. Studies have demonstrated that F-actin is required for Golgi trafficking in granule neurons of the cerebellum (Venkatesh *et al*, 2020). Cytoskeletal analysis in CI neurons showed that while *mts* knockdown led to a significant reduction in MTs, F-actin levels were locally upregulated. Therefore, the differences in Golgi outpost localization between CI and CIV neurons may be explained by the differential effects of *mts* knockdown on F-actin, and CI neurons may exhibit higher Golgi outpost trafficking to branches due to the resultant local increases in F-actin rich branches.

In addition to the changes seen in the dendrites, disruption of the PP2A complex also led to Golgi outpost mislocalization in the axons. *Drosophila* neurons are highly polarized with Golgi outpost localization restricted predominantly to dendrites (Horton *et al*, 2005; Kelliher *et al*, 2018; Ye *et al*, 2007). Loss of *mts* led to the mislocalization of Golgi outposts into the proximal axonal segments. Further, this mislocalization was also associated with increased branching in the axon initial segment. PP2A disruption also led to mislocalization of the dendritic marker DenMark into the axon along with the increased expression of the axonal synaptic marker, Synaptotagmin into the dendrites. Our results therefore suggest that PP2A is required for maintaining proper neuronal compartmentalization. These results are consistent with a recent study which showed that PP2A mutations led to mislocalization of the dendritic marker Nod-β-gal into the axons along with the mislocalization of the axonal marker Kin-β-gal into the dendrites of CIV md neurons (Rui *et al*, 2020). Loss of Patronin also leads to aberrant localization of dendritic markers in the axons and vice versa (Wang *et al*, 2019). As our data show Patronin disruption in *mts* mutations, PP2A may be required for neuronal compartmentalization through its regulation of Patronin.

### β-tubulin is a target of PP2A in md neurons

Based upon our results demonstrating roles for PP2A in dendritic cytoskeletal organization and especially on MTs, we decided to investigate whether PP2A directly regulates MT subunits. Microtubules are highly dynamic structures composed of heterodimers of α- and β-tubulin molecules (Khan & Ludueña, 1996; Janke & Magiera, 2020). The serine 172 (S172) residue in β-tubulin is highly conserved and undergoes phosphorylation that influences its dynamic state (Wloga & Gaertig, 2010). Phosphorylation of β-tubulin S172 by DYRK1A in *Drosophila* md neurons reduces dendritic arborization while its phosphorylation by CDK1 prevents tubulin polymerization (Janke & Magiera, 2020). Knockdown of *mts* leads to increased levels of phosphorylated β-tubulin S172 suggesting that PP2A may be required for β-tubulin dephosphorylation. This, along with phenotypic analysis in both CI and CIV md neurons indicated that β-tubulin 85D may be a putative regulatory target of PP2A in these two md neuron subtypes. Overexpression of *β-tubulin 85D* in *mts-IR* background partially rescued the severely disrupted phenotype arising from the knockdown of *mts.* We speculate that an increase in β-tubulin levels in this sensitized background could outcompete the available kinases that phosphorylate tubulin, creating pools of unphosphorylated β-tubulin. These unphosphorylated β-tubulin could then be incorporated into microtubules and rescue some of the phenotypic defects caused by the knockdown of *mts.* Supporting this theory, we found that expressing the constitutively active phospho-resistant β-tubulin85D did not affect dendritic morphology. However, expression of the phosphomimetic form of β-tubulin85D led to dendritic hypotrophy. Collectively, our data are supportive of a putative regulatory interaction between PP2A and β-tubulin 85D in md neuron dendritic development.

The present study demonstrates the requirement of PP2A in regulating cell-type specific dendritic morphology in md neurons. In simpler CI neurons, PP2A restricts dendritic growth, but in more complex CIV neurons, PP2A promotes dendritogenesis. Our study also provides mechanistic insights into how Cut regulates morphogenesis through the PP2A complex in CIV neurons, while PP2A interacts with the TF FoxO to regulate CI architecture. At the cytoskeletal level, Mts functions to stabilize MTs and organize F-actin. Our study also demonstrates that Mts is required to maintain MT polarity. While previous studies have demonstrated that PP2A maintains MT polarity by modulating Klp10, our data also show that PP2A is required for proper γ-tubulin localization and Patronin organization, both of which are required to maintain MT polarity. PP2A may partially regulate MT stability and organization through direct targeting of β-tubulin85D, which appears to affect CI and CIV arbors equally. Combined, this study provides novel insights into the developmental role of PP2A in regulating dendritic morphology through multivarious interactions with the cytoskeleton, both direct and indirect.

## Figure Legends

**Figure S1: *Regulatory subunits of PP2A have mild effects on dendritic morphology*: (A, A’)** Immunohistochemical analysis shows that Mts is expressed in md neurons. **(B, C)** Sholl quantitative morphometric analyses. Representative images of CIV neurons in **(D)** control, **(E)** *wdb-IR* **(F)** *wrd-IR*, **(G)** *tws-IR* **(H)** *CG4733-IR*. **(I, J)** Quantitative morphometric analyses. Representative images of CI neurons in **(K)** control, **(L)** *wrd-IR*, **(M)** *tws-IR* **(N)** *CG4733-IR*. **(O, P)** Quantitative morphometric analyses. Statistical tests performed: **(B, C, J, O)** Kruskal-Wallis with Dunn’s multiple comparisons test (n = 10-16 per genotype), **(I, P)** One-way ANOVA with Sidak’s multiple comparisons test (n = 10-16 per genotype). ***=p≤0.001, **=p≤0.01, *=p≤0.05. For detailed genotypes see Table S1 and for detailed statistics see Table S2. Scale bar = 5 µm for **(A-A’)**, 200 µm for **(D)**, and 100 µm for **(K).**

**Figure S2: *PP2A is required for MT stabilization:*** Representative images of immunohistochemical analyses showing the expression of Futsch in md neurons in **(A)** control and **(A’)** *mts-IR* animals. **(B)** Quantitative analysis of the mean fluorescence intensities of Futsch signaling normalized to area in md neurons. Representative images of immunohistochemical analysis showing the expression of acetylated tubulin in CIV neurons in **(C, C’)** control and **(D, D’)** *mts-IR* animals. **(E)** Quantitative analysis of the mean fluorescence intensities of acetylated tubulin normalized to area. Statistical tests performed: **(B)** Two-way ANOVA with Dunnett’s multiple comparisons test (n = 14-18 per genotype), **(E)** Student’s t-test (n = 11-16 per genotype). ***=p≤0.001. For detailed genotypes see Table S1 and for detailed statistics see Table S2. Scale bar = 50 µm for **(A)** and 5 µm for **(C-C’)**.

**Figure S3: *PP2A knockdown disrupts MT polarity in both axons and dendrites of md neurons but not dendritic terminals:*** Kymographs showing the EB1::GFP comet trajectories in dendrites of CI neurons in **(A)** control and **(B)** *mts-IR*. **(C, D)** Knockdown of *mts* in CI or CIV md neurons does not affect MT polarity at dendritic terminals. Representative kymographs of EB1::GFP comets in axons of CIV and CI neurons in **(E, G)** control and **(F, H)** *mts-IR* respectively. **(I, J)** Compared to controls, knockdown of *mts* reverses MT polarity in CI axons along with an increase in the number of comets. Statistical tests performed: **(C, D, I)** Two-way ANOVA with Sidak’s multiple comparison test (n = 28-129 comets per genotype), **(J)** Mann-Whitney U test (n = 8-12 per genotype). ***=p≤0.001. For detailed genotypes see Table S1 and for detailed statistics see Table S2.

**Figure S4: *mts knockdown disrupts neural compartment specification:* (A-D)** Representative images showing the distribution of DenMark and Synaptotagmin signal intensities in proximal axons and distal dendrites in CI neurons in **(A, C)** control and **(B, D)** *mts-IR*. **(E)** Quantitative analysis of the mean fluorescence intensities of Synaptotagmin in dendrites **(F)** Quantitative analysis of the mean fluorescence intensities of DenMark in axons. Statistical tests performed: Mann-Whitney U test **=p≤0.01 (n = 12-15 per genotype) Scale bar = 50µm.For detailed genotypes see Table S1 and for detailed statistics see Table S2.

**Figure S5: *Phenotypic analysis of β-tubulins:*** Representative images of CIV md neurons in **(A)** control,

**(A)** *β-tubulin56D-IR*, **(C)** *β-tubulin60D-IR*, **(D)** *β-tubulin97EF-IR*, **(E)** *CG32396-IR*. **(F, G)** Quantitative morphometric analyses. Statistical tests performed: One-way ANOVA with Dunnet’s multiple comparison test (n = 11-16 per genotype). ***=p≤0.001, **=p≤0.01. For detailed genotypes see Table S1 and for detailed statistics see Table S2. Scale bar = 200 µm.

**Figure S6: *Expression of phosphomimetic form of β-tubulin85D disrupts dendritic morphology:* (A)** Schematic representation showing the amino acids mutated in *β*-tubulin85D to generate phospho-resistant and phosphomimetic forms of the protein. Representative images of CIV neurons in **(B)** control, **(C)** *β-tubulin85D-S172A_T219A*, and **(D)** *β-tubulin85D-S172E_T219E*. **(E, F)** Quantitative morphometric analyses. Statistical tests performed: **(E, F)** One-way ANOVA with Dunnett’s multiple comparison test (n = 7-11 per genotype). ***=p≤0.001, **=p≤0.01. For detailed genotypes see Table S1 and for detailed statistics see Table S2. Scale bar = 200 µm.

## Notes

### Competing Interest Statement

The authors have declared no competing interest.

