## Supplementary figures and images for "PP2A phosphatase regulates cell-type specific cytoskeletal organization to drive dendritic diversification"

### Supplemental Figures

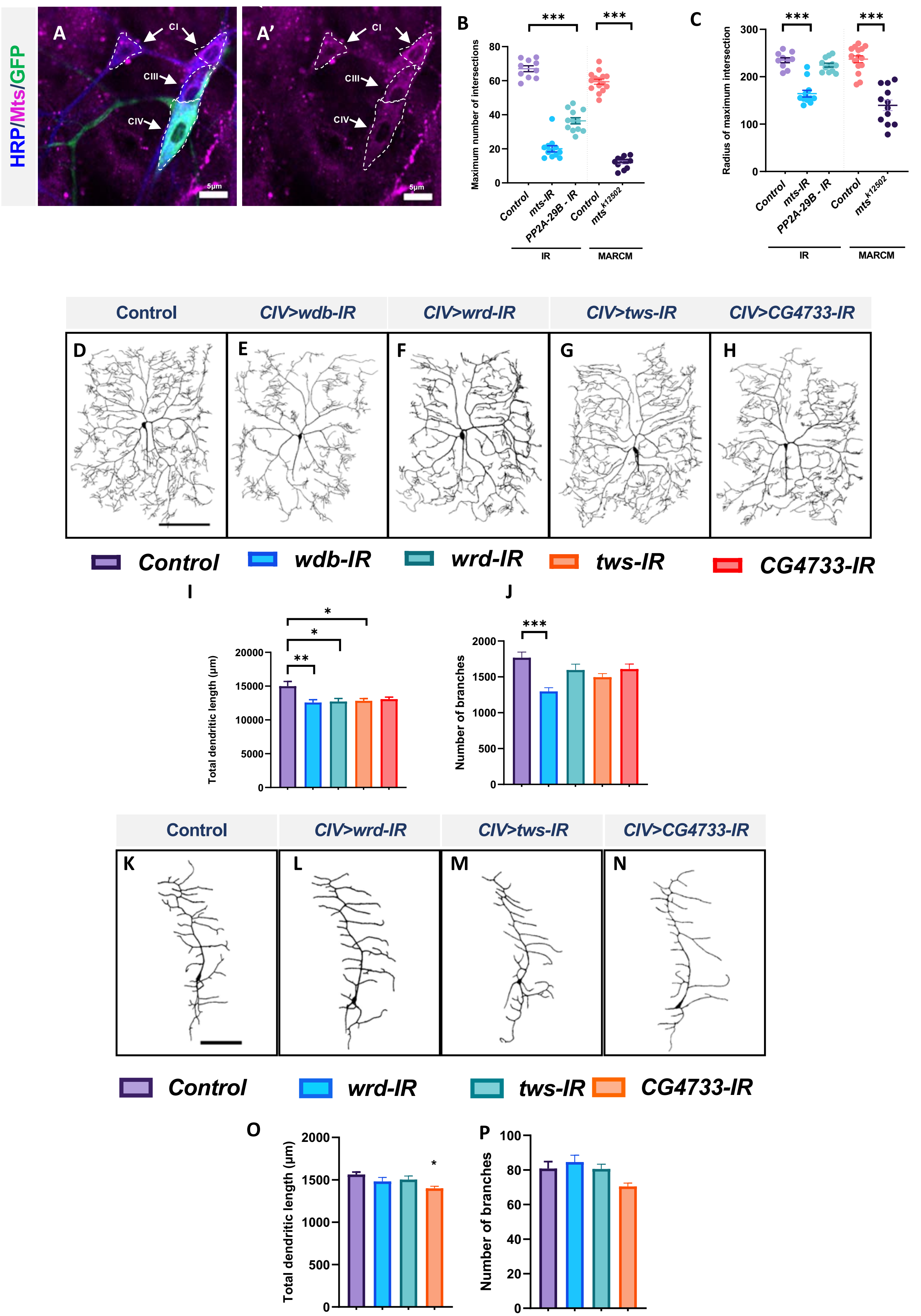

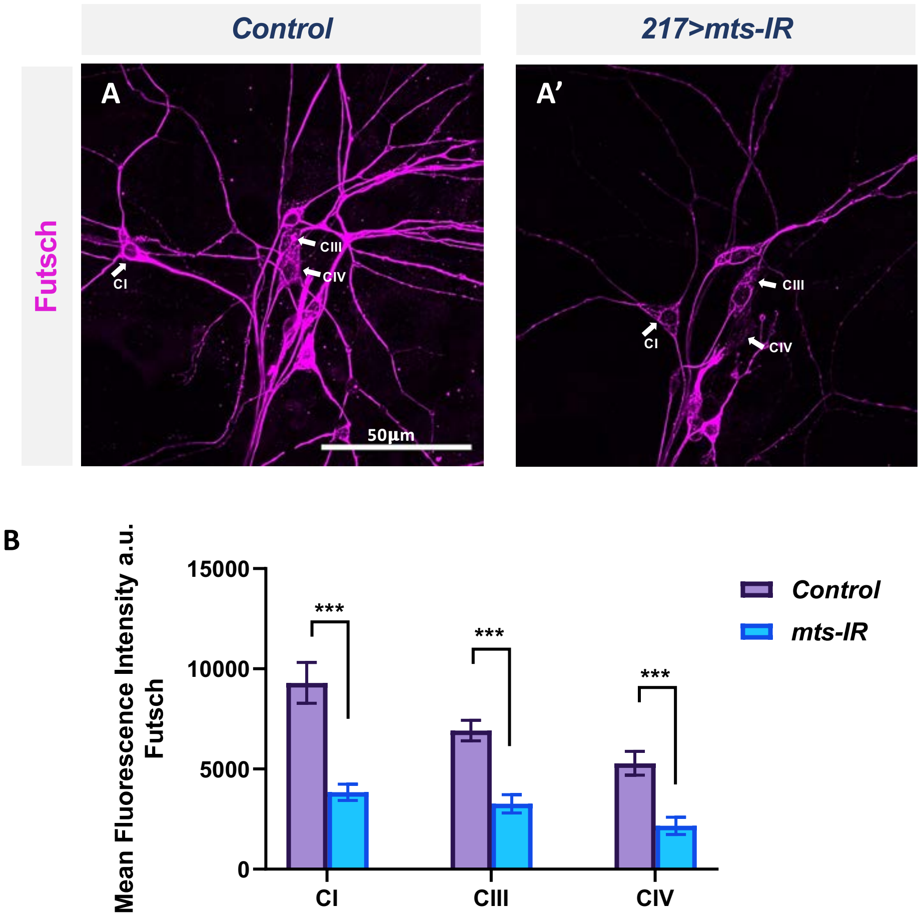

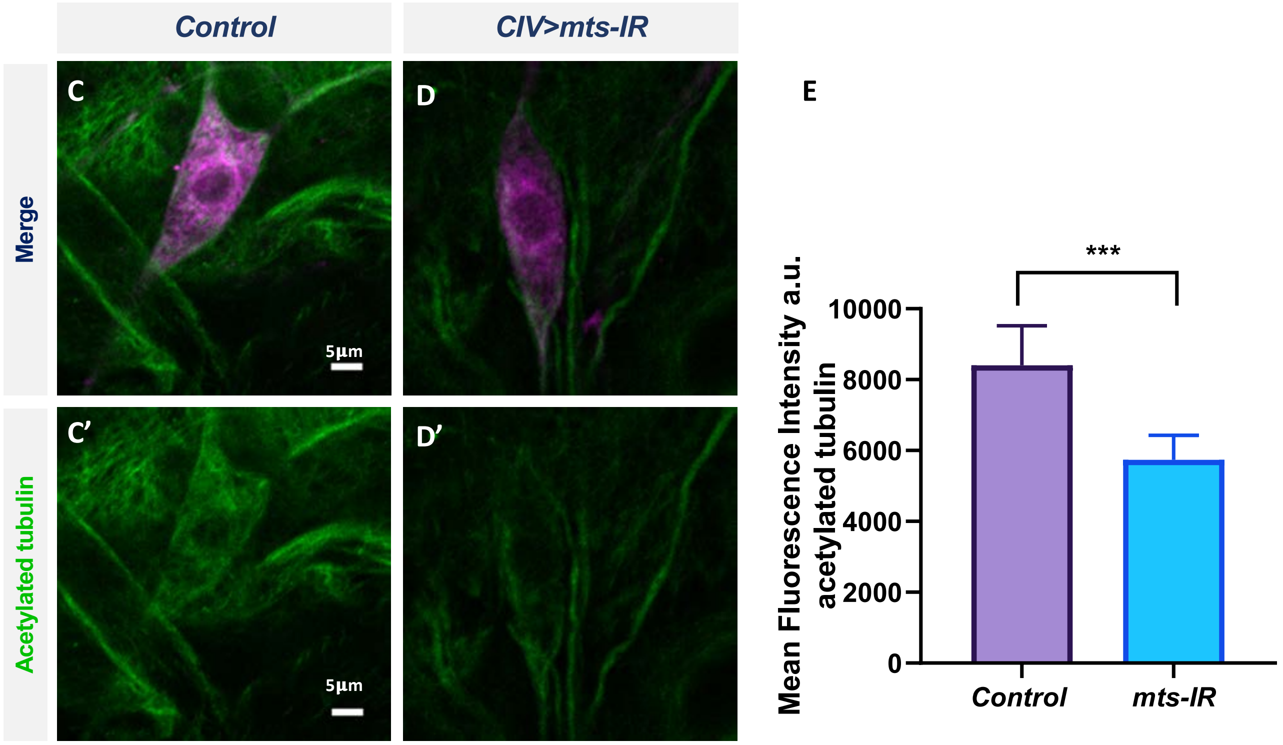

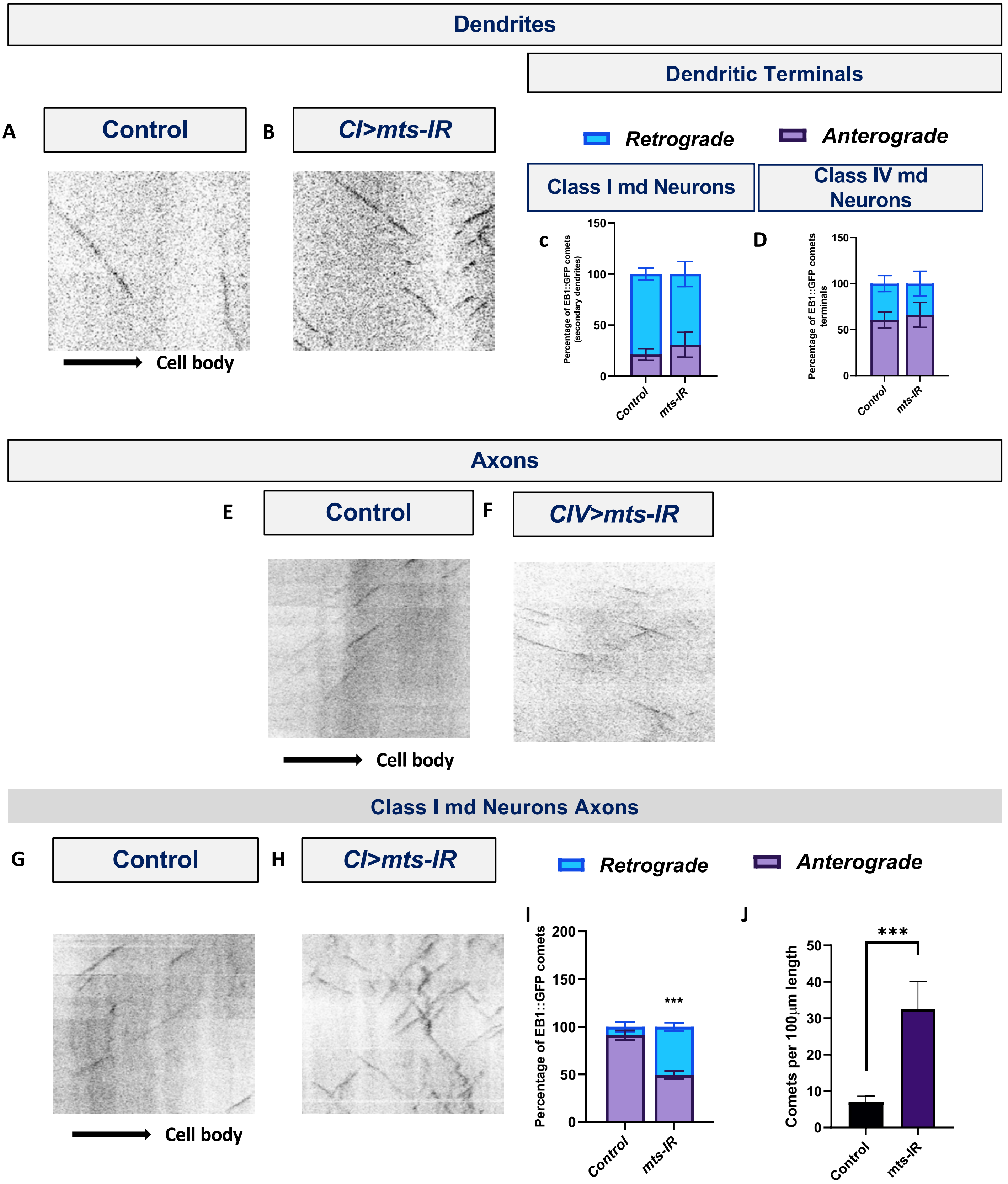

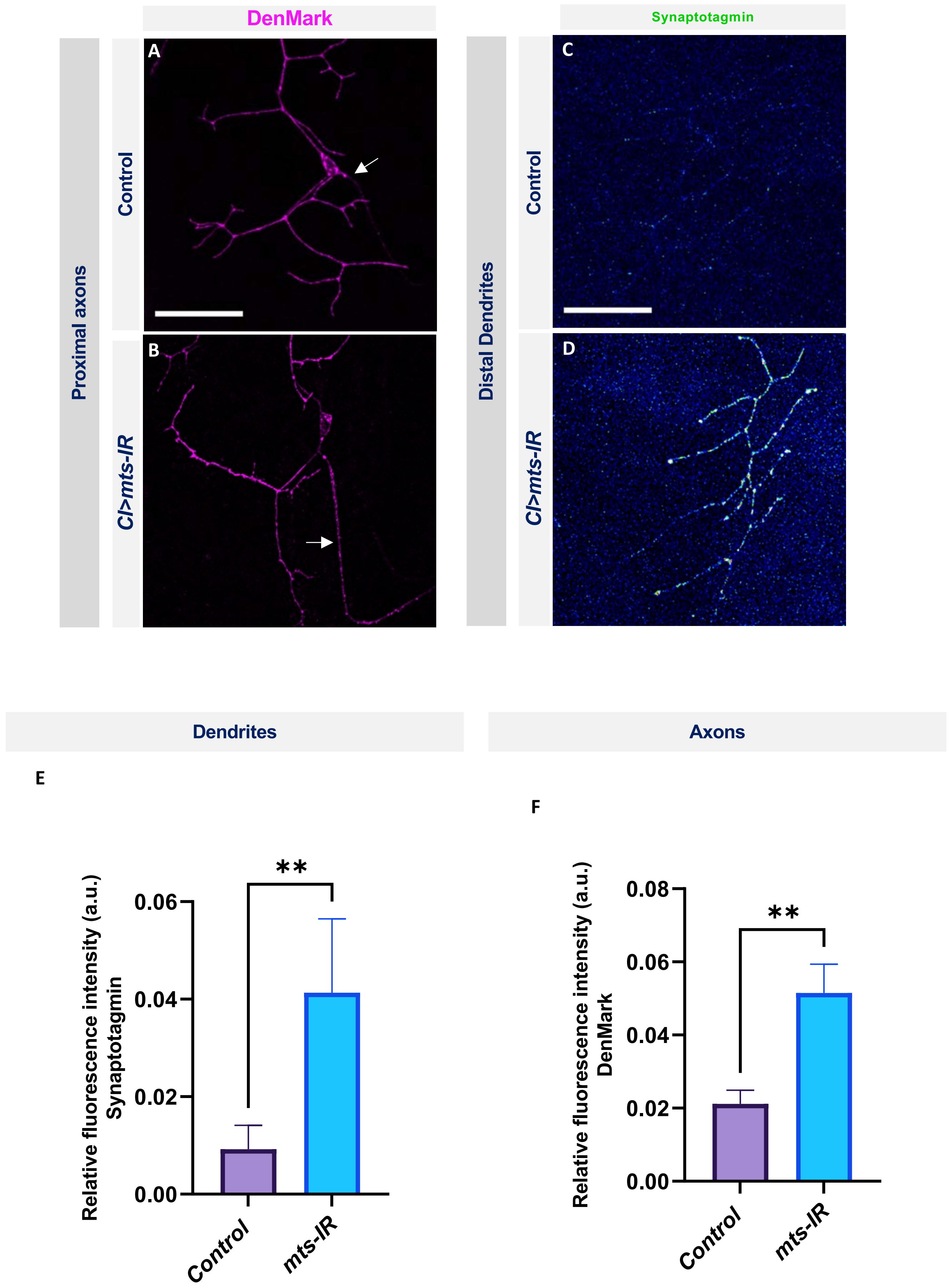

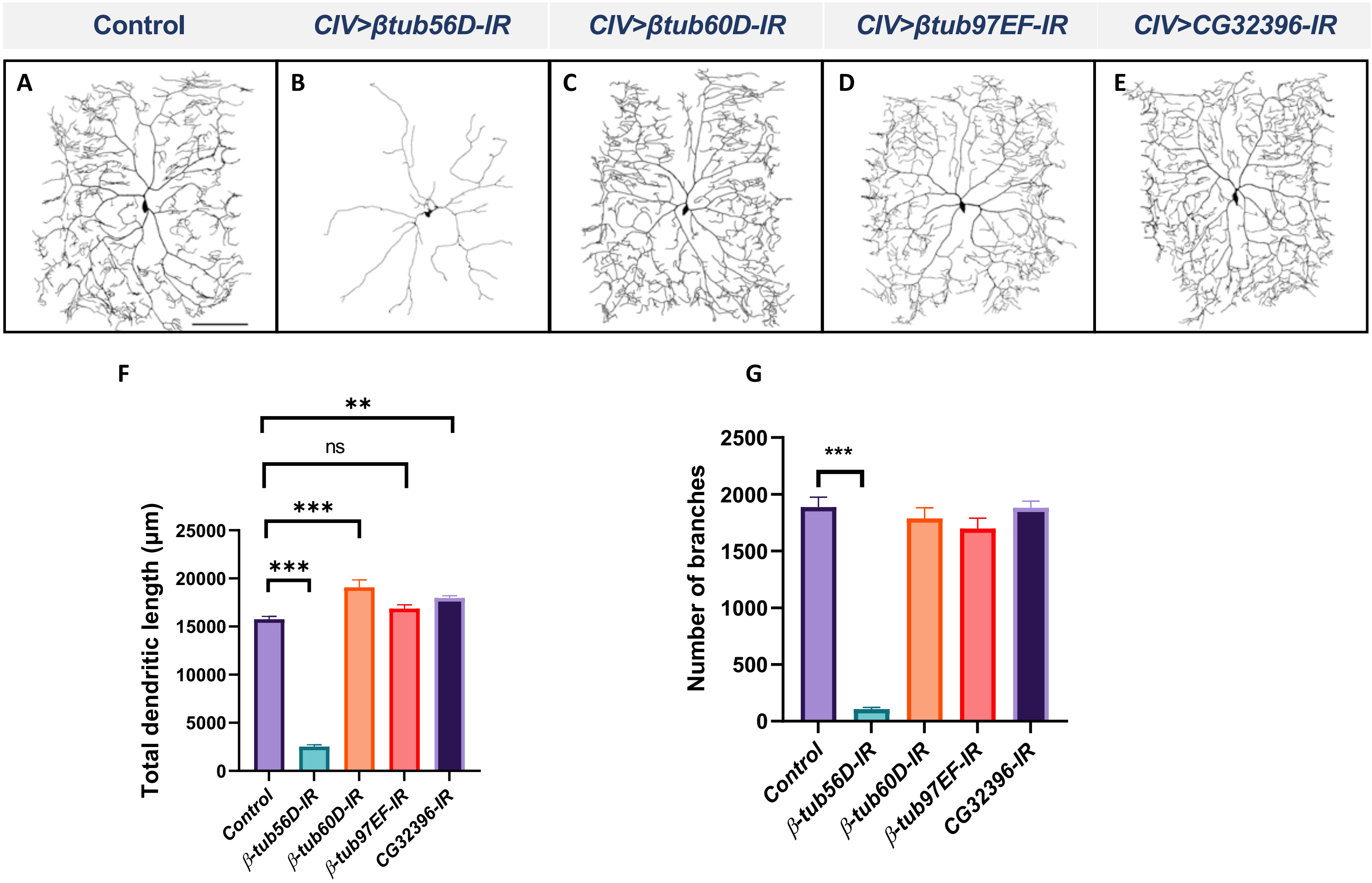

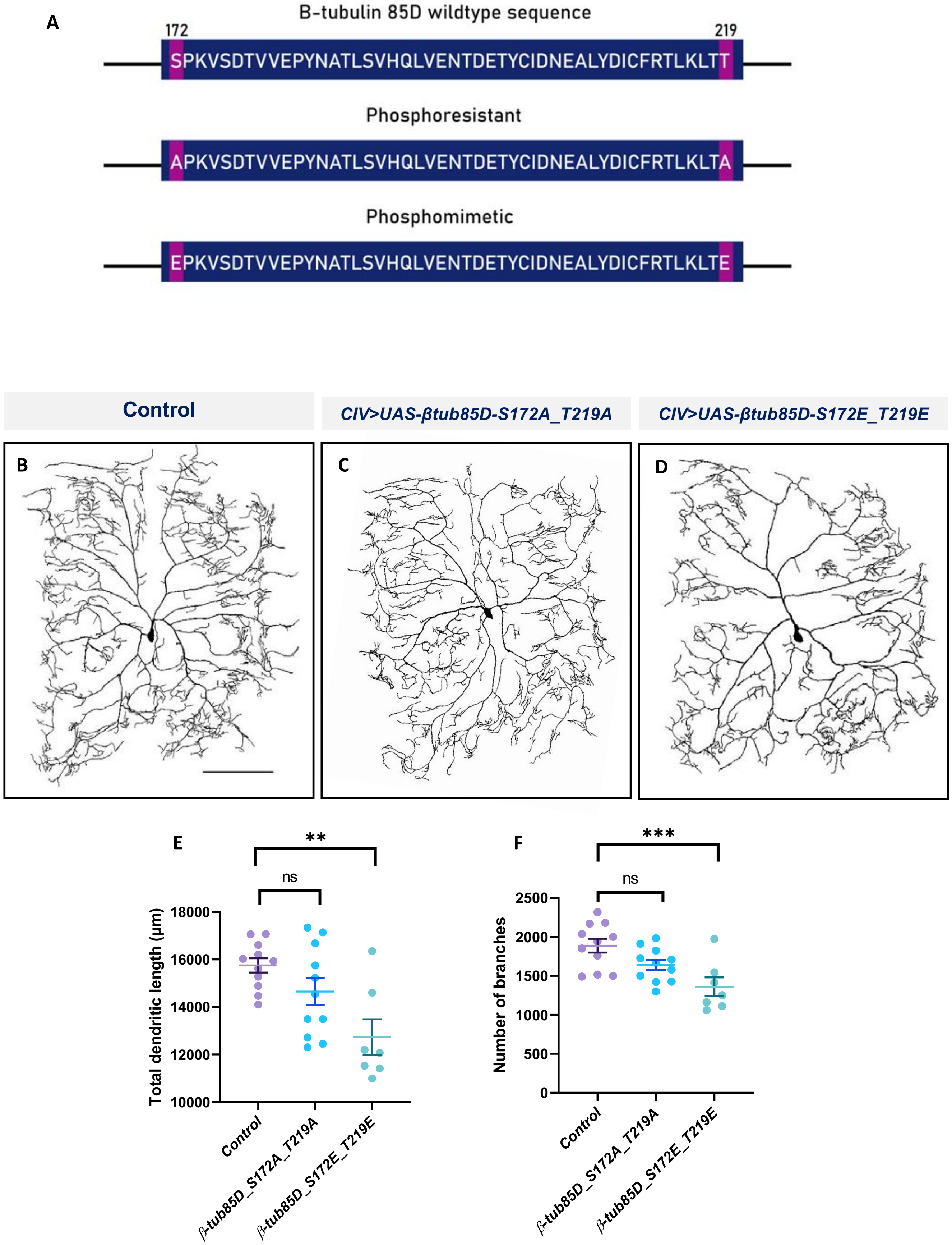
