## Supplementary material for "PP2A phosphatase regulates cell-type specific cytoskeletal organization to drive dendritic diversification": Table S1

| FIGURE | GENOTYPE |
|---|---|
| 1A | GAL4$^{477}$,UAS-mCD8::GFP/+;GAL4$^{ppk1.9}$,UAS-mCD8::GFP/+ |
| 1B | GAL4$^{477}$,UAS-mCD8::GFP/UAS-mts-IR;GAL4$^{ppk1.9}$,UAS-mCD8::GFP/+ |
| 1C | GAL4$^{5\text{-}40}$,UAS-Venus:pmSOP-FLP$^{\#42}$FRT$^{40A}$mts$^{k12502}$ |
| 1D | GAL4$^{477}$,UAS-mCD8::GFP/UAS-PP2A-29B-IR;GAL4$^{ppk1.9}$,UAS-mCD8::GFP/+ |
| 1N | GAL4$^{221}$,UAS-mCD8::GFP/+ |
| 1O | UAS-mts-IR/+;GAL4$^{221}$,UAS-mCD8::GFP/+ |
| 1P | GAL4$^{5\text{-}40}$,UAS-Venus:pmSOP-FLP$^{\#42}$FRT$^{40A}$mts$^{k12502}$ |
| 1Q | UAS-PP2A-29B-IR/+;GAL4$^{221}$,UAS-mCD8::GFP/+ |
| 1R | GAL4$^{221}$,UAS-mCD8::GFP/UAS-wd-IR |
| 1S | GAL4$^{109(2)80}$,UAS-mCD8::GFP SOP-FLP$^{\#73}$;FRT$^{82B}$wdb$^{14}$ |
| 2A | GAL4$^{477}$,UAS-mCD8::GFP/+;GAL4$^{ppk1.9}$,UAS-mCD8::GFP/+ |
| 2B | GAL4$^{477}$,UAS-mCD8::GFP/UAS-mts;GAL4$^{ppk1.9}$,UAS-mCD8::GFP/+ |
| 2C | GAL4$^{477}$,UAS-mCD8::GFP/UAS-PP2A-29B;GAL4$^{ppk1.9}$,UAS-mCD8::GFP/+ |
| 2D | GAL4$^{477}$,UAS-mCD8::GFP/UAS-wdb;GAL4$^{ppk1.9}$,UAS-mCD8::GFP/+ |
| 2H | GAL4$^{477}$,UAS-mCD8::GFP/+;GAL4$^{ppk1.9}$,UAS-mCD8::GFP/+ |
| 2I | GAL4$^{477}$,UAS-mCD8::GFP/UAS-cka-IR;GAL4$^{ppk1.9}$,UAS-mCD8::GFP/+ |
| 2J | GAL4$^{477}$,UAS-mCD8::GFP/+;GAL4$^{ppk1.9}$,UAS-mCD8::GFP/UAS-cka-EGFP |
| 2K | GAL4$^{477}$,UAS-mCD8::GFP/UAS-BFP-cka$^{\Delta PP2A}$;GAL4$^{ppk1.9}$,UAS-mCD8::GFP/+ |
| 3A-A’’’ | nos-GAL4/+;ppk-hCD4-tdTOMATO/+;ppk-GAL4/+ |
| 3 B-B’’’ | nos-GAL4/+;UAS-mts-IR/ppk-hCD4-tdTOMATO;ppl-GAL4/+ |
| 4A | GAL4$^{477}$,UAS-mCD8::GFP/+;UAS-CD4-tdTOMATO/+ |
| 4B | GAL4$^{477}$,UAS-mCD8::GFP/+;UAS-ct-IR/UAS-CD4-tdTOMATO |

| | |
|---|---|
| **4C** | *GAL4$^{477}$,UAS-mCD8::GFP/UAS-mts;UAS-ct-IR/+* |
| **5A-A”** | *UAS-GMA/+;GAL4$^{477}$,UAS-mCherry::JUPITER/+* |
| **5B-B”** | *UAS-GMA/+;GAL4$^{477}$,UAS-mCherry::JUPITER/UAS-mts-IR* |
| **5L-L”** | *UAS-GMA/+; GAL4$^{221}$,UAS-mCherry::JUPITER/+* |
| **5M-M”** | *UAS-GMA/+; UAS-mts-IR/+;GAL4$^{221}$,UAS-mCherry::JUPITER/+* |
| **6A-A’’’** | *GAL4$^{477}$/+;UAS-alphaTUB84BtdEOS/+* |
| **6B-B’’’** | *GAL4$^{477}$/UAS-mts-IR;UAS-alphaTUB84BtdEOS/+* |
| **6D-D’’’** | *+/+;GAL4$^{ppk1.9}$/LifeAct::tdEOS* |
| **6E-E’’’** | *UAS-mts-IR/+;GAL4$^{ppk1.9}$/ LifeAct::tdEOS* |
| **7A** | *GAL4$^{ppk}$/+;ppk-EB1::GFP/+* |
| **7B** | *GAL4$^{ppk}$/UAS-mts-IR;ppk-EB1::GFP/+* |
| **8A-A”** | *UAS-γ-tubulin23C-GFP/+;GAL4$^{ppk1.9}$,UAS-mCD8::GFP/+* |
| **8B-B”** | *UAS-γ-tubulin23C-GFP/UAS-mts-IR;GAL4$^{ppk1.9}$,UAS-mCD8::GFP/+* |
| **8D** | *UAS-Patronin-GFP/+; GAL4$^{ppk1.9}$,UAS-mCD8::RFP/+* |
| **8E** | *UAS-Patronin-GFP/UAS-mts-IR; GAL4$^{ppk1.9}$,UAS-mCD8::RFP/+* |
| **8G, G’** | *GAL4$^{477}$,UAS-mitoGFP.AP/+;ppk-hCD4-tdTOMATO/+* |
| **8H, H’** | *GAL4$^{477}$,UAS-mitoGFP.AP/UAS-mts-IR;ppk-hCD4-tdTOMATO/+* |
| **8I, I’** | *UAS-mitoGFP.AP/+;GAL4$^{221}$,UAS-mCD8::RFP/+* |
| **8J, J’** | *UAS-mitoGFP.AP/UAS-mts-IR;GAL4$^{221}$,UAS-mCD8::RFP/+* |
| **9A, A’** | *GAL4$^{477}$,UAS-MANII-eGFP/+;ppk-hCD4-tdTOMATO/+* |
| **9B, B’** | *GAL4$^{477}$,UAS-MANII-eGFP/UAS-mts-IR;ppk-hCD4-tdTOMATO/+* |
| **9C, C’** | *UAS-MANII-eGFP/+;GAL4$^{221}$,UAS-mCD8::RFP/+* |
| **9D, D’** | *UAS-MANII-eGFP/UAS-mts-IR;GAL4$^{221}$,UAS-mCD8::RFP/+* |

| | |
|---|---|
| **9I** | *GAL4$^{477}$/+;ppk-hCD4-tdTOMATO/+* |
| **9J** | *GAL4$^{477}$/UAS-mts-IR;ppk-hCD4-tdTOMATO/+* |
| **9K** | *GAL4$^{477}$,UAS-mCD8::GFP/+;GAL4$^{ppk1.9}$,UAS-mCD8::GFP/+* |
| **9L** | *GAL4$^{477}$,UAS-mCD8::GFP/UAS-mts-IR;GAL4$^{ppk1.9}$,UAS-mCD8::GFP/+* |
| **9N, N'** | *GAL4$^{477}$,UAS-MANII-eGFP/+;ppk-hCD4-tdTOMATO/+* |
| **9O, O'** | *GAL4$^{477}$,UAS-MANII-eGFP/UAS-mts-IR;ppk-hCD4-tdTOMATO/+* |
| **9Q, Q'** | *UAS-MANII-eGFP/+;GAL4$^{221}$,UAS-mCD8::RFP/+* |
| **9R, R'** | *UAS-MANII-eGFP/UAS-mts-IR;GAL4$^{221}$,UAS-mCD8::RFP/+* |
| **10A** | *GAL4$^{221}$,UAS-mCD8::GFP/+* |
| **10B** | *UAS-foxo.P/+;GAL4$^{221}$,UAS-mCD8::GFP/+* |
| **10C, C'** | *GAL4$^{221}$,UAS-mCD8::GFP/+* |
| **10D, D'** | *UAS-mts-IR/+;GAL4$^{221}$,UAS-mCD8::GFP/+* |
| **10I** | *GAL4$^{221}$,UAS-mCD8::GFP/+* |
| **10J** | *UAS-mts-IR/+;GAL4$^{221}$,UAS-mCD8::GFP/+* |
| **10K** | *UAS-mts/UAS-foxo.P; GAL4$^{221}$,UAS-mCD8::GFP/+* |
| **11A** | *GAL4$^{477}$,UAS-mCD8::GFP/+;GAL4$^{ppk1.9}$,UAS-mCD8::GFP/+* |
| **11B** | *GAL4$^{477}$,UAS-mCD8::GFP/UAS-β-tubulin85D-IR;GAL4$^{ppk1.9}$,UAS-mCD8::GFP/+* |
| **11C** | *GAL4$^{477}$,UAS-mCD8::GFP/+;GAL4$^{ppk1.9}$,UAS-mCD8::GFP/UAS-β-tubulin85D* |
| **11F** | *GAL4$^{ppk1.9}$,UAS-mCD::GFP/UAS-hCD4-tdTOMATO* |
| **11G** | *UAS-mts-IR/+;GAL4$^{ppk1.9}$,UAS-mCD::GFP/UAS-hCD4-tdTOMATO* |
| **11H** | *UAS-mts-IR/+;GAL4$^{ppk1.9}$,UAS-mCD::GFP/ UAS-β-tubulin85D* |
| **11P,P'** | *GAL4$^{ppk1.9}$,UAS-mCD::GFP/+* |

| | |
|---|---|
| **11Q,Q'** | *UAS-mts-IR/+;GAL4$^{ppk1.9}$,UAS-mCD::GFP/+* |
| **12A** | *GAL4$^{221}$,UAS-mCD8::GFP/+* |
| **12B** | *GAL4$^{221}$,UAS-mCD8::GFP/ UAS-β-tubulin85D-IR* |
| **12G** | *GAL4$^{221}$,UAS-mCD8::GFP/+* |
| **12H** | *UAS-mts-IR/+;GAL4$^{221}$,UAS-mCD8::GFP/+* |
| **12I** | *UAS-mts-IR/+;GAL4$^{221}$,UAS-mCD8::GFP/ UAS-β-tubulin85D* |
| **S1A, A'** | *GAL4$^{ppk1.9}$,UAS-mCD8::GFP/+* |
| **S1D** | *GAL4$^{477}$,UAS-mCD8::GFP/+;GAL4$^{ppk1.9}$,UAS-mCD8::GFP/+* |
| **S1E** | *GAL4$^{477}$,UAS-mCD8::GFP/+;GAL4$^{ppk1.9}$,UAS-mCD8::GFP/UAS-wdb-IR* |
| **S1F** | *GAL4$^{477}$,UAS-mCD8::GFP/UAS-wrd-IR;GAL4$^{ppk1.9}$,UAS-mCD8::GFP/+* |
| **S1G** | *GAL4$^{477}$,UAS-mCD8::GFP/ UAS-tws-IR;GAL4$^{ppk1.9}$,UAS-mCD8::GFP/+* |
| **S1H** | *GAL4$^{477}$,UAS-mCD8::GFP/UAS-CG4733-IR;GAL4$^{ppk1.9}$,UAS-mCD8::GFP/+* |
| **S1K** | *GAL4$^{221}$,UAS-mCD8::GFP/+* |
| **S1L** | *UAS-wrd-IR/+;GAL4$^{221}$,UAS-mCD8::GFP/+* |
| **S1M** | *UAS-tws-IR /+;GAL4$^{221}$,UAS-mCD8::GFP/+* |
| **S1N** | *UAS-CG4733-IR/+;GAL4$^{221}$,UAS-mCD8::GFP/+* |
| **S2A** | *GAL4$^{217}$,UAS-mCD8::GFP/+* |
| **S2A'** | *GAL4$^{217}$,UAS-mCD8::GFP/UAS-mts-IR* |
| **S2C, C'** | *GAL4$^{ppk1.9}$,UAS-mCD8::RFP/+* |
| **S2D, D'** | *UAS-mts-IR/+;GAL4$^{ppk1.9}$,UAS-mCD8::RFP/+* |
| **S3A, G** | *GAL4$^{221}$,UAS-EB1::GFP/+* |
| **S3B, H** | *UAS-mts-IR/+;GAL4$^{221}$,UAS-EB1::GFP/+* |
| **S3E** | *GAL4$^{ppk}$/+;ppk-EB1::GFP/+* |

| | |
|---|---|
| **S3F** | *$GAL4^{ppk}$/UAS-mts-IR;ppk-EB1::GFP/+* |
| **S4A, C** | *UAS-DenMark,UAS-syn.eGFP/+;$GAL4^{221}$/+* |
| **S4B, D** | *UAS-DenMark,UAS-syn.eGFP/UAS-mts-IR;$GAL4^{221}$/+* |
| **S5A** | *$GAL4^{477}$,UAS-mCD8::GFP/+;$GAL4^{ppk1.9}$,UAS-mCD8::GFP/UAS-Luc-IR* |
| **S5B** | *$GAL4^{477}$,UAS-mCD8::GFP/UAS-β-tubulin56D-IR;$GAL4^{ppk1.9}$,UAS-mCD8::GFP/+* |
| **S5C** | *$GAL4^{477}$,UAS-mCD8::GFP/UAS-β-tubulin60D-IR;$GAL4^{ppk1.9}$,UAS-mCD8::GFP/+* |
| **S5D** | *$GAL4^{477}$,UAS-mCD8::GFP/UAS-β-tubulin97EF-IR;$GAL4^{ppk1.9}$,UAS-mCD8::GFP/+* |
| **S5E** | *$GAL4^{477}$,UAS-mCD8::GFP/UAS-CG32396-IR;$GAL4^{ppk1.9}$,UAS-mCD8::GFP/+* |
| **S6B** | *$GAL4^{477}$,UAS-mCD8::GFP/+;$GAL4^{ppk1.9}$,UAS-mCD8::GFP/ UAS-Luc-IR* |
| **S6C** | *$GAL4^{477}$,UAS-mCD8::GFP/+;$GAL4^{ppk1.9}$,UAS-mCD8::GFP/UAS-β-tubulin85D-S172A_T219A* |
| **S6D** | *$GAL4^{477}$,UAS-mCD8::GFP/+;$GAL4^{ppk1.9}$,UAS-mCD8::GFP/UAS-β-tubulin85D-S172E_T219E* |
