## Supplementary material for "PP2A phosphatase regulates cell-type specific cytoskeletal organization to drive dendritic diversification": Table S2

| Figure/Label | Passed Shapiro-Wilk Normality Test | Statistical test used | p-value | Number of neurons (N) |
|---|---|---|---|---|
| **Fig 1E** | | **One-way ANOVA with Sidak's multiple comparison test** | | |
| Control | Yes | | | 10 |
| *mts-IR* | Yes | | <0.0001 | 12 |
| *PP2A-29B-IR* | Yes | | <0.0001 | 12 |
| Control-MARCM | Yes | | | 15 |
| $mts^{k12502}$ | Yes | | <0.0001 | 12 |
| **Fig 1F** | | **One-way ANOVA with Sidak's multiple comparison test** | | |
| Control | Yes | | | 10 |
| *mts-IR* | Yes | | <0.0001 | 12 |
| *PP2A-29B-IR* | Yes | | <0.0001 | 12 |
| Control-MARCM | Yes | | | 15 |
| $mts^{k12502}$ | Yes | | <0.0001 | 12 |
| **Fig 1G** | | **One-way ANOVA with Sidak's multiple comparisons** | | |
| Control | Yes | | | 10 |
| *mts-IR* | Yes | | <0.0001 | 12 |
| *PP2A-29B-IR* | Yes | | <0.0001 | 12 |
| Control-MARCM | Yes | | <0.0001 | 15 |
| $mts^{k12502}$ | Yes | | <0.0001 | 12 |
| **Fig 1H** | | **Kruskal-Wallis with Dunn's multiple comparison test** | | |
| Control | No | | | 10 |
| *mts-IR* | No | | <0.0001 | 12 |
| *PP2A-29B-IR* | No | | <0.0001 | 12 |
| **Fig 1I** | | **Kruskal-Wallis with Dunn's multiple comparison test** | | |
| Control-MARCM | No | | | 15 |
| $mts^{k12502}$ | No | | <0.0001 | 12 |
| **Fig 1J 1$^{st}$ order** | | **Two-way ANOVA with Dunnett's test for multiple comparison** | | |
| Control | | | | 10 |
| *mts-IR* | | | <0.0001 | 12 |

| | | | | |
|---|---|---|---|---|
| *PP2A-29B-IR* | | | <0.0001 | 12 |
| **Fig 1J 2$^{nd}$order** | | **Two-way ANOVA with Dunnett's test for multiple comparison** | | |
| Control | | | | 10 |
| *mts-IR* | | | <0.0001 | 12 |
| *PP2A-29B-IR* | | | <0.0001 | 12 |
| **Fig 1J 3$^{rd}$ order** | | **Two-way ANOVA with Dunnett's test for multiple comparison** | | |
| Control | | | | 10 |
| *mts-IR* | | | <0.0001 | 12 |
| *PP2A-29B-IR* | | | <0.0001 | 12 |
| **Fig 1J 4$^{th}$ order** | | **Two-way ANOVA with Dunnett's test for multiple comparison** | | |
| Control | | | | 10 |
| *mts-IR* | | | 0.0008 | 12 |
| *PP2A-29B-IR* | | | 0.0522 | 12 |
| **Fig 1K 5$^{th}$ order** | | **Two-way ANOVA with Dunnett's test for multiple comparison** | | |
| Control | | | | 10 |
| *mts-IR* | | | <0.0001 | 12 |
| *PP2A-29B-IR* | | | <0.0001 | 12 |
| **Fig 1K 6$^{th}$ order** | | **Two-way ANOVA with Dunnett's test for multiple comparison** | | |
| Control | | | | 10 |
| *mts-IR* | | | 0.7324 | 12 |
| *PP2A-29B-IR* | | | 0.7531 | 12 |
| **Fig 1K 7$^{th}$ order** | | **Two-way ANOVA with Dunnett's test for multiple comparison** | | |
| Control | | | | 10 |
| *mts-IR* | | | 0.9976 | 12 |
| *PP2A-29B-IR* | | | 0.9976 | 12 |
| **Fig 1L 1$^{st}$ order** | | **Two-way ANOVA with Dunnett's test for multiple comparison** | | |
| Control-MARCM | | | | 15 |
| $mts^{k12502}$ | | | <0.0001 | 12 |
| **Fig 1L 2$^{nd}$order** | | **Two-way ANOVA with Dunnett's test for multiple comparison** | | |

| | | | | |
|---|---|---|---|---|
| Control-MARCM | | | | 15 |
| *mts*$^{k12502}$ | | | <0.0001 | 12 |
| **Fig 1L 3rd order** | | **Two-way ANOVA with Dunnett's test for multiple comparison** | | |
| Control-MARCM | | | | 15 |
| *mts*$^{k12502}$ | | | <0.0001 | 12 |
| **Fig 1L 4th order** | | **Two-way ANOVA with Dunnett's test for multiple comparison** | | |
| Control-MARCM | | | | 15 |
| *mts*$^{k12502}$ | | | 0.0054 | 12 |
| **Fig 1M 5th order** | | **Two-way ANOVA with Dunnett's test for multiple comparison** | | |
| Control-MARCM | | | | 15 |
| *mts*$^{k12502}$ | | | 0.0019 | 12 |
| **Fig 1M 6th order** | | **Two-way ANOVA with Dunnett's test for multiple comparison** | | |
| Control-MARCM | | | | 15 |
| *mts*$^{k12502}$ | | | 0.7686 | 12 |
| **Fig 1M 7th order** | | **Two-way ANOVA with Dunnett's test for multiple comparison** | | |
| Control-MARCM | | | | 15 |
| *mts*$^{k12502}$ | | | 0.9942 | 12 |
| **Fig 1T** | | **Kruskal-Wallis with Dunn's multiple comparison test** | | |
| Control | No | | | 20 |
| *mts-IR* | Yes | | 0.0007 | 21 |
| *PP2A-29B-IR* | Yes | | >0.9999 | 18 |
| *wdb-IR* | Yes | | >0.9999 | 18 |
| Control-MARCM | No | | | 34 |
| *mts*$^{k12502}$ | No | | <0.0001 | 12 |
| *wdb*$^{14}$ | No | | 0.0143 | 12 |
| **Fig 1U** | | **Kruskal-Wallis with Dunn's multiple comparison test** | | |
| Control | Yes | | | 20 |
| *mts-IR* | Yes | | <0.0001 | 21 |
| *PP2A-29B-IR* | Yes | | <0.0001 | 18 |
| *wdb-IR* | No | | 0.0176 | 18 |
| Control-MARCM | No | | | 34 |

| | | | | |
|---|---|---|---|---|
| *mts*$^{k12502}$ | Yes | | 0.18390 | 12 |
| *wdb*$^{14}$ | Yes | | >0.9999 | 12 |
| **Fig 1V** | | **Kruskal-Wallis with Dunn's multiple comparison test** | | |
| Control | No | | | 20 |
| *mts-IR* | Yes | | <0.0001 | 21 |
| *PP2A-29B-IR* | Yes | | <0.0001 | 18 |
| *wdb-IR* | Yes | | 0.1337 | 18 |
| Control-MARCM | No | | | 34 |
| *mts*$^{k12502}$ | Yes | | 0.0011 | 12 |
| *wdb*$^{14}$ | Yes | | >0.9999 | 12 |
| **Fig 1W 1$^{st}$ order** | | **Two-way ANOVA with Dunnett's test for multiple comparison** | | |
| Control | | | | 21 |
| *mts-IR* | | | <0.0001 | 20 |
| *PP2A-29B-IR* | | | <0.0001 | 19 |
| *wdb-IR* | | | 0.0567 | 21 |
| **Fig 1W 2$^{nd}$ order** | | **Two-way ANOVA with Dunnett's test for multiple comparison** | | |
| Control | | | | 21 |
| *mts-IR* | | | 0.0032 | 20 |
| *PP2A-29B-IR* | | | <0.0001 | 19 |
| *wdb-IR* | | | 0.3571 | 21 |
| **Fig 1W 3$^{rd}$ order** | | **Two-way ANOVA with Dunnett's test for multiple comparison** | | |
| Control | | | | 21 |
| *mts-IR* | | | 0.4371 | 20 |
| *PP2A-29B-IR* | | | 0.0036 | 19 |
| *wdb-IR* | | | 0.9226 | 21 |
| **Fig 1W 4$^{th}$ order** | | **Two-way ANOVA with Dunnett's test for multiple comparison** | | |
| Control | | | | 21 |
| *mts-IR* | | | 0.9817 | 20 |
| *PP2A-29B-IR* | | | 0.8562 | 19 |
| *wdb-IR* | | | 0.9841 | 21 |
| **Fig 1X 1$^{st}$ order** | | **Two-way ANOVA with Dunnett's test for multiple comparison** | | |
| Control-MARCM | | | | 33 |
| *mts*$^{k12502}$ | | | <0.0001 | 12 |

| | | | | |
|---|---|---|---|---|
| *wdb*$^{14}$ | | | 0.1298 | 12 |
| **Fig 1X 2$^{nd}$ order** | | **Two-way ANOVA with Dunnett's test for multiple comparison** | | |
| Control-MARCM | | | | 33 |
| *mts*$^{k12502}$ | | | 0.0038 | 12 |
| *wdb*$^{14}$ | | | 0.7993 | 12 |
| **Fig 1X 3$^{rd}$ order** | | **Two-way ANOVA with Dunnett's test for multiple comparison** | | |
| Control-MARCM | | | | 33 |
| *mts*$^{k12502}$ | | | 0.0721 | 12 |
| *wdb*$^{14}$ | | | 0.6939 | 12 |
| **Fig 1X 4$^{th}$ order** | | **Two-way ANOVA with Dunnett's test for multiple comparison** | | |
| Control-MARCM | | | | 33 |
| *mts*$^{k12502}$ | | | 0.8137 | 12 |
| *wdb*$^{14}$ | | | 0.7288 | 12 |
| **Fig 2E** | | **One-way ANOVA with Dunnett's multiple comparison test** | | |
| Control | Yes | | | 10 |
| *mts-OE* | Yes | | <0.0001 | 10 |
| *PP2A-29B-OE* | Yes | | 0.0084 | 14 |
| *wdb-OE* | Yes | | <0.0001 | 11 |
| **Fig 2F** | | **One-way ANOVA with Dunnett's multiple comparison test** | | |
| Control | Yes | | | 10 |
| *mts-OE* | Yes | | <0.0001 | 10 |
| *PP2A-29B-OE* | Yes | | <0.0001 | 14 |
| *wdb-OE* | Yes | | <0.0001 | 11 |
| **Fig 2G** | | **One-way ANOVA with Dunnett's multiple comparison test** | | |
| Control | Yes | | | 10 |
| *mts-OE* | Yes | | <0.0001 | 10 |
| *PP2A-29B-OE* | Yes | | <0.0001 | 14 |
| *wdb-OE* | Yes | | <0.0001 | 11 |
| **Fig 2L** | | **Kruskal-Wallis with Dunn's multiple comparison test** | | |
| Control | Yes | | | 10 |
| *cka-IR* | Yes | | <0.0001 | 11 |

| | | | | |
|---|---|---|---|---|
| *cka-OE* | Yes | | 0.0551 | 13 |
| *BFP-cka*$^{\Delta PP2A}$ | No | | >0.9999 | 13 |
| **Fig 2M** | | **Kruskal-Wallis with Dunn's multiple comparison test** | | |
| Control | Yes | | | 10 |
| *cka-IR* | No | | 0.0006 | 11 |
| *cka-OE* | Yes | | >0.9999 | 13 |
| *BFP-cka*$^{\Delta PP2A}$ | No | | 0.9246 | 13 |
| **Fig 2N 1$^{st}$ order** | | **Two-way ANOVA with Dunnett's test for multiple comparison** | | |
| Control | | | | 10 |
| *cka-IR* | | | <0.0001 | 11 |
| *cka-OE* | | | 0.4681 | 13 |
| *BFP-cka*$^{\Delta PP2A}$ | | | 0.0069 | 13 |
| **Fig 2N 2$^{nd}$ order** | | **Two-way ANOVA with Dunnett's test for multiple comparison** | | |
| Control | | | | 10 |
| *cka-IR* | | | 0.0001 | 11 |
| *cka-OE* | | | 0.6904 | 13 |
| *BFP-cka*$^{\Delta PP2A}$ | | | 0.4179 | 13 |
| **Fig 2N 3$^{rd}$ order** | | **Two-way ANOVA with Dunnett's test for multiple comparison** | | |
| Control | | | | 10 |
| *cka-IR* | | | 0.0808 | 11 |
| *cka-OE* | | | 0.9979 | 13 |
| *BFP-cka*$^{\Delta PP2A}$ | | | >0.9999 | 13 |
| **Fig 2N 4$^{th}$ -7$^{th}$ order** | | **Two-way ANOVA with Dunnett's test for multiple comparison** | | |
| Control | | | | 10 |
| *cka-IR* | | | 0.4080 | 11 |
| *cka-OE* | | | 0.9780 | 13 |
| *BFP-cka*$^{\Delta PP2A}$ | | | 0.3034 | 13 |
| **Fig 3C** | | **Two-way ANOVA with Sidak's test for multiple comparison** | | |
| Control-24h AEL | | | | 17 |
| *mts-IR*-24h AEL | | | >0.9999 | 11 |
| Control-48h AEL | | | | 20 |
| *mts-IR*-48h AEL | | | 0.9998 | 10 |
| Control-72h AEL | | | | 11 |

| | | | | |
|---|---|---|---|---|
| *mts-IR*-72h AEL | | | <0.0001 | 10 |
| Control-96h AEL | | | | 10 |
| *mts-IR*-96h AEL | | | <0.0001 | 9 |
| **Fig 3D** | | **Two-way ANOVA with Sidak's test for multiple comparison** | | |
| Control-24h AEL | | | | 17 |
| *mts-IR*-24h AEL | | | 0.9773 | 11 |
| Control-48h AEL | | | | 20 |
| *mts-IR*-48h AEL | | | 0.2244 | 10 |
| Control-72h AEL | | | | 11 |
| *mts-IR*-72h AEL | | | <0.0001 | 10 |
| Control-96h AEL | | | | 10 |
| *mts-IR*-96h AEL | | | <0.0001 | 9 |
| **Fig 3E** | | **Two-way ANOVA with Sidak's test for multiple comparison** | | |
| Control-24h AEL | | | | 17 |
| *mts-IR*-24h AEL | | | 0.0762 | 11 |
| Control-48h AEL | | | | 20 |
| *mts-IR*-48h AEL | | | 0.0004 | 10 |
| Control-72h AEL | | | | 11 |
| *mts-IR*-72h AEL | | | <0.0001 | 10 |
| Control-96h AEL | | | | 10 |
| *mts-IR*-96h AEL | | | <0.0001 | 9 |
| **Fig 4D** | | **Unpaired t-test** | | |
| Control *(mts)* | Yes | | | 4 |
| *ct-IR (mts)* | Yes | | <0.0001 | 4 |
| **Fig 4E** | | **One-way ANOVA with Sidak's multiple comparison test** | | |
| Control | Yes | | | 9 |
| Control *vs ct-IR* | Yes | | <0.0001 | 9, 8 |
| *ct-IR vs mts-OE;ct-IR* | Yes | | 0.0167 | 8, 17 |
| **Fig 4F** | | **One-way ANOVA with Sidak's multiple comparison test** | | |
| Control | Yes | | | 9 |
| Control *vs ct-IR* | Yes | | <0.0001 | 9, 8 |
| *ct-IR vs mts-OE;ct-IR* | Yes | | 0.0187 | 8, 17 |
| **Fig 4G 1$^{st}$ order** | | **Two-way ANOVA with Tukey's multiple comparison test** | | |
| Control | | | | 9 |
| Control *vs ct-IR* | | | <0.0001 | 9, 8 |

| | | | | |
|---|---|---|---|---|
| *ct-IR vs mts-OE;ct-IR* | | | <0.0001 | 8, 17 |
| **Fig 4G**<br>**2nd order** | | **Two-way ANOVA with Tukey's multiple comparison test** | | |
| Control | | | | 9 |
| Control *vs ct-IR* | | | <0.0001 | 9, 8 |
| *ct-IR vs mts-OE;ct-IR* | | | 0.1024 | 8, 17 |
| **Fig 4G**<br>**3rd order** | | **Two-way ANOVA with Tukey's multiple comparison test** | | |
| Control | | | | 9 |
| Control *vs ct-IR* | | | <0.0001 | 9, 8 |
| *ct-IR vs mts-OE;ct-IR* | | | 0.8353 | 8, 17 |
| **Fig 4G**<br>**4th order** | | **Two-way ANOVA with Tukey's multiple comparison test** | | |
| Control | | | | 9 |
| Control *vs ct-IR* | | | 0.1291 | 9, 8 |
| *ct-IR vs mts-OE;ct-IR* | | | 0.8353 | 8, 17 |
| **Fig 4H**<br>**5th order** | | **Two-way ANOVA with Tukey's multiple comparison test** | | |
| Control | | | | 9 |
| Control *vs ct-IR* | | | <0.0001 | 9, 8 |
| *ct-IR vs mts-OE;ct-IR* | | | 0.0648 | 8, 17 |
| **Fig 4H**<br>**6th order** | | **Two-way ANOVA with Tukey's multiple comparison test** | | |
| Control | | | | 9 |
| Contro*l vs ct-IR* | | | 0.3278 | 9, 8 |
| *ct-IR vs mts-OE;ct-IR* | | | 0.9915 | 8, 17 |
| **Fig 4H**<br>**7th order** | | **Two-way ANOVA with Tukey's multiple comparison test** | | |
| Control | | | | 9 |
| Control *vs ct-IR* | | | 0.9544 | 9, 8 |
| *ct-IR vs mts-OE;ct-IR* | | | 0.9421 | 8, 17 |
| **Fig 5E**<br>**40µm from soma** | | **Unpaired t-test with false discovery rate correction** | | |
| Control | | | | 11 |
| *mts-IR* | | | <0.0001 | 11 |
| **Fig 5E**<br>**80µm from soma** | | **Unpaired t-test with false discovery rate correction** | | |

| | | | | |
|---|---|---|---|---|
| Control | | | | 11 |
| *mts-IR* | | | <0.0001 | 11 |
| **Fig 5E 120μm from soma** | | **Unpaired t-test with false discovery rate correction** | | |
| Control | | | | 11 |
| *mts-IR* | | | <0.0001 | 11 |
| **Fig 5E 160μm from soma** | | **Unpaired t-test with false discovery rate correction** | | |
| Control | | | | 11 |
| *mts-IR* | | | <0.0001 | 11 |
| **Fig 5E 200μm from soma** | | **Unpaired t-test with false discovery rate correction** | | |
| Control | | | | 11 |
| *mts-IR* | | | <0.0001 | 11 |
| **Fig 5E 240μm from soma** | | **Unpaired t-test with false discovery rate correction** | | |
| Control | | | | 11 |
| *mts-IR* | | | <0.0001 | 11 |
| **Fig 5E 280μm from soma** | | **Unpaired t-test with false discovery rate correction** | | |
| Control | | | | 11 |
| *mts-IR* | | | <0.0001 | 11 |
| **Fig 5E 320μm from soma** | | **Unpaired t-test with false discovery rate correction** | | |
| Control | | | | 11 |
| *mts-IR* | | | <0.0001 | 11 |
| **Fig 5E 360μm from soma** | | **Unpaired t-test with false discovery rate correction** | | |
| Control | | | | 11 |
| *mts-IR* | | | <0.0001 | 11 |
| **Fig 5E 400μm from soma** | | **Unpaired t-test with false discovery rate correction** | | |
| Control | | | | 11 |
| *mts-IR* | | | <0.0001 | 11 |
| **Fig 5E 440μm from soma** | | **Unpaired t-test with false discovery rate correction** | | |

| | | | | |
|---|---|---|---|---|
| Control | | | | 11 |
| *mts-IR* | | | <0.0001 | 11 |
| **Fig 5E**<br>**480µm from soma** | | **Unpaired t-test with false discovery rate correction** | | |
| Control | | | | 11 |
| *mts-IR* | | | <0.0001 | 11 |
| **Fig 5E**<br>**520µm from soma** | | **Unpaired t-test with false discovery rate correction** | | |
| Control | | | | 11 |
| *mts-IR* | | | 0.0106 | 11 |
| **Fig 5E**<br>**560µm from soma** | | **Unpaired t-test with false discovery rate correction** | | |
| Control | | | | 11 |
| *mts-IR* | | | 0.1065 | 11 |
| **Fig 5E**<br>**600µm from soma** | | **Unpaired t-test with false discovery rate correction** | | |
| Control | | | | 11 |
| *mts-IR* | | | 0.1140 | 11 |
| **Fig 5E**<br>**640µm from soma** | | **Unpaired t-test with false discovery rate correction** | | |
| Control | | | | 11 |
| *mts-IR* | | | 0.1502 | 11 |
| **Fig 5F**<br>**1$^{st}$ order** | | **Unpaired t-test with false discovery rate correction** | | |
| Control | | | | 11 |
| *mts-IR* | | | 0.0058 | 11 |
| **Fig 5F**<br>**2$^{nd}$ order** | | **Unpaired t-test with false discovery rate correction** | | |
| Control | | | | 11 |
| *mts-IR* | | | <0.0001 | 11 |
| **Fig 5F**<br>**3$^{rd}$ order** | | **Unpaired t-test with false discovery rate correction** | | |
| Control | | | | 11 |
| *mts-IR* | | | <0.0001 | 11 |
| **Fig 5F**<br>**4$^{th}$ order** | | **Unpaired t-test with false discovery rate correction** | | |

| | | | | |
|---|---|---|---|---|
| Control | | | | 11 |
| *mts-IR* | | | <0.0001 | 11 |
| **Fig 5F<br>5th order** | | **Unpaired t-test with false discovery rate correction** | | |
| Control | | | | 11 |
| *mts-IR* | | | <0.0001 | 11 |
| **Fig 5G<br>40µm from soma** | | **Unpaired t-test with false discovery rate correction** | | |
| Control | | | | 11 |
| *mts-IR* | | | 0.1732 | 11 |
| **Fig 5G<br>80µm from soma** | | **Unpaired t-test with false discovery rate correction** | | |
| Control | | | | 11 |
| *mts-IR* | | | 0.1114 | 11 |
| **Fig 5G<br>120µm from soma** | | **Unpaired t-test with false discovery rate correction** | | |
| Control | | | | 11 |
| *mts-IR* | | | 0.0582 | 11 |
| **Fig 5G<br>160µm from soma** | | **Unpaired t-test with false discovery rate correction** | | |
| Control | | | | 11 |
| *mts-IR* | | | 0.0582 | 11 |
| **Fig 5G<br>200µm from soma** | | **Unpaired t-test with false discovery rate correction** | | |
| Control | | | | 11 |
| *mts-IR* | | | 0.0497 | 11 |
| **Fig 5G<br>240µm from soma** | | **Unpaired t-test with false discovery rate correction** | | |
| Control | | | | 11 |
| *mts-IR* | | | 0.1732 | 11 |
| **Fig 5G<br>280µm from soma** | | **Unpaired t-test with false discovery rate correction** | | |
| Control | | | | 11 |
| *mts-IR* | | | 0.0727 | 11 |
| **Fig 5G<br>320µm from soma** | | **Unpaired t-test with false discovery rate correction** | | |

| | | | | |
|---|---|---|---|---|
| Control | | | | 11 |
| *mts-IR* | | | <0.0001 | 11 |
| **Fig 5G 360µm from soma** | | **Unpaired t-test with false discovery rate correction** | | |
| Control | | | | 11 |
| *mts-IR* | | | <0.0001 | 11 |
| **Fig 5G 400µm from soma** | | **Unpaired t-test with false discovery rate correction** | | |
| Control | | | | 11 |
| *mts-IR* | | | <0.0001 | 11 |
| **Fig 5G 440µm from soma** | | **Unpaired t-test with false discovery rate correction** | | |
| Control | | | | 11 |
| *mts-IR* | | | <0.0001 | 11 |
| **Fig 5G 480µm from soma** | | **Unpaired t-test with false discovery rate correction** | | |
| Control | | | | 11 |
| *mts-IR* | | | 0.0025 | 11 |
| **Fig 5G 520µm from soma** | | **Unpaired t-test with false discovery rate correction** | | |
| Control | | | | 11 |
| *mts-IR* | | | 0.0510 | 11 |
| **Fig 5G 560µm from soma** | | **Unpaired t-test with false discovery rate correction** | | |
| Control | | | | 11 |
| *mts-IR* | | | 0.0582 | 11 |
| **Fig 5G 600µm from soma** | | **Unpaired t-test with false discovery rate correction** | | |
| Control | | | | 11 |
| *mts-IR* | | | 0.2530 | 11 |
| **Fig 5G 640µm from soma** | | **Unpaired t-test with false discovery rate correction** | | |
| Control | | | | 11 |
| *mts-IR* | | | 0.4808 | 11 |
| **Fig 5H** | | **Unpaired t-test** | | |
| Control | Yes | | | 11 |
| *mts-IR* | Yes | | <0.0001 | 11 |

| **Fig 5I** | | **Unpaired t-test** | | |
|---|---|---|---|---|
| Control | Yes | | | 11 |
| *mts-IR* | Yes | | <0.0001 | 11 |
| **Fig 5J**<br>**1st order** | | **Mann-Whitney test with FDR correction** | | |
| Control | | | | 11 |
| *mts-IR* | | | <0.0001 | 11 |
| **Fig 5J**<br>**2nd order** | | **Mann-Whitney test with FDR correction** | | |
| Control | | | | 11 |
| *mts-IR* | | | <0.0001 | 11 |
| **Fig 5J**<br>**3rd order** | | **Mann-Whitney test with FDR correction** | | |
| Control | | | | 11 |
| *mts-IR* | | | <0.0001 | 11 |
| **Fig 5J**<br>**4th order** | | **Mann-Whitney test with FDR correction** | | |
| Control | | | | 11 |
| *mts-IR* | | | <0.0001 | 11 |
| **Fig 5J**<br>**5th order** | | **Mann-Whitney test with FDR correction** | | |
| Control | | | | 11 |
| *mts-IR* | | | 0.0169 | 11 |
| **Fig 5J**<br>**6th order** | | **Mann-Whitney test with FDR correction** | | |
| Control | | | | 11 |
| *mts-IR* | | | 0.0004 | 11 |
| **Fig 5O** | | **Unpaired t-test** | | |
| Control | | | | 8 |
| *mts-IR* | | | <0.0001 | 9 |
| **Fig 5P**<br>**1st order** | | **Unpaired t-test with false discovery rate correction** | | |
| Control | | | | 8 |
| *mts-IR* | | | <0.0001 | 9 |
| **Fig 5P**<br>**2nd order** | | **Unpaired t-test with false discovery rate correction** | | |
| Control | | | | 8 |
| *mts-IR* | | | <0.0001 | 9 |
| **Fig 5P**<br>**3rd order** | | **Unpaired t-test with false discovery rate correction** | | |
| Control | | | | 8 |

| | | | | |
|---|---|---|---|---|
| *mts-IR* | | | 0.0001 | 9 |
| **Fig 5P<br>4$^{th}$ order** | | **Unpaired t-test with false discovery rate correction** | | |
| Control | | | | 8 |
| *mts-IR* | | | 0.0384 | 9 |
| **Fig 5Q** | | **Unpaired t-test** | | |
| Control | | | | 8 |
| *mts-IR* | | | 0.0002 | 9 |
| **Fig 5R<br>1$^{st}$ order** | | **Mann-Whitney test with FDR correction** | | |
| Control | | | | 8 |
| *mts-IR* | | | 0.0002 | 9 |
| **Fig 5R<br>2$^{nd}$ order** | | **Mann-Whitney test with FDR correction** | | |
| Control | | | | 8 |
| *mts-IR* | | | 0.0003 | 9 |
| **Fig 5R<br>3$^{rd}$ order** | | **Mann-Whitney test with FDR correction** | | |
| Control | | | | 8 |
| *mts-IR* | | | 0.03646 | 9 |
| **Fig 5R<br>4$^{th}$ order** | | **Mann-Whitney test with FDR correction** | | |
| Control | | | | 8 |
| *mts-IR* | | | 0.3813 | 9 |
| **Fig 5S** | | **Mann-Whitney test** | | |
| Control | No | | | 8 |
| *mts-IR* | No | | 0.0498 | 9 |
| **Fig 6C<br>0 min** | | **Two-way ANOVA with Sidak's multiple comparison test** | | |
| Control | | | | 11 |
| *mts-IR* | | | >0.9999 | 8 |
| **Fig 6C<br>30 min** | | **Two-way ANOVA with Sidak's multiple comparison test** | | |
| Control | | | | 11 |
| *mts-IR* | | | 0.0001 | 8 |
| **Fig 6C<br>60 min** | | **Two-way ANOVA with Sidak's multiple comparison test** | | |
| Control | | | | 11 |
| *mts-IR* | | | 0.0028 | 8 |

| | | | | |
|---|---|---|---|---|
| **Fig 6F<br>0 min** | | **Two-way ANOVA with Sidak's multiple comparison test** | | |
| Control | | | | 13 |
| *mts-IR* | | | >0.9999 | 10 |
| **Fig 6F<br>30 min** | | **Two-way ANOVA with Sidak's multiple comparison test** | | |
| Control | | | | 13 |
| *mts-IR* | | | 0.0006 | 10 |
| **Fig 6F<br>60 min** | | **Two-way ANOVA with Sidak's multiple comparison test** | | |
| Control | | | | 13 |
| *mts-IR* | | | <0.0001 | 10 |
| **Fig 7C<br>Retrograde** | | **Two-way ANOVA with Sidak 's multiple comparison test** | | |
| Control | | | | 72<br>(comets) |
| *mts-IR* | | | <0.0001 | 70<br>(comets) |
| **Fig 7C<br>Anterograde** | | **Two-way ANOVA with Sidak 's multiple comparison test** | | |
| Control | | | | 72 (comets) |
| *mts-IR* | | | <0.0001 | 70 (comets) |
| **Fig 7D** | | **Mann Whitney test** | | |
| Control | No | | | 13 |
| *mts-IR* | Yes | | <0.0001 | 9 |
| **Fig 7E<br>Retrograde** | | **Two-way ANOVA with Dunnett's multiple comparison test** | | |
| Control | | | | 54 (comets) |
| *mts-IR* | | | 0.0074 | 43 (comets) |
| **Fig 7E<br>Anterograde** | | **Two-way ANOVA with Dunnett's multiple comparison test** | | |
| Control | | | | 54 (comets) |
| *mts-IR* | | | 0.0074 | 43 (comets) |
| **Fig 7F** | | **Unpaired t-test** | | |
| Control | Yes | | | 14 |
| *mts-IR* | Yes | | 0.9595 | 9 |
| **Fig 7G** | | **Mann-Whitney test** | | |
| Control | No | | | 60 (comets) |

| | | | | |
|---|---|---|---|---|
| *mts-IR* | Yes | | 0.8499 | 29 (comets) |
| **Fig 7H** | | **Mann-Whitney test** | | |
| Control | No | | | 67 (comets) |
| *mts-IR* | Yes | | 0.0001 | 36 (comets) |
| **Fig 7I** | | **Mann-Whitney test** | | |
| Control | No | | | 27 (comets) |
| *mts-IR* | No | | 0.4224 | 19 (comets) |
| **Fig 7J** | | **Mann-Whitney test** | | |
| Control | No | | | 34 (comets) |
| *mts-IR* | No | | 0.0312 | 26 (comets) |
| **Fig 7K CI Anterograde** | | **Two-way ANOVA with Dunnett's multiple comparison test** | | |
| Control | Yes | | | 42 (comets) |
| *mts-IR* | Yes | | <0.0001 | 53 (comets) |
| **Fig 7K CI Retrograde** | | **Two-way ANOVA with Dunnett's multiple comparison test** | | |
| Control | Yes | | | 42 (comets) |
| *mts-IR* | Yes | | <0.0001 | 53 (comets) |
| **Fig 7L** | | **Mann-Whitney test** | | |
| Control | No | | | 52 (comets) |
| *mts-IR* | No | | <0.0001 | 67(comets) |
| **Fig 7M** | | **Mann-Whitney test** | | |
| Control | No | | | 15 |
| *mts-IR* | Yes | | <0.0001 | 9 |
| **Fig 7N** | | **Mann-Whitney test** | | |
| Control | No | | | 52 (comets) |
| *mts-IR* | No | | 0.1145 | 67(comets) |
| **Fig 8C** | | **Unpaired t-test** | | |
| Control | Yes | | | 10 |
| *mts-IR* | Yes | | <0.0001 | 10 |
| **Fig 8F** | | **Mann-Whitney test** | | |
| Control | Yes | | | 8 |
| *mts-IR* | No | | 0.0062 | 10 |
| **Fig 8K** | | **Unpaired t-test** | | |
| Control | Yes | | | 10 |
| *mts-IR* | Yes | | <0.0001 | 11 |
| **Fig 8L** | | **Unpaired t-test** | | |
| Control | Yes | | | 10 |
| *mts-IR* | Yes | | <0.0001 | 11 |
| **Fig 8M** | | **Unpaired t-test** | | |
| Control | Yes | | | 7 |
| *mts-IR* | Yes | | 0.0004 | 14 |
| **Fig 8N** | | **Unpaired t-test** | | |

| | | | | |
|---|---|---|---|---|
| Control | Yes | | | 7 |
| *mts-IR* | Yes | | 0.0002 | 14 |
| **Fig 9E** | | **Unpaired t-test** | | |
| Control | Yes | | | 19 |
| *mts-IR* | Yes | | <0.0001 | 13 |
| **Fig 9F** | | **Unpaired t-test** | | |
| Control | Yes | | | 19 |
| *mts-IR* | Yes | | <0.0001 | 13 |
| **Fig 9G** | | **Unpaired t-test** | | |
| Control | Yes | | | 10 |
| *mts-IR* | Yes | | 0.0357 | 12 |
| **Fig 9H** | | **Mann-Whitney test** | | |
| Control | No | | | 10 |
| *mts-IR* | Yes | | 0.6676 | 11 |
| **Fig 9P** | | **Mann-Whitney test** | | |
| Control | No | | | 18 |
| *mts-IR* | No | | <0.0001 | 12 |
| **Fig 9S** | | **Unpaired t-test** | | |
| Control | Yes | | | 10 |
| *mts-IR* | Yes | | 0.0025 | 12 |
| **Fig 10E** | | **Unpaired t-test** | | |
| Control | Yes | | | 23 |
| *mts-IR* | Yes | | 0.0005 | 29 |
| **Fig 10F** | | **Unpaired t-test** | | |
| Control | Yes | | | 10 |
| *foxo-OE* | Yes | | <0.0001 | 12 |
| **Fig 10G** | | **Unpaired t-test** | | |
| Control | Yes | | | 10 |
| *foxo-OE* | Yes | | 0.0005 | 12 |
| **Fig 10H** | | **Unpaired t-test** | | |
| Control | Yes | | | 10 |
| *foxo-OE* | Yes | | <0.0001 | 12 |
| **Fig 10M** | | **One-way ANOVA with Sidak's multiple comparison test** | | |
| Control | Yes | | | 10 |
| Control *vs mts-OE/foxo-OE* | Yes | | 0.0113 | 10, 14 |
| *mts-IR vs mts-OE/foxo-OE* | Yes | | <0.0001 | 12, 14 |
| **Fig 10N** | | **One-way ANOVA with Sidak's multiple comparison test** | | |
| Control | Yes | | | 10 |

| | | | | |
|---|---|---|---|---|
| Control *vs mts-OE/foxo-OE* | Yes | | 0.1984 | 10, 14 |
| *mts-IR vs mts-OE/foxo-OE* | Yes | | <0.0001 | 12, 14 |
| **Fig 11D** | | **One-way ANOVA with Dunnett's multiple comparison test** | | |
| Control | Yes | | | 13 |
| *β-tubulin 85D-IR* | Yes | | <0.0001 | 13 |
| *β-tubulin 85D-OE* | Yes | | 0.9620 | 13 |
| **Fig 11E** | | **One-way ANOVA with Dunnett's multiple comparison test** | | |
| Control | Yes | | | 13 |
| *β-tubulin 85D-IR* | Yes | | <0.0001 | 13 |
| *β-tubulin 85D-OE* | Yes | | 0.3584 | 13 |
| **Fig 11I** | | **Kruskal-Wallis with Dunn's multiple comparison test** | | |
| Control | Yes | | | 10 |
| Control *vs mts-IR* | No | | <0.0001 | 10, 11 |
| *mts-IR vs mts-IR; β-tubulin 85D-OE* | Yes | | 0.0451 | 11, 12 |
| **Fig 11J** | | **Kruskal-Wallis with Dunn's multiple comparison test** | | |
| Control | Yes | | | 10 |
| Control *vs mts-IR* | No | | <0.0001 | 10, 11 |
| *mts-IR vs mts-IR; β-tubulin 85D-OE* | Yes | | 0.0290 | 11, 12 |
| **Fig 11K** | | **One-way ANOVA with Sidak's multiple comparison test** | | |
| Control | Yes | | | 10 |
| Control *vs mts-IR* | Yes | | <0.0001 | 10, 11 |
| *mts-IR vs mts-IR; β-tubulin 85D-OE* | Yes | | 0.0001 | 11, 12 |
| **Fig 11L 1$^{st}$ order** | | **Two-way ANOVA with Tukey's multiple comparison test** | | |
| Control | | | | 10 |
| Control *vs mts-IR* | | | <0.0001 | 10, 11 |
| *mts-IR vs mts-IR; β-tubulin 85D-OE* | | | <0.0001 | 11, 12 |

| | | | | |
|---|---|---|---|---|
| **Fig 11L 2nd order** | | **Two-way ANOVA with Tukey's multiple comparison test** | | |
| Control | | | | 10 |
| Control *vs mts-IR* | | | <0.0001 | 10, 11 |
| *mts-IR vs mts-IR; β-tubulin 85D-OE* | | | <0.0001 | 11, 12 |
| **Fig 11L 3rd order** | | **Two-way ANOVA with Tukey's multiple comparison test** | | |
| Control | | | | 10 |
| Control *vs mts-IR* | | | <0.0001 | 10, 11 |
| *mts-IR vs mts-IR; β-tubulin 85D-OE* | | | 0.1186 | 11, 12 |
| **Fig 11L 4th order** | | **Two-way ANOVA with Tukey's multiple comparison test** | | |
| Control | | | | 10 |
| Control *vs mts-IR* | | | 0.0006 | 10, 11 |
| *mts-IR vs mts-IR; β-tubulin 85D-OE* | | | 0.3313 | 11, 12 |
| **Fig 11L 5th order** | | **Two-way ANOVA with Tukey's multiple comparison test** | | |
| Control | | | | 10 |
| Control *vs mts-IR* | | | 0.6880 | 10, 11 |
| *mts-IR vs mts-IR; β-tubulin 85D-OE* | | | 0.9638 | 11, 12 |
| **Fig 11M 6th order** | | **Two-way ANOVA with Tukey's multiple comparison test** | | |
| Control | | | | 10 |
| Control *vs mts-IR* | | | 0.1067 | 10, 11 |
| *mts-IR vs mts-IR; β-tubulin 85D-OE* | | | 0.7650 | 11, 12 |
| **Fig 11M 7th order** | | **Two-way ANOVA with Tukey's multiple comparison test** | | |
| Control | | | | 10 |
| Control *vs mts-IR* | | | 0.9247 | 10, 11 |
| *mts-IR vs mts-IR; β-tubulin 85D-OE* | | | 0.9961 | 11, 12 |
| **Fig 11N** | | **One-way ANOVA with Sidak's multiple comparison test** | | |

| | | | | |
|---|---|---|---|---|
| Control | Yes | | | 10 |
| Control *vs mts-IR* | Yes | | <0.0001 | 10, 11 |
| *mts-IR vs mts-IR; β-tubulin 85D-OE* | Yes | | <0.0001 | 11, 12 |
| **Fig 11R** | | **Unpaired t-test** | | |
| Control | Yes | | | 24 |
| *mts-IR* | Yes | | <0.0001 | 32 |
| **Fig 12C** | | **Unpaired t-test** | | |
| Control | Yes | | | 10 |
| *β-tubulin 85D-IR* | Yes | | 0.0628 | 19 |
| **Fig 12D** | | **Unpaired t-test** | | |
| Control | Yes | | | 10 |
| *β-tubulin 85D-IR* | Yes | | 0.0237 | 19 |
| **Fig 12E** | | **Unpaired t-test** | | |
| Control | Yes | | | 10 |
| *β-tubulin 85D-IR* | Yes | | <0.0001 | 19 |
| **Fig 12F 1st order** | | **Unpaired t-test with FDR correction** | | |
| Control | Yes | | | 10 |
| *β-tubulin 85D-IR* | Yes | | 0.0048 | 19 |
| **Fig 12F 2nd order** | | **Unpaired t-test with FDR correction** | | |
| Control | Yes | | | 10 |
| *β-tubulin 85D-IR* | Yes | | 0.0023 | 19 |
| **Fig 12F 3rd order** | | **Unpaired t-test with FDR correction** | | |
| Control | Yes | | | 10 |
| *β-tubulin 85D-IR* | Yes | | 0.1082 | 19 |
| **Fig 12F 4th order** | | **Unpaired t-test with FDR correction** | | |
| Control | Yes | | | 10 |
| *β-tubulin 85D-IR* | Yes | | 0.0285 | 19 |
| **Fig 12K** | | **One-way ANOVA with Sidak's multiple comparison test** | | |
| Control | Yes | | | 10 |
| Control *vs mts-IR;β-tubulin 85D-OE* | Yes | | 0.0338 | 10, 19 |
| *mts-IR vs mts-IR;β-tubulin 85D-OE* | Yes | | <0.0001 | 14, 19 |
| **Fig 12L** | | **One-way ANOVA with Sidak's multiple comparison test** | | |
| Control | Yes | | | 10 |

| | | | | |
|---|---|---|---|---|
| Control *vs* *mts-IR;β-tubulin 85D-OE* | Yes | | 0.0003 | 10, 19 |
| *mts-IR vs mts-IR;β-tubulin 85D-OE* | Yes | | 0.0037 | 14, 19 |
| **Fig 12M 1st order** | | **Two-way ANOVA with Tukey's multiple comparison test** | | |
| Control | Yes | | | 10 |
| Control *vs* *mts-IR;β-tubulin 85D-OE* | Yes | | 0.0060 | 10, 19 |
| *mts-IR vs mts-IR;β-tubulin 85D-OE* | Yes | | <0.0001 | 14, 19 |
| **Fig 12M 2nd order** | | **Two-way ANOVA with Tukey's multiple comparison test** | | |
| Control | Yes | | | 10 |
| Control *vs* *mts-IR;β-tubulin 85D-OE* | Yes | | 0.0213 | 10, 19 |
| *mts-IR vs mts-IR;β-tubulin 85D-OE* | Yes | | 0.0046 | 14, 19 |
| **Fig 12M 3rd order** | | **Two-way ANOVA with Tukey's multiple comparison test** | | |
| Control | | | | 10 |
| Control *vs* *mts-IR;β-tubulin 85D-OE* | Yes | | 0.9899 | 10, 19 |
| *mts-IR vs mts-IR;β-tubulin 85D-OE* | Yes | | 0.0181 | 14, 19 |
| **Fig 12M 4th order** | | **Two-way ANOVA with Tukey's multiple comparison test** | | |
| Control | | | | 10 |
| Control *vs* *mts-IR;β-tubulin 85D-OE* | Yes | | 0.9658 | 10, 19 |
| *mts-IR vs mts-IR;β-tubulin 85D-OE* | Yes | | 0.6491 | 14, 19 |

### SUPPLEMENTALS

| | | | | |
|---|---|---|---|---|
| **Fig S1B** | | **Kruskal-Wallis with Dunn's multiple comparison test** | | |
| Control | Yes | | | 10 |
| *mts-IR* | No | | <0.0001 | 12 |

| | | | | |
|---|---|---|---|---|
| *PP2A-29B-IR* | Yes | | 0.0002 | 12 |
| Control-MARCM | Yes | | | 15 |
| $mts^{k12502}$ | Yes | | <0.0001 | 12 |
| **Fig S1C** | | **Kruskal-Wallis with Dunn's multiple comparison test** | | |
| Control | Yes | | | 10 |
| *mts-IR* | No | | 0.1769 | 12 |
| *PP2A-29B-IR* | Yes | | 0.0018 | 12 |
| Control-MARCM | Yes | | | 15 |
| $mts^{k12502}$ | Yes | | <0.0001 | 12 |
| **Fig S1I** | | **One-way ANOVA with Sidak's multiple comparison test** | | |
| Control | Yes | | | 10 |
| *wdb-IR* | Yes | | 0.0082 | 13 |
| *wrd-IR* | Yes | | 0.0212 | 12 |
| *tws-IR* | Yes | | 0.0192 | 16 |
| *CG4733-IR* | Yes | | 0.1024 | 12 |
| **Fig S1J** | | **Kruskal-Wallis with Dunn's multiple comparison test** | | |
| Control | Yes | | | 10 |
| *wdb-IR* | Yes | | 0.0002 | 13 |
| *wrd-IR* | No | | 0.4980 | 12 |
| *tws-IR* | Yes | | 0.0522 | 16 |
| *CG4733-IR* | Yes | | 0.6844 | 12 |
| **Fig S1O** | | **Kruskal-Wallis with Dunn's multiple comparison test** | | |
| Control | Yes | | | 10 |
| *wrd-IR* | Yes | | 0.9407 | 11 |
| *tws-IR* | No | | >0.9999 | 10 |
| *CG4733-IR* | No | | 0.0278 | 16 |
| **Fig S1P** | | **Kruskal-Wallis with Dunn's multiple comparison test** | | |
| Control | Yes | | | 10 |
| *wrd-IR* | Yes | | >0.9999 | 11 |
| *tws-IR* | Yes | | >0.9999 | 10 |
| *CG4733-IR* | No | | 0.4583 | 16 |
| **Fig S2B CI** | | **Two-way ANOVA with Dunnett's multiple comparison test** | | |
| Control | | | | 14 |

| | | | | |
|---|---|---|---|---|
| *mts-IR* | | | <0.0001 | 18 |
| **Fig S2B CIII** | | **Two-way ANOVA with Dunnett's multiple comparison test** | | |
| Control | | | | 14 |
| *mts-IR* | | | <0.0001 | 18 |
| **Fig S2B CIV** | | **Two-way ANOVA with Dunnett's multiple comparison test** | | |
| Control | | | | 14 |
| *mts-IR* | | | 0.0007 | 18 |
| **Fig S2E** | | **Unpaired t-test** | | |
| Control | | | | 11 |
| *mts-IR* | | | 0.0427 | 16 |
| **Fig S3C CI Anterograde** | | **Two-way ANOVA with Sidak's multiple comparison test** | | |
| Control | | | | 53 (comets) |
| *mts-IR* | | | 0.6881 | 35 (comets) |
| **Fig S3C CI Retrograde** | | **Two-way ANOVA with Sidak's multiple comparison test** | | |
| Control | | | | 53 (comets) |
| *mts-IR* | | | 0.6881 | 35 (comets) |
| **Fig S3D CIV Anterograde** | | **Two-way ANOVA with Sidak's multiple comparison test** | | |
| Control | | | | 38 (comets) |
| *mts-IR* | | | 0.9237 | 28 (comets) |
| **Fig S3D CIV Retrograde** | | **Two-way ANOVA with Sidak's multiple comparison test** | | |
| Control | | | | 38 (comets) |
| *mts-IR* | | | 0.9237 | 28 (comets) |
| **Fig S3I CI Anterograde** | | **Two-way ANOVA with Sidak's multiple comparison test** | | |
| Control | | | | 49 (comets) |
| *mts-IR* | | | 0.9237 | 129 (comets) |
| **Fig S3I CI Retrograde** | | **Two-way ANOVA with Sidak's multiple comparison test** | | |
| Control | | | | 49 (comets) |
| *mts-IR* | | | 0.9237 | 129 (comets) |
| **Fig S3J** | | **Mann-Whitney test** | | |

| | | | | |
|---|---|---|---|---|
| Control | Yes | | | 12 |
| *mts-IR* | No | | <0.0001 | 8 |
| **Fig S4E** | | **Mann-Whitney test** | | |
| Control | No | | | 12 |
| *mts-IR* | No | | 0.0070 | 15 |
| **Fig S4F** | | **Mann-Whitney test** | | |
| Control | No | | | 12 |
| *mts-IR* | Yes | | 0.0087 | 15 |
| **Fig S5F** | | **One-way ANOVA with Sidak's multiple comparison test** | | |
| Control | Yes | | | 11 |
| *β-tubulin 56D-IR* | Yes | | <0.0001 | 10 |
| *β-tubulin 60D-IR* | Yes | | <0.0001 | 10 |
| *β-tubulin 97EF-IR* | Yes | | 0.2713 | 16 |
| *CG32396-IR* | Yes | | 0.0065 | 9 |
| **Fig S5G** | | **One-way ANOVA with Dunnett's multiple comparison test** | | |
| Control | Yes | | | |
| *β-tubulin 56D-IR* | Yes | | <0.0001 | 10 |
| *β-tubulin 60D-IR* | Yes | | 0.8552 | 10 |
| *β-tubulin 97EF-IR* | Yes | | 0.2081 | 16 |
| *CG32396-IR* | Yes | | >0.9999 | 9 |
| **Fig S6E** | | **One-way ANOVA with Dunnett's multiple comparison test** | | |
| Control | Yes | | | 11 |
| *β-tubulin_S172A_T219A* | Yes | | 0.2192 | 11 |
| *β-tubulin_S172E_T219E* | Yes | | 0.0015 | 7 |
| **Fig S6F** | | **One-way ANOVA with Dunnett's multiple comparison test** | | |
| Control | Yes | | | 11 |
| *β-tubulin_S172A_T219A* | Yes | | 0.0809 | 11 |
| *β-tubulin_S172E_T219E* | Yes | | 0.0010 | 7 |
