## Supplementary material for "PP2A phosphatase regulates cell-type specific cytoskeletal organization to drive dendritic diversification": Table S3

### LIST OF GENETIC STRAINS USED IN THIS STUDY

| ***NO.*** | Gene Symbol | Stock number/Source |
|---|---|---|
| **1.** | *mts-IR* | **B57034** |
| **2.** | *mts-IR* | B27723 |
| **3.** | *mts-IR* | B38337 |
| **4.** | *PP2A-29B-IR* | **B50533** |
| **5.** | *PP2A-29B-IR* | B29384 |
| **6.** | *PP2A-29B-IR* | B43283 |
| **7.** | *PP2A-29B-IR* | v49672 |
| **8.** | *wdb-IR* | **B38950** |
| **9.** | *wdb-IR* | B38901 |
| **10.** | *wdb-IR* | v27470 |
| **11.** | *wdb-IR* | v101406 |
| **12.** | *wrd-IR* | **B38900** |
| **13.** | *wrd-IR* | B30512 |
| **14.** | *wrd-IR* | v22614 |
| **15.** | *wrd-IR* | v22615 |
| **16.** | *tws-IR* | **B58053** |
| **17.** | *tws-IR* | B28714 |
| **18.** | *tws-IR* | B36698 |
| **19.** | *tws-IR* | v34340 |
| **20.** | *tws-IR* | v104167 |
| **21.** | *CG4733-IR* | **v34894** |
| **22.** | *CG4733-IR* | v107621 |
| **23.** | *cka-IR* | **v35234** |
| **24.** | *cka-IR* | v35232 |
| **25.** | *β-tubulin85D-IR* | **B65163** |
| **25.** | *β-tubulin56D-IR* | **B65028** |
| **26.** | *β-tubulin60D-IR* | **B65856** |
| **27.** | *β-tubulin97EF-IR* | **B64858** |
| **28.** | *CG32396-IR* | **B68474** |
| **29.** | *UAS-mts* | **B53709** |
| | ***Overexpression lines*** | |
| **30.** | *UAS-PP2A-29B* | **B55049** |
| **31.** | *UAS-PP2A-29B* | B55048 |
| **32.** | *UAS-PP2A-29B* | B55050 |
| **33.** | *UAS-wdb* | **B55052** |
| **34.** | *UAS-wdb* | B55051 |
| **35.** | *UAS-cka-eGFP* | **B53757** |
| **36.** | *UAS-cka-eGFP* | B53786 |
| **37.** | *UAS-foxo* | **B9575** |
| **38.** | *UAS-foxo* | **B42221** |
| **39.** | *UAS-β-tubulin85D* | **F001711** |
| | ***Mutant lines*** | |
| **40.** | $mts^{k12502}$ | **111466 Kyoto Stock Center** |
| **41.** | $wdb^{14}$ | **B53712** |
| **42.** | $UAS\text{-}BFP\text{-}cka^{\Delta PP2A}$ | (Neisch *et al*, 2017) |
| | ***Additional fly lines used*** | |
| **43.** | $GAL4^{477}$*,UASmCD8::GFP/CyO,tubP-GAL80;*$GAL4^{ppk.1.9}$*,UAS-mCD8::GFP (CIV-GAL4)* | |
| **44.** | $GAL4^{221}$*, UAS-mCD8::GFP (CI-GAL4)* | |

| | | |
|---|---|---|
| **45.** | *$GAL^{5\text{-}40}$UAS-Venus:pm SOP-FLP#42;tubP-GAL80FRT40A (2L MARCM)* | (Shimono *et al*, 2014) |
| **46.** | *hsFLP-UASmCD8:GFP; $GAL4^{109(2)80}$UAS-mCD8::GFP SOP-$FLP^{\#73}$/CyO; FRT82B tub-GAL80 (3R MARCM)* | (Shimono *et al*, 2014) |
| **47.** | *$GAL4^{477}$;ppk-hCD4::tdTomato, ppk-CD8-eGFP* | |
| **48.** | *$GAL4^{Nanos}$;+;ppk-::tdTomato* | **B7303** |
| **49.** | *$GAL4^{477}$,UAS-mCD8::GFP;UAS-Ct-IR/TM3,Ser* | |
| **50.** | *UAS-GMA;$GAL4^{477}$,UAS-mCherry::JUPITER* | (Das *et al*, 2017) |
| **51.** | *UAS-GMA;+:$GAL4^{221}$,UAS-mCherry::JUPITER* | (Das *et al*, 2017) |
| **52.** | *UAS-alphaTUB84B.tdEOS* | **B51314** |
| **53.** | *UAS-LifeAct.tdEOS* | This study |
| **54.** | *$GAL4^{ppk}$/CyO;ppk-EB1::GFP* | **Gift from Dr. Jill Wildonger** |
| **55.** | *UAS-EB1::GFP* | **B35512** |
| **56.** | *UAS-γ-tubulin23C-GFP* | **Gift from Dr. Melissa Rolls** |
| **57.** | *UAS-YFP-Patronin* | (Feng *et al*, 2019) |
| **58.** | *UAS-mitoGFP.AP* | **B8442** |
| **59.** | *$GAL4^{477}$,UAS-MANII-eGFP;ppk-hCD4-tdTOMATO* | |
| **60.** | *UAS-MANII-eGFP;$GAL4^{221}$,UAS-mCD8::RFP* | |
| **61.** | *UAS-CD4-tdTom* | **B35837** |
| **62.** | *UAS-Luc-IR* | **B31603** |
| **63.** | *UAS-DenMark, UAS-syt.eGFP* | **B33064** |
| **64.** | *UAS-βtubulin85D_S172A_219A* | This study |
| **65.** | *UAS-βtubulin85D_S172E_219E* | This study |
| **66.** | *OregonR(ORR) (control strain)* | |

Stock numbers refers to either Bloomington Stock Center or if it starts with 'v', it refers to Vienna Drosophila Research Centre; F refers to FlyORF. Stock numbers of the genes for which representative data are presented in this paper are highlighted in **bold.**


Das R, Bhattacharjee S, Patel AA, Harris JM, Bhattacharya S, Letcher JM, Clark SG, Nanda S, Iyer EPR, Ascoli GA, *et al* (2017) Dendritic Cytoskeletal Architecture Is Modulated by Combinatorial Transcriptional Regulation in Drosophila melanogaster. *Genetics* 207: genetics.300393.2017

Feng C, Thyagarajan P, Shorey M, Seebold DY, Weiner AT, Albertson RM, Rao KS, Sagasti A, Goetschius DJ & Rolls MM (2019) Patronin-mediated minus end growth is required for dendritic microtubule polarity. *J Cell Biol* 218: 2309–2328

Neisch AL, Neufeld TP & Hays TS (2017) A STRIPAK complex mediates axonal transport of autophagosomes and dense core vesicles through PP2A regulation. *J Cell Biol* 216: 441–461

Shimono K, Fujishima K, Nomura T, Ohashi M, Usui T, Kengaku M, Toyoda A & Uemura T (2014) An evolutionarily conserved protein CHORD regulates scaling of dendritic arbors with body size. *Sci Rep* 4: 1–8
